# Consensus native-like hepatitis C virus E1E2 engages broadly neutralizing antibody precursors

**DOI:** 10.64898/2026.08.25.746952

**Authors:** Fabian Mulder, Fabien Cannac, Joan Capella-Pujol, Stan Peters, Meliawati Poniman, Wouter Olijhoek, Luke Granger, Marco Briones-Orta, Kostas Paschos, Sophie van der Pol, Ruben Walen, Maddy L. Newby, Wen-Hsin Lee, Laura Radić, Ian Zon, Timm Weber, Max Crispin, Florian Klein, Robin J. Shattock, Rogier W. Sanders, Andrew B. Ward, Janke Schinkel, Kwinten Sliepen

## Abstract

A major goal for hepatitis C virus (HCV) vaccine development is to elicit broadly neutralizing antibodies (bNAbs) against the E1E2 glycoprotein complex located on the viral surface. Inducing HCV bNAbs requires engagement of their germline B cell precursors. HCV glycoproteins usually do not bind and activate inferred germline precursors of bNAbs (igl-bNAbs), possibly because most circulating strains contain non-conserved isolate-specific residues, even in bNAb epitopes. Here, we generated stabilized native-like soluble E1E2 (sE1E2) antigens based on a consensus sequence of HCV (HepCon) to limit the exposure of antigenically rare residues. The antigenicity and glycosylation profiles show that HepCon sE1E2 resembles a native-like E1E2 heterodimer. HepCon sE1E2 induced cross-reactive neutralizing antibody responses as a soluble protein immunogen and as membrane-anchored mRNA-delivered immunogen in animals. Importantly, HepCon sE1E2 engages multiple igl-bNAbs against two major epitopes: antigenic region 3 (AR3), which is targeted by igl-bNAbs derived from the widely expressed human *VH1-69* B cell gene, and antigenic region 4 (AR4), which is only present on native-like E1E2. Nanoparticles with HepCon sE1E2 efficiently activated B cell lines expressing AR3 and AR4 igl-bNAb B cell receptors *in vitro*. Finally, using HepCon sE1E2 we elucidated the atomic contacts of an AR3 igl-bNAb by cryo-electron microscopy. Thus, HepCon sE1E2 is a promising candidate for germline-targeting vaccination strategies.

## Introduction

Hepatitis C virus (HCV) is a bloodborne virus that causes a slowly progressing chronic liver infection that can result in liver cirrhosis and/or cancer. Highly effective direct acting antiviral agents became available in 2014, which prompted the WHO in 2016 to aim for HCV elimination by 2030.^1^ However, HCV infection still afflicts approximately 47 million people and caused an estimated 239,000 HCV related deaths in 2024.^2^ Approximately 0.9 million new infections occur annually and liver-related end-stage outcomes are expected to increase over the coming years.^2,3^ It is therefore now clear that an effective preventive vaccine will be critical for global HCV elimination.

The only target for neutralizing antibodies (NAbs) is the E1E2 glycoprotein complex, which mediates cell entry by binding to scavenger receptor class B type 1 and tetraspanin CD81 on the host cell surface.^4–10^ E1E2 is a conformationally dynamic protein complex^11,12^ covered by an extensive glycan shield, and is extremely sequence diverse, which makes it difficult to induce potent cross-reactive NAbs by vaccination.^13,14^ However, some infected individuals develop broadly neutralizing antibodies (bNAbs)^15–23^ which neutralize a wide range of HCV isolates by targeting conserved epitopes on E1E2 and provide protection against chronic HCV (re-)infection in chimeric liver mice and humans.^17,18,24–32^ These studies provide strong impetus to develop antigens and vaccine strategies aimed at eliciting bNAbs.

During natural infection HCV bNAbs emerge after an evolutionary process initiated by recognition of E1E2 by the B cell receptor (BCR) on rare naïve B cells.^29^ The activated precursor B cells undergo rounds of affinity maturation to develop into bNAb-producing B cells.^33^ Germline-targeting vaccination strategies utilize specifically designed priming immunogens to activate these rare naive B cells, followed by boosting immunogens to guide the bNAb maturation process. Proof-of-concept vaccination studies in the HIV-1 field demonstrated that specific engagement of precursor bNAb B cells in humans is feasible and these studies are considered to be important steps towards a vaccine that induces HIV-1 bNAbs.^34–37^ Important tools for studying and designing germline-targeting vaccines are the inferred germline precursors of bNAbs (igl-bNAbs).^38^

The germline-targeting concept may also hold promise against other sequence-diverse viruses, including HCV. However, current E1E2-based HCV immunogens are not designed to and generally do not bind to HCV igl-bNAbs and are thus unlikely to consistently induce HCV bNAbs in humans. Furthermore, as the presence of NAbs against multiple epitopes is associated with enhanced neutralization breadth, limited viral escape and enhanced HCV clearance, an HCV vaccine candidate should probably engage igl-bNAbs directed against multiple epitopes.^30,39–42^

One major target for HCV bNAbs is antigenic region (AR) 3 which overlaps with the binding site for CD81. As a consequence AR3 bNAbs neutralize by inhibiting the interaction of E1E2 with CD81.^15,18^ AR3 bNAbs have been isolated from several individuals, including individuals that cleared their HCV infection, implying that these bNAbs contributed to clearance. Furthermore, AR3 bNAbs appear to provide protection against reinfection.^17,25^ Virtually all AR3 bNAbs originate from the *VH1-69* B cell gene family, which is abundantly expressed in all healthy humans through multiple *VH1-69* alleles.^18,29,43–45^ Compared to HIV-1 bNAbs, *VH1-69*-derived AR3-directed HCV bNAbs require little somatic hypermutation.^29^ Most of these bNAbs contain an elongated CDRH3 loop and/or an inter-CDRH3-loop disulfide bond, but these are not absolutely required to achieve neutralizing potency and breadth.^19,44,46,47^ Because of these features, it should be feasible to induce AR3 bNAbs, provided that the desired germline precursor *VH1-69* B cells are engaged.

AR4 is a metastable epitope on E2 that is only present on viral E1E2 or on native-like soluble E1E2 (sE1E2) and is recognized by a number of potent bNAbs.^12,15,17,20,48–50^ However, AR4 bNAbs have largely eluded structural characterization at high resolution, except for the AR4A bNAb.^12,15^ Therefore, they have not been studied as extensively as AR3 bNAbs and no shared genetic and/or structural signatures have been discovered.^12,15,17,20,51^ In any case, for a multi-epitope germline-targeting strategy to be successful, it is paramount to target precursor bNAb B cells against the dominant epitopes in AR3 and AR4.

One strategy to overcome HCV’s diversity is to generate an E1E2 immunogen based on a consensus sequence, which should be most closely related to all sequences in a population.^52^ Because consensus sequences by definition contain the most common residue at each position, a consensus E1E2 immunogen should induce fewer isolate-specific antibody (Ab) responses, which might favor recognition by bNAbs and their inferred germline precursors.^52^ However, without applying methods for stabilization, consensus-based viral glycoprotein sequences do not generally assume a native-like conformation compatible with presentation of the AR4 epitopes.^52^

Here, we applied structure-based design templates to generate native-like HCV sE1E2 antigens based on a consensus sequence (HepCon).^50^ HepCon sE1E2 induced cross-NAb responses in rabbits and mice when delivered as soluble proteins or through mRNA. Remarkably, HepCon sE1E2 engaged a wide range of igl-bNAbs against AR3 and AR4. Finally, we solved the cryo-electron microscopy (cryo-EM) structure of HepCon sE1E2 in complex with an igl-bNAb targeting AR3, igl-1416_01_E03. These studies identify HepCon sE1E2 as a promising HCV vaccine candidate, in particular in the context of germline-targeting vaccination strategies.

## Results

### Screening and selection of recombinant consensus sE1E2 antigens

To generate an E1E2 consensus sequence we first obtained ten separate consensus E1E2 sequences of subtypes 1a, 1b, 1c, 2a, 2b, 3a, 4a, 5a, 6a and 6k, covering the six major genotypes, from the Los Alamos HCV sequence database.^53^ Next, we generated a single consensus sequence (HepCon) of these ten consensus sequences and reference strain H77 (Genbank AF009606) (Supplementary Fig. 1A). In a maximum likelihood tree of over 1,000 E1E2 sequences, the subtype-specific consensus sequences were located near the branch bases of each subtype, while HepCon was located near the base of the tree, indicating that HepCon has a favorable sequence similarity to all circulating strains (Fig. 1A).^53^ We applied the sE1E2.v3-LZ design to the eleven consensus sequences to produce consensus sE1E2 antigens (Fig. 1B).^50^ In short, in sE1E2.v3-LZ the E1 and E2 ectodomains were fused to a Jun-Fos leucine zipper (LZ)^48^, separated by a furin site and stabilized by a short hydrophilic linker in E1 plus the 326P, 622A^54^ and 682P substitutions (Supplementary Fig. 1B, 2A).

**Figure 1:**
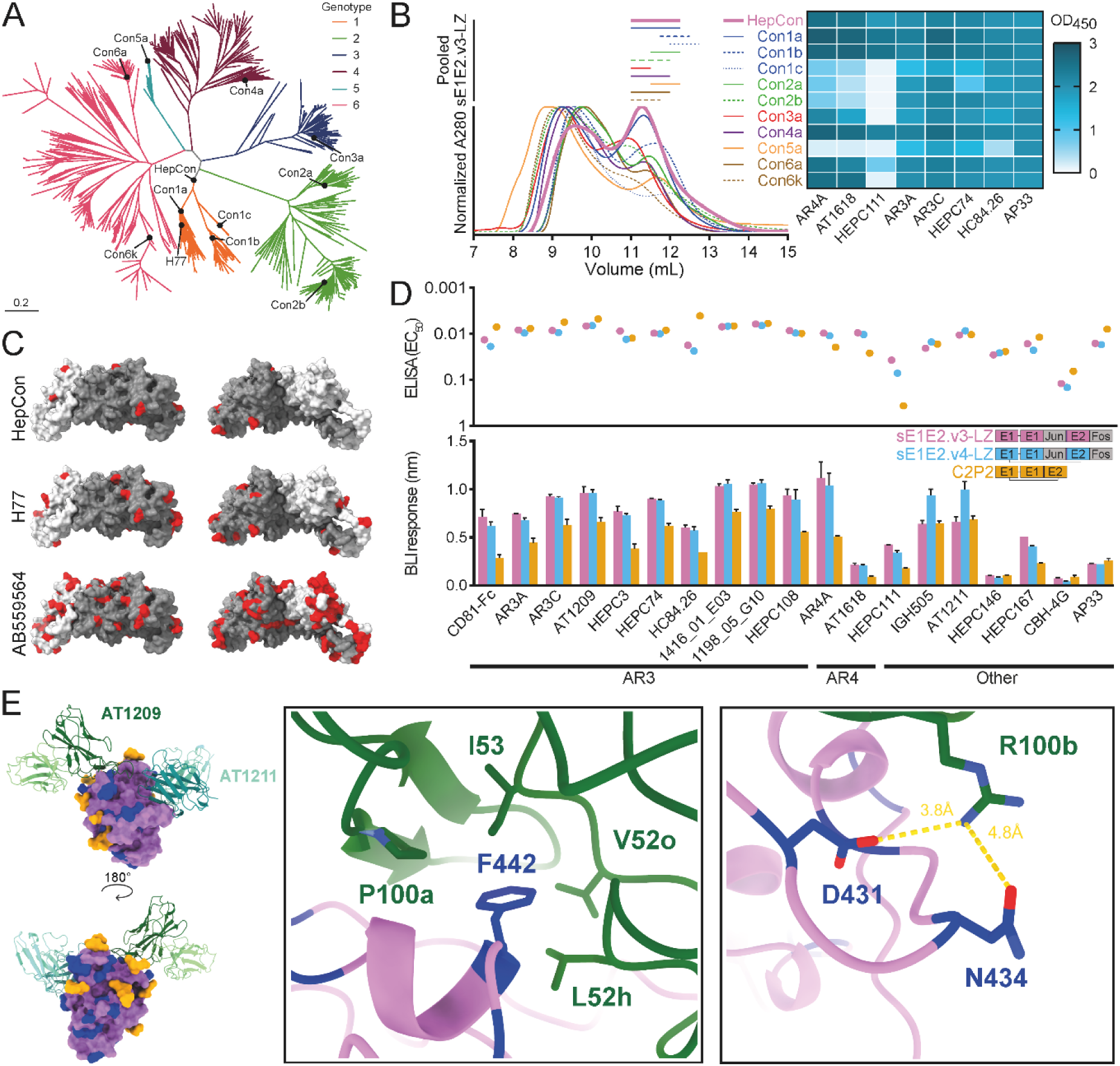
Design, screening and biophysical characterization of consensus-based sE1E2.v3-LZ, sE1E2.v4-LZ and C2P2. **A** Maximum likelihood tree generated using IQTREE version 2 of 1146 concatenated E1E2 protein sequences, using the LG substitution model with four discrete Gamma rate categories, visualized with ggtree using ‘daylight’ layout.^53,65,66^ The 10 different subtype consensuses, H77 and HepCon are indicated as well as the genetic distance bar (substitutions per sequence position). **B** Normalized SEC profiles of Con1a, −1b, −1c, −2a, −2b, −3a, −4a, −5a, −6a, −6k and HepCon sE1E2.v3-LZ expressed in HEK293F cells on a Superdex 200 Increase 10/300 GL column. Pooled sE1E2.v3-LZ heterodimer fractions are indicated for each construct. Additionally, ELISA screening is shown for the 10 subtype consensus and HepCon sE1E2.v3-LZ. OD_450_ was measured at Ab concentrations of 5 µg/mL. **C** E1E2 structure (PDB: 7T6X) with E1 colored light grey, E2 colored dark grey and uncommon (<25% present at each position amongst the 15,330 isolates^53^) residues colored red based on the corresponding residues from HepCon (top), H77 (middle) and the randomly selected genotype 2b sequence AB559564 (bottom).^67^ **D** Antigenicity profiling. Top: ELISA. The antigens were immobilized on Strep-TactinXT ELISA plates and probed using a panel of 18 mAbs and CD81-Fc. The EC_50_-values are derived from sigmoidal binding curves (Supplementary Fig. 5A). Bottom: BLI. The same panel of mAbs and CD81-Fc were immobilized on protein A sensors followed by 300 second incubation of 250 nM HepCon antigens and subsequent 300 second incubation of running buffer. The maximum binding signal was obtained from the full BLI curve and corrected for the mock (Supplementary Fig. 5B). The means with standard deviation (SD) are plotted. All ELISAs and BLI assays were performed in duplo. **E** (left) Cryo-EM model of AT1209 and AT1211 bNAb in complex with HepCon sE1E2.v4-LZ. Both bNAbs are represented as ribbon, with AT1211 heavy and light chains colored teal and light blue and AT1209 heavy and light chains colored dark and light green, respectively. E2 is represented as a violet surface model, with modelled glycosylations in orange, and positions of the different mutations inserted on E2 in dark blue. (right) Main epitope / paratope interactions between AT1209 and E2, following the same color code as above. Interacting residues shown as sticks.

We expressed these constructs in HEK293F cells and used Strep-TactinXT for purification. The yields ranged from 2.4 mg/L (Con1c) to 8.3 mg/L (Con6k). Size-exclusion chromatography (SEC) revealed varying amounts of heterodimers compared to dimers of heterodimers and higher order molecular weight species, with HepCon resulting in the highest proportion of heterodimers (51%) and Con1c the lowest (21%) (Fig. 1B, Supplementary Fig. 2B). Next, we screened the purified sE1E2 heterodimers for binding to conformational AR4-targeting bNAbs AR4A and AT1618, and other bNAbs, in Enzyme-Linked Immunosorbent Assay (ELISA) (Fig. 1B). The Con1a, 1b, 3a, 4a, 6a, 6k and HepCon sE1E2.v3-LZ proteins showed strong binding to AR4A and AT1618 (OD_450_ > 1.5) indicating native-like folding, while others showed weaker (OD_450_ < 1.0; Con1c, 2a and 2b) or no (OD_450_ < 0.2; Con5a) binding to AR4A and AT1618. Overall, seven out of eleven consensus sE1E2.v3-LZ proteins displayed favorable native-like antigenicity.

### HepCon E1E2 displays few uncommon antigenic determinants

We assessed to what extend the eleven consensus E1E2s contained rare amino acids compared to 15,330 isolate sequences using a genotype weighting strategy.^53^ As expected, HepCon exhibited fewer uncommon amino acids (<25% present at each position) than the subtype consensuses (Supplementary Fig. 3A) or circulating isolates (Supplementary Fig. 3B). We visualized the uncommon amino acids on the surface of the E1E2 heterodimer for HepCon E1E2, the reference strain H77 E1E2 and a randomly selected genotype 2b E1E2 (strain AB559564) (Fig. 1C). HepCon E1E2 contained 12 uncommon amino acids, of which two (T444 (20%) and S522 (16%)) are near AR3. Note that residues 444 and 522 are inherently diverse with the highest positional frequency for Y444 (33%) and K522 (22%). H77 contained 26 uncommon residues, of which five (Q444 (0.7%), S522 (16%), S528 (15%), A531 (16%) and D533 (21%)) are located near or in AR3. The genotype 2b AB559564 E1E2 sequence contained 112 uncommon amino acids, of which six near or in AR3 and five in AR4. Based on the rarity analyses, proportional heterodimer yields and antigenic screening, we decided to focus our efforts on optimizing the HepCon E1E2 construct.

### HepCon optimization yields stable and soluble native-like sE1E2 heterodimers

To increase the antigenic stability, we added an interprotomer disulfide bond between residues 244 in E1 and 687 in E2 to generate HepCon sE1E2.v4-LZ. Subsequently, we removed the immunogenic LZ dimerization domains to generate HepCon C2P2, as described previously^50^ (Supplementary Fig. 2A) We produced and purified HepCon sE1E2.v3-LZ, sE1E2.v4-LZ and C2P2 as described above, which resulted in protein yields of 4.8, 6.2 and 8.2 mg/L, respectively. All three constructs produced a mixture of heterodimers and dimers of heterodimers (Supplementary Fig. 4A). Reducing Sodium Dodecyl Sulfate Polyacrylamide Gel Electrophoresis (SDS-PAGE) on the heterodimeric fractions confirmed complete furin cleavage (Supplementary Fig. 4B) and non-reducing SDS-PAGE revealed a single band for sE1E2.v4-LZ and C2P2 indicating the formation of the designed C244-C687 disulfide bond. An E1-Jun-E2-Fos band was also visible for sE1E2.v3-LZ, which points at the presence of an unintended covalent linkage likely due to disulfide scrambling.^12,55–59^

Next, we compared their antigenicity using ELISA (Supplementary Fig. 5A) with the half maximal effective concentrations (EC_50_) summarized in Figure 1D, top. The proteins engaged all 18 tested monoclonal Abs (mAbs), including those against the metastable AR4 epitope (AR4A (EC_50_: 0.010 - 0.019 μg/mL) and AT1618 (EC_50_: 0.009 - 0.026 μg/mL)), indicative of native-like folding.^12,15,50^ Efficient binding by mAbs to other epitopes and binding by CD81-Fc implies overall native-like conformation and antigenicity. In biolayer interferometry (BLI) we detected binding to the same mAbs as in ELISA (Fig. 1D, bottom, Supplementary Fig. 5B) and binding values in both assays correlated well (Supplementary Fig. 6A). The smaller size of C2P2 might partially explain the lower maximum binding signal in BLI. HepCon C2P2 displayed slightly weaker binding to conformational bNAbs AR4A, AT1618 and HEPC111 specifically, indicating that the LZ domains act as folding chaperones and/or enhance heterodimeric stability.^50^ But in general, the design template did not perturb the overall antigenicity profile since binding values in both ELISA and BLI correlated well between the three HepCon constructs (Supplementary Fig. 6B).

### HepCon sE1E2 heterodimers are extensively glycosylated

*N*-linked glycosylation is crucial for E1E2 folding, antigenicity and stability.^13^ Most E1E2 sequences contain between ∼14-17 highly conserved potential *N*-linked glycosylation sites (PNGS).^12,60^ To assess whether the non-natural HepCon sequence contains a native-like glycan shield, we performed site-specific glycan analysis (Supplementary Fig. 6C).^61^ We found that all 16 PNGS were glycosylated to some degree. Of note, N325 contains a highly conserved proline directly downstream of its PNGS sequon (NxT or NxS) and is therefore never glycosylated.^12,62^ Most sites contained processed complex glycans, except for N645 and N305, which contained higher proportions of oligomannose glycans. For some PNGS occupancy differed substantially, most notably for the noncanonical NXV motif of N695, which was more occupied on sE1E2.v3-LZ (°60%) than sE1E2.v4-LZ (°20%) and C2P2 (°5%).^12^ Other notable differences include the higher glycan occupancy at N209 and N305 for sE1E2.v3-LZ and slightly lower N417 occupancy for sE1E2.v4-LZ. The decreased glycan occupancy at N430 and N695 in C2P2 might partly explain its decreased binding to some bNAbs, as these glycans surround AR3 and AR4.^12,57,63^ Overall, glycosylation of HepCon sE1E2 is similar to that of sE1E2 based on natural sequences.^50^

### Cryo-EM structure of HepCon sE1E2.v4-LZ in complex with AT1209 and AT1211

Thus far, structures of E2 and E1E2 proteins based on naturally occurring primary HCV strains have been solved.^12,46,57,60,64^ To determine whether HepCon sE1E2 is structurally similar to these proteins and to inform future structure-based vaccine design, we determined the structure of HepCon sE1E2.v4-LZ in complex with two VH1-69 bNAbs, AT1209 targeting AR3 and AT1211 targeting domain C, at 3.3 Å by cryo-EM (Fig.1E, Supplementary Fig. 7A-C). We previously reported the structure of AMS0232 sE1E2.v4-LZ in complex with AT1211, and the structure of HepCon sE1E2.v4-LZ is highly similar to that previous structure, with an RMSD of 0.86 Å for the antigen and 0.75 Å for the AT1211 antibody.^50^ We also observed glycosylation at multiple PNGSs and were able to build glycans at positions N423, N430, N448, N532, N540, N556, N623 and N645 (Supplementary Fig. 7D). Furthermore, the presence of AT1209 stabilized some residues in the N-terminal region of E2. This includes PNGSs and the antigenic region between amino acids 430 and 450 on E2, residues that were not resolved in the previous sE1E2.v4-LZ structure (Supplementary Fig. 7C).

We also previously reported structures of the full-length, unmodified E1E2 in complex with several Abs including AT1209 (PDB: 7T6X).^12^ While the epitope – paratope interface between both complexes is conserved (RMSD: 0.79 Å between both models for E2 and AT1209), the HepCon sE1E2 contains a phenylalanine at residue 442 instead of isoleucine in AMS0232 E1E2. F442 forms many interactions with hydrophobic residues on the atypically long CDRH2 of AT1209 (Fig. 1E) including CH-π interactions with the nearby methyl groups of L52h and I53, probably strengthening the interface. Substitutions T431D and E434N in HepCon compared to AMS0232 also stabilize binding to AT1209 by interacting with R100b of the CDRH3. As previously reported, the long AT1209 CDRH2 has a hydrophobic motif rich in glycines and leucines that wedges into a hydrophobic pocket of E2 (buried surface area (B.S.A): 475 Å^2^), but remarkably, also forms heterotypic contacts with V53 and Y97 from the heavy chain (HC) of AT1211 (B.S.A: 112 Å^2^, Supplementary Fig. 7E).

While previous structures of E2 in complex with AT1209 and AT1211 separately delineated their individual epitopes, the new structure with both Abs reveal how these Abs can bind simultaneously without impeding each other, but instead interact with each other. We note that AT1211 and AT1209 were isolated from the same individual.^16^ That fact and the structural observations made here suggest that these antibody lineages have co-evolved to accommodate simultaneous binding.

Overall, the structural characteristics of the resolved region of HepCon E1E2 are similar to those of E1E2 and E2 structures from circulating strains (PDBs: 7T6X^12^, 8FSJ^60^ and 9Z5U^50^). Furthermore, the structure reveals that HepCon E1E2 appropriately presents the important AR3.

### HepCon sE1E2.v4-LZ induces cross-reactive antibodies

Next, we assessed immunogenicity of HepCon sE1E2.v4-LZ, AMS0232 sE1E2.v4-LZ (genotype 1a, data previously reported^50^) and a tetravalent cocktail of AMS0232, H77 (genotype 1a), UKNP4.1.1 (genotype 4a) and AMS3a (genotype 3a) sE1E2.v4-LZ (Fig. 2A). Rabbits were immunized three times (weeks 0, 4 and 20) with 18 μg total protein per immunization in squalene emulsion adjuvant.

**Figure 2:**
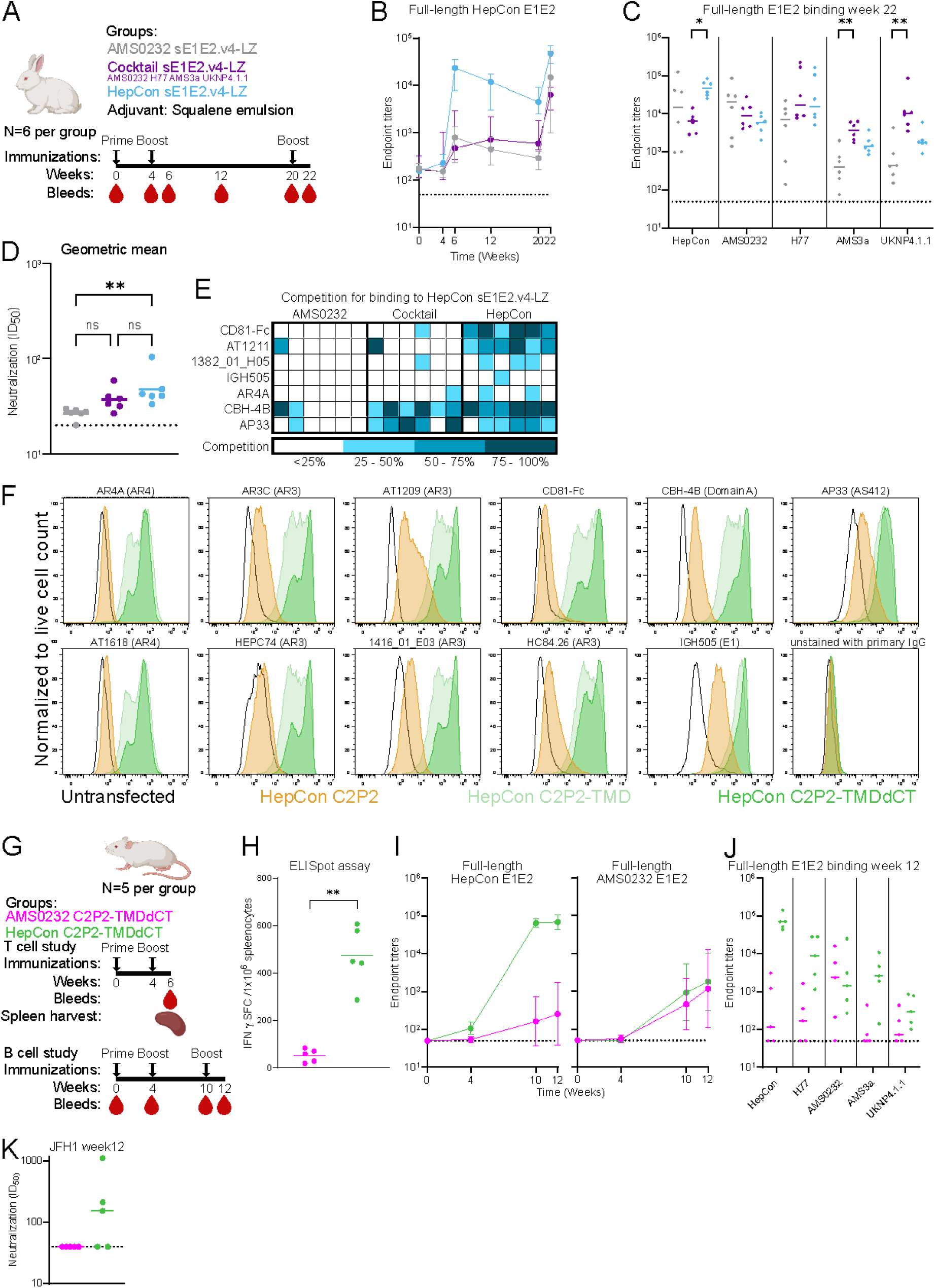
Immunogenicity of recombinant sE1E2 and mRNA vaccines. **A** Groups of six New Zealand White rabbits were immunized at weeks 0, 4 and 20 with 18 µg of either AMS0232 sE1E2.v4-LZ, HepCon sE1E2.v4-LZ or a cocktail of AMS0232, H77, UKNP4.1.1 and AMS3a sE1E2.v4-LZ immunogens. Bleeds were taken at weeks 0, 4, 6, 12, 20 and 22. **B,C** Rabbit serum ELISA endpoint binding titers to full-length HepCon E1E2 for all six timepoints (**B**) and to full-length HepCon, AMS0232, H77, UKNP4.1.1 and AMS3a E1E2 for week 22 (**C**). AMS0232 binding data has been reported previously.^50^ Data in **B** reported as median with interquartile range and in **C** the median is indicated. **D** Global geometric mean titers (GGMT) for neutralization of 13 HCVpp strains at week 22. GGMT were calculated based on the neutralization ID_50_-values against 13 HCVpp strains and each dot represents a single rabbit serum. The geometric means are indicated. Neutralization ID_50_-values are depicted in Supplementary Fig. 8B,C. **E** Serum antibody binding competition with several known HCV mAbs as measured by Luminex assay. Beads were coated with either AMS0232, H77, UKNP4.1.1, AMS3a or HepCon sE1E2.v4-LZ. Each column represents one rabbit. Stratification of competition is indicated. **F** Flow cytometry analysis of HEK293T cells transfected with mRNA encoding optimized membrane-bound E1E2 designs based on HepCon C2P2. Engagement of several HCV mAbs was determined for, and compared between, untransfected HEK293T cells (black line) and HEK293T cells transfected with HepCon C2P2 (orange), C2P2 with a SARS-CoV-2 TMD (C2P2-TMD) (light green) or a SARS-CoV-2 TMDdCT (C2P2-TMDdCT) (green). All signals have been normalized to single live cell counts. **G** (top) Groups of five female BALB/c mice were immunized at weeks 0 and 4 with 5 µg lipid nanoparticle (LNP) encapsulated mRNA encoding AMS0232 or HepCon C2P2-TMDdCT. At week 6 bleeds were taken and the spleens were harvested. (bottom) Groups of five female BALB/c mice were immunized at weeks 0 and 4 with 5 µg and at week 10 with 1µg LNP encapsulated mRNA encoding AMS0232 or HepCon C2P2-TMDdCT. At weeks 0, 4, 10 and 12 bleeds were taken. **H** ELISpot analysis of spleens harvested at week 6. E1E2-specific IFNγ positive spot forming cells (SFCs) per 10^6^ splenocytes were plotted for both groups. Means are indicated. **I,J** Mice serum ELISA endpoint binding titers to full-length AMS0232 and HepCon E1E2 for all four timepoints (**I**) and to full-length HepCon, H77, AMS0232, AMS3a and UKNP4.1.1 E1E2 for week 12 (**J**). in (**I**) the means and SD are indicated and in (**J**) medians are indicated. **K** Week 12 neutralization ID_50_-values of heterologous HCVpp JFH1. Medians are indicated. Significant differences between groups in **C**, **D**, **J** were determined using a Kruskal-Wallis test, followed by a Dunn’s post-test and in **H**, **K** using a Mann-Whitney test. Significant differences are indicated on top of the graphs (* <0.05; ** < 0.01). Figures **A**, **G** were created with Biorender.com.

We determined serum binding titers longitudinally against sequence-matched full-length membrane-bound versions of HepCon and AMS0232 to measure on-target E1E2 responses and exclude responses to the LZ domain (Fig. 2B, Supplementary Fig. 8A). After priming, binding responses were low or undetectable. After the boosts, rabbits immunized with HepCon sE1E2.v4-LZ induced stronger HepCon E1E2 serum binding responses (week 6: median endpoint titer: 23,442, week 22: 47,315) than AMS0232 at week 6 (800; Kruskal-Wallis *p* = 0.0088) and the cocktail at week 6 and week 22 (476; *p* = 0.0125; 6,383; *p* = 0.0449, respectively). Remarkably, HepCon sE1E2.v4-LZ and the cocktail induced similar anti-AMS0232 E1E2 responses, even though AMS0232 sE1E2.v4-LZ was part of the cocktail (Supplementary Fig. 8A). The rabbits immunized with AMS0232 sE1E2.v4-LZ elicited the highest AMS0232 E1E2 titers, but differences between the groups were not statistically significant at any time point.

To assess binding breadth at peak serum titer (week 22), we measured binding antibodies against full-length E1E2s of the other cocktail strains (H77, UKNP4.1.1 and AMS3a) (Fig. 2C). The cocktail group induced higher binding titers than the AMS0232 group against AMS3a (medians: 3,724 and 406, Kruskal-Wallis *p* = 0.0229) and UKNP4.1.1 E1E2 (10,139 and 430, *p* = 0.002), which belong to different genotypes than AMS0232. Strikingly, the HepCon group induced binding titers similar to the cocktail group against the cocktail sequence-matched E1E2s, indicating that HepCon sE1E2.v4-LZ can elicit serum binding responses against heterologous HCV strains.

Next, we measured serum neutralization against a panel of 13 diverse HCVpp (Supplementary Fig. 8B-D) at peak titer (week 22).^68^ We were not able to produce an infectious HepCon HCVpp to measure autologous HepCon neutralization. As reported previously, AMS0232 sE1E2.v4-LZ induced a relatively narrow response.^50^ Five out of six rabbit sera neutralized the autologous AMS0232 and the neutralization-sensitive JFH1 virus and some of the sera weakly neutralized AMS0230, AMS0229 and/or UKNP4.1.1 HCVpp.^68,69^ The rabbits in the cocktail group developed broader serum neutralization, although only one rabbit neutralized all autologous viruses from the cocktail. We also detected a few more serum neutralization hits (ID_50_>20) for the cocktail-immunized animals against additional heterologous viruses (UKNP2.4.1, UKNP5.2.1 and AMS4d.k9 HCVpp). Overall, the HepCon sE1E2.v4-LZ immunized animals showed a similar neutralizing breadth as the cocktail immunized animals. Sera of two HepCon sE1E2.4-LZ immunized rabbits (UA285 and UA286) respectively cross-neutralized 11/13 and 8/13 of tested HCVpps, including the heterologous AMS2b, UKNP2.2.1 and UKNP3.2.2 (UA285), which were not neutralized by any of the sera in the cocktail group (Supplementary Fig. 8C,D). To determine the overall breadth and potency for each animal, we calculated the global geometric mean titers (GGMT) of each rabbit serum based on the neutralization ID_50_-values in the 13-virus panel (Fig. 2D). The GGMT values for the rabbits in the HepCon group were higher than those in the monovalent AMS0232 group (geomeans of 48 and 26; Kruskall-Wallis *p* = 0.0043), and not statistically different from that of the tetravalent cocktail group (geomean 37, p=0.53).

To determine the specificities of the polyclonal rabbit Ab responses, we performed a Luminex competition binding assay using HepCon (Fig. 2E) and H77, UKNP4.1.1, AMS3a and AMS0232 sE1E2.v4-LZ as antigens (Supplementary Fig. 9).^69^ We detected competition with Abs targeting multiple epitopes. Strongest competition for HepCon sE1E2.v4-LZ binding was observed with CD81, with the HepCon group outperforming both cocktail and AMS0232 groups (medians: 66%, 17% and 13% competition, *p*=0.030 and p=0.0031, respectively). Competition was also observed for AT1211 (domain C), and somewhat weaker by 1382_01_H05 (AR3) and AR4A (AR4) to HepCon sE1E2.v4-LZ. Notably, we observed also relatively strong competition with non-neutralizing mAb CBH-4B, indicating that the non-neutralizing domain A epitope is immunogenic on HepCon sE1E2.v4-LZ.

Altogether, HepCon sE1E2.v4-LZ induced cross-neutralizing responses in rabbits at least as efficiently as a tetravalent cocktail vaccine and the polyclonal serum Abs targeted known bNAb and non-NAb epitopes.

### mRNA-encoded membrane-bound HepCon C2P2 induces an E1E2 specific immune response in mice

Cell surface expression is considered key for the immunogenicity of mRNA delivery of vaccine antigens, probably because it increases antigen valency to optimally stimulate B cells.^70,71^ However, native E1E2 contains an endoplasmic reticulum retention signal, leading to poor cell surface expression.^72–74^ We hypothesized that we could leverage the C2P2 design for membrane expression of HepCon sE1E2. We assessed compatibility of heterologous transmembrane domains (TMDs)^75^ for cell surface expression of AMS0232 C2P2 and identified the SARS-CoV-2 TMD (TMD truncated by 19 amino acids^76,77^) and TMDdCT (TMD without the cytoplasmic tail to decrease retrograde trafficking^78–80)^ as prime candidates, yielding C2P2-TMD and C2P2-TMDdCT (Supplementary Fig. 1B, 10A-C). We next tested soluble HepCon C2P2, HepCon C2P2-TMD and HepCon C2P2-TMDdCT mRNA constructs for cell surface expression and antigenicity using flow cytometry (Fig. 2F). Overall, cell surface expression was highest for the HepCon C2P2-TMDdCT. All tested mAbs bound to the two cell surface-expressed HepCon C2P2 constructs, including AR3- and AR4-targeting bNAbs and the non-neutralizing CBH-4B, indicating that cell-surface expression of HepCon C2P2 does not perturb accessibility of these antibody epitopes.

Next, we immunized mice twice at weeks 0 and 4 with 5 µg lipid nanoparticle (LNP) encapsulated mRNA encoding soluble HepCon C2P2, HepCon C2P2-TMD or HepCon C2P2-TMDdCT (Supplementary Fig. 10D). Four weeks after the prime, both C2P2-TMD and C2P2-TMDdCT induced detectable serum binding Abs against full-length HepCon E1E2, while soluble C2P2 did not (Supplementary Fig. 10E). After the boost, C2P2-TMD and C2P2-TMDdCT induced more than 1,000-fold higher binding titers than C2P2 (medians: 297,898 and 320,627, versus 216; Kruskal-Wallis *p* = 0.0267 and *p* = 0.0216, respectively). Next, we assessed serum binding breadth at week 7 against full-length AMS0232, H77, UKNP4.1.1 and AMS3a E1E2 (Supplementary Fig. 10F). Again, HepCon C2P2-TMDdCT induced 5-to 100-fold higher binding titers than soluble C2P2 (for H77, AMS0232, AMS3a and UKNP4.1.1: Kruskal-Wallis *p* = 0.0026, *p* = 0.0042, *p* = 0.0059 and *p* = 0.0031, respectively). The data suggested that HepCon C2P2-TMDdCT outperformed C2P2-TMD, but the differences were not statistically significant.

We assessed the neutralization capacity of the week 7 sera (Supplementary Fig. 10G,H). Serum neutralization potency was limited, but we did detect neutralization of the heterologous JFH-1 HCVpp by HepCon C2P2-TMD and C2P2-TMDdCT, but not for soluble C2P2 (median neutralization C2P2-TMD, 39%; C2P2-TMDdCT, 77%, and soluble C2P2, 2%, Kruskal-Wallis *p* = 0.0112). Taken together, this demonstrates that cell surface expression of HepCon C2P2 is crucial for the (early) induction of serum Ab responses in the context of a HepCon mRNA-LNP vaccine.

### HepCon C2P2-TMDdCT induces an E1E2 specific T cell response in mice

We next determined T cell responses and if a second boost would benefit serum Ab responses in T cell and B cell focused mouse immunogenicity studies, respectively. In both studies we used LNP encapsulated mRNA encoding HepCon C2P2-TMDdCT, or AMS0232 C2P2-TMDdCT as a control (Supplementary Fig. 10I), immunized two times (5 µg at weeks 0 and 4) or three times (5 µg at weeks 0 and 4; 1µg at week 10) (Fig. 2G). In the T cell study, E1E2-specific IFNγ postive spot forming cells (SFCs) were determined by an Enzyme-Linked Immunospot (ELISpot) assay using a HepCon-based peptide pool (Fig. 2H). The HepCon group induced a strong T cell response with a median of 449 SFC/1×10^6 splenocytes, which was significantly higher than for AMS0232 C2P2-TMDdCT, as expected. This indicates that HepCon C2P2-TMDdCT mRNA elicits E1E2 specific T cell responses.

For the B cell study, we did not detect higher serum binding titers against full-length HepCon E1E2 nor against H77, AMS3a and UKNP4.1.1 E1E2 compared to the previous study in which the mice were immunized only twice (Fig. 2I,J, Supplementary Fig. 10E,F). We observed median ID_50_-values of 314 for the HepCon group and 40 for the AMS0232 group for JFH1 HCVpp neutralization by week 12 sera. This indicates once more that membrane-anchored mRNA-encoded C2P2 can induce a NAb response in mice to some extent (Fig. 2K, Supplementary Fig. 10J).

Taken together, HepCon mRNA-LNPs induced E1E2-specific T cell responses and serum binding in mice. This further substantiated HepCon E1E2 as lead candidate for vaccine development.

### Recombinant HepCon sE1E2 heterodimers engage a broad panel of inferred bNAb precursors

To consistently induce human HCV bNAbs by vaccination, it is imperative to effectively target the germline precursor bNAb B cells.^38,43,44,81^ Considering the huge diversity within the human naïve B cell repertoire, a germline-targeting HCV glycoprotein immunogen should be able to engage many different potential precursor B cells to induce the rare desired bNAb B cell lineages in humans.^38,44,52,81–84^ However, most HCV glycoproteins engage no or only few HCV inferred germline (igl-)bNAbs.^43,44,50^ We reasoned that this is because most HCV glycoproteins are based on viral sequences from chronically infected individuals and have evolved to escape from neutralizing antibodies by preventing their binding and concomitantly also the binding of their precursors. In principle, an E1E2 consensus sequence should lack these strain-specific escape mutations and we hypothesized that HepCon E1E2 might therefore support igl-bNAb binding. Therefore, we next assessed whether HepCon sE1E2 binds HCV igl-bNAbs (Fig. 3A). We obtained the inferred germline sequences from a panel of HCV bNAbs from published literature^43,46^ or by inferring their sequences using the IMGT/V-QUEST software tool.^85^ (Supplementary Fig. 11A-C). Because of VDJ rearrangement, it is not always possible to reliably infer the CDRH3 sequence of a germline bNAb. Thus, we utilized a partially inferred germline CDHR3 when the germline D gene sequence could be reliably inferred and then added the N1 and N2 junction nucleotides from the mature bNAb. If the germline D gene could not be inferred reliably, we utilized the inferred VH gene allele and the mature CDRH3 sequence. For AR4A, which contains an insertion in its CDRH2 we generated three igl-versions: with CDRH2 insertion and mostly mature CDRH3 (igl-AR4A-im3), with insertion and germline inferred CDRH3 (igl-AR4A-ig3) and without insertion and germline inferred CDRH3 (igl-AR4A-g3).^12^ We determined the number of improbable amino acids mutations induced by somatic hypermutation using the ARMADiLLO webserver tool.^86^ We scored for each bNAb HC and light chain (LC) and highlighted the improbable mutations (<1% likelihood) in underscored bold (Supplementary Fig. 11A,B).^86,87^ Compared to HIV-1, HCV bNAbs require significantly fewer improbable mutations to acquire breadth (Supplementary Fig. 11D). AR3 bNAbs are mostly derived from “F allele” VH1-69 genes, which contain phenylalanine (F) at position 54.^88^ However, around 11% of the human population only express “L allele” VH1-69 genes that contain a leucine (L) at position 54, often accompanied by an arginine (R) at position 50, which seems detrimental for binding.^44,88^ Thus far, 1198_05_G10 is the only igl-bNAb derived of an L allele VH1-69 gene.^19^

**Figure 3:**
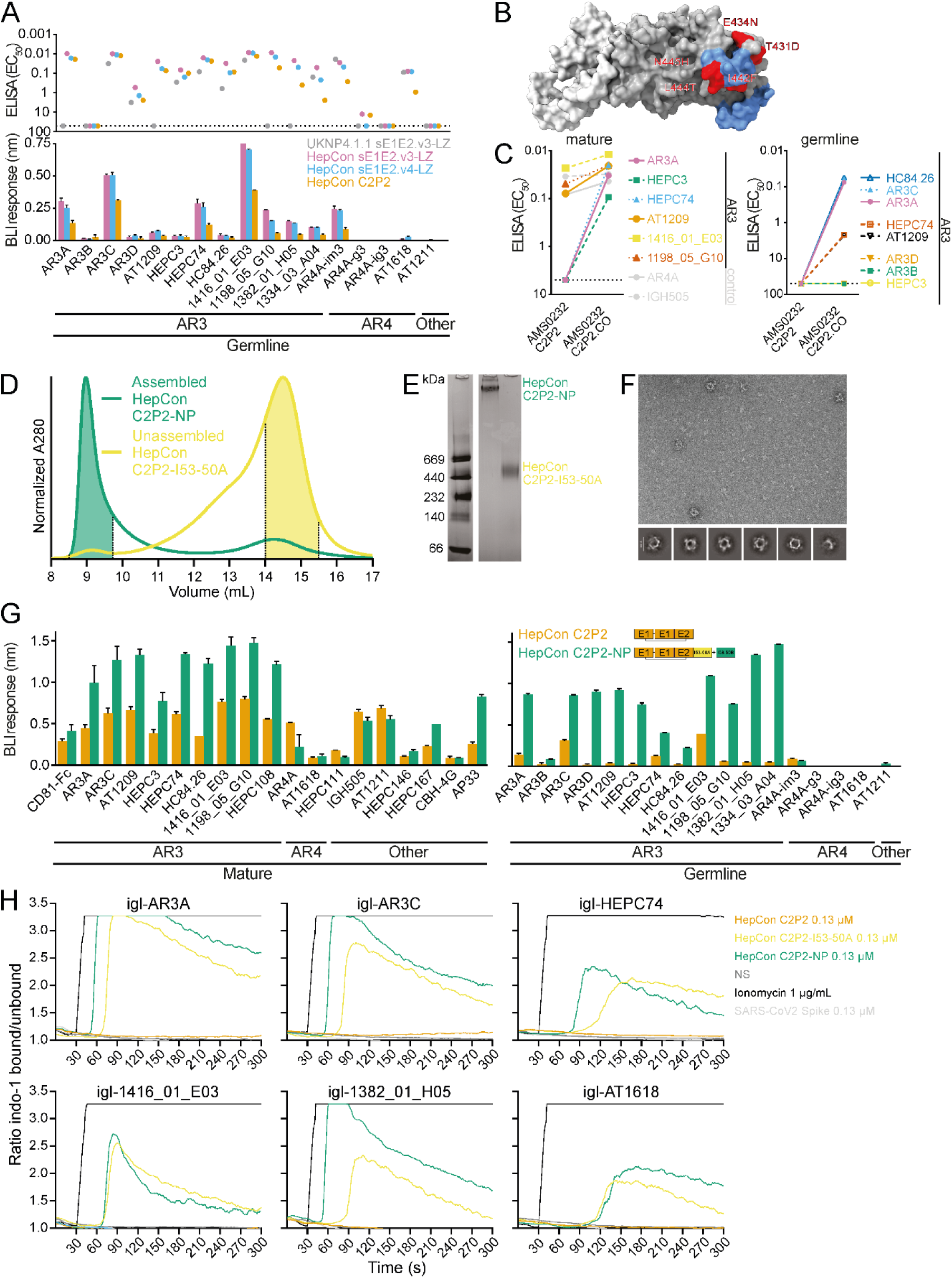
Engagement of igl-bNAbs, NP characterization and activation of HCV-specific B cells expressing igl-bNAbs. **A** Antigenicity profile of the HepCon sE1E2s and UKNP4.1.1 sE1E2v3-LZ probed by HCV igl-bNAbs. Top: EC_50_-values derived from Strep-TactinXT ELISA assays (Supplementary Fig. 12A) are shown per igl-bNAb. UKNP4.1.1 sE1E2.v3-LZ is known to bind several igl-bNAbs and was used for comparison.^50^ Bottom: maximum BLI response values derived from BLI assays measuring binding in nm over time (Supplementary Fig. 12B) are shown per igl-bNAb. **B** Surface representation of AMS0232 E1E2 (PDB: 7T6X) in which consensus optimized amino acids for the AMS0232 C2P2.CO design are depicted in red. The AR3 epitope is indicated in blue. **C** EC_50_-values derived from Strep-TactinXT ELISA assays (Supplementary Fig. 13) for eight mature bNAbs (left) and eight AR3-targeting igl-bNAbs (right) binding to AMS0232 C2P2 and AMS0232 C2P2.CO. **D** SEC profile of assembled HepCon C2P2-NP and unassembled HepCon C2P2-I53-50A trimer on a Superose 6 Increase 10/300 GL column. Pooled fractions are indicated. **E** BN-PAGE gel of HepCon C2P2-NP and HepCon C2P2-I53-50A. **F** NS-EM and 2D class averages of HepCon C2P2-NPs. **G** Antigenicity profile of HepCon C2P2, C2P2-I53-50A and C2P2-NP with CD81-Fc, HCV mature bNAb and igl-bNAb binding. The maximum BLI response values were derived from BLI assays measuring binding in nm over time (Supplementary Fig. 5B, 12B). Binding data of HepCon C2P2 from Figures 1D and 3A were plotted as comparison. **H** Activation of Ramos B cells expressing igl-bNAbs by HepCon C2P2, C2P2-I53-50A trimers and C2P2-NPs. Activation was determined by measuring calcium flux. Ionomycin was added as positive control and absence of stimulation and the SARS-CoV2 spike were used as negative controls. The SARS-CoV-2-specific COVA1-16 was used to validate the SARS-CoV2 Spike negative control (Supplementary Fig. 14D). All ELISAs and BLI assays used for Figure 3A and 3G were performed in duplo. The maximum binding signals in **A** and **G** were obtained from the full BLI curve and corrected for the mock (Supplementary Figures 5B and 12B).

We then performed ELISAs on the HepCon E1E2 antigens and used UKNP4.1.1 sE1E2.v3-LZ, our current best germline-targeting vaccine candidate, as a comparator (Fig. 3A, Supplementary Fig. 12A).^50^ The three HepCon sE1E2 antigens bound eleven out of twelve tested *VH1-69*-derived AR3 igl-bNAbs, while UKNP4.1.1 sE1E2.v3-LZ bound seven. Compared to UKNP4.1.1 sE1E2.v3-LZ, HepCon sE1E2 antigens also reacted to igl-AR3A, igl-AT1209, L-allelic igl-1198_05_G10 and igl-1382_01_H05. Furthermore, the HepCon proteins consistently bound more strongly to the seven *VH1-69*-derived AR3 igl-bNAbs that also bound UKNP4.1.1 sE1E2.v3-LZ. Both UKNP4.1.1 sE1E2.v3-LZ and the HepCon sE1E2 proteins also bound igl-AT1618 and igl-AR4A-im3, which target the AR4 epitope.

Next, we assessed igl-bNAb binding using BLI (Fig. 3A, Supplementary Fig. 12B) and found that binding in BLI correlated well with binding in ELISA and between the different iterations of HepCon sE1E2 antigens (Supplementary Fig. 6A,B). The strongest binding signals were detected for igl-AR3A, igl-AR3C, igl-HEPC74, igl-1416_01_E03, igl-1198_05_G10, igl-1382_01_H05, igl-1334_03_A04 and igl-AR4A-im3 for each of the HepCon antigens.

As expected, binding of each igl-bNAb was weaker than its mature counterpart, because of the difference in affinity maturation (Supplementary Fig. 12C). Lastly, fusion of HepCon C2P2 to TMD or TMDdCT to enable cell surface-expression did not perturb igl-bNAb binding (Supplementary Fig. 12D). In summary, stabilized HepCon-based sE1E2 heterodimers engage a wide range of igl-bNAbs against distinct conformational epitopes, which represents a significant improvement over current state-of-the-art E2 and sE1E2 antigens.^43,44,50^

### Antigenic optimization using HepCon-based residues improves (igl-)bNAb binding of sE1E2 from other isolates

Next, we wanted to determine if the broad igl-bNAb binding profile of HepCon sE1E2 can directly be attributed to its consensus residues. Therefore, we aimed to optimize binding of AR3 bNAbs to AMS0232 C2P2, by replacing five of its uncommon residues in AR3 by the consensus residues of HepCon (T431D, E434N, I442F, L444T and N445H) (Fig. 3B).^12,50,89^ The consensus optimized (CO) version of AMS0232 C2P2 (AMS0232 C2P2.CO) acquired binding to three AR3 bNAbs, showed enhanced binding to AT1209 and now measurably reacted to several AR3 igl-bNAbs (Fig. 3C, Supplementary Fig. 13), while AR4A binding was unaffected. This indicated that replacing strain-specific amino acids within bNAb epitopes by consensus residues can improve (igl-)bNAb binding. This CO-strategy might be useful for optimizing antigenicity profiles of existing HCV glycoprotein immunogens.

### Multivalent presentation of HepCon C2P2 on nanoparticles improves igl-bNAb binding

Multivalent immunogen presentation on nanoparticles (NPs) increases epitope avidity, which enhances BCR binding and cross-linking, which is especially important for engaging low affinity BCRs of naïve B cells.^90^ The clinically validated two-component I53-50 NP^91^ system is well-established to generate NPs for viral vaccines.^89,92–96^ We first produced a trimeric fusion protein consisting of HepCon C2P2 fused to the trimeric I53-50A component (HepCon C2P2-I53-50A). We then mixed HepCon C2P2-I53-50A trimers with the pentameric I53-50B component to induce assembly of icosahedral HepCon C2P2-NPs. Successful NP assembly was confirmed by SEC (Fig. 3D, Supplementary Fig. 2B) and Blue Native PAGE (BN-PAGE) (Fig. 3E). Negative-stain electron microscopy (NS-EM) on the SEC-purified NP fractions confirmed that the majority formed fully assembled particles of ∼30 nm in diameter. The NS-EM 2D class averages depicted a NP core with HepCon C2P2 densities on their surface, although some partially assembled species were visible (Fig. 3F).

We confirmed that the antigenicity of C2P2-I53-50A trimers and NPs was maintained using Strep-TactinXT and *Galanthus nivalis* lectin-coated ELISA, respectively (Supplementary Fig. 14A,B,C). To assess antigenicity of the NPs, we tested binding of bNAbs and igl-bNAbs using BLI and compared this to C2P2 (Fig 3G, Supplementary Fig. 5B, 12B). Overall, most mature mAbs showed increased binding signal to C2P2-NP compared to C2P2, especially for mAbs against AR3 and AS412 (AP33). Conversely, we measured lower binding signal for C2P2-NPs to mAbs against AR4, E1 (IGH505), domain A (CBH-4G) and domain C (AT1211). Thus, AR3 and AS412 are comparatively more accessible on the NPs than epitopes in E1, near the E1/E2 interface (AR4) and weakly or non-neutralizing epitopes located in the E2 back layer (domain A and C). For igl-bNAbs, the differences were more pronounced (Fig. 3G). The HepCon C2P2-NPs displayed much higher binding signals than soluble HepCon C2P2 for igl-AR3A, igl-AR3D, igl-AT1209, igl-HEPC3, igl-1198_05_G10, igl-1382_01_H05 and igl-1334_03_A04 with virtually no binding dissociation (Supplementary Fig. 12B). The HepCon C2P2-NPs afford sufficient epitope avidity to enable binding of eleven out of twelve tested *VH1-69*-derived AR3 igl-bNAbs, including the L allelic 1198_05_G10 igl-bNAb.

### HepCon C2P2-NPs activate inferred germline HCV-specific B cells *in vitro*

To determine if HepCon C2P2 proteins can activate precursor B cells, we measured Ca^2+^ dependent activation of Ramos B cell lines expressing AR3-targeting igl-AR3A, igl-AR3C, igl-HEPC74, igl-1416_01_E03, igl-1382_01_H05 and the AR4-targeting igl-AT1618 as BCR (Fig. 3H). At 0.13 µM, both the HepCon C2P2-NPs and C2P2-I53-50A trimers activated all six igl-bNAb B cell lines, with the NPs displaying the highest activation. In contrast, soluble HepCon C2P2 induced weak or no activation of these B cell lines. Soluble HepCon C2P2 did activate B cells expressing mature AR3C and HEPC74 BCRs, but less efficiently than the trimers and NPs (Supplementary Fig. 14D). Strongest activation was observed for the B cell lines expressing igl-AR3A, igl-AR3C and igl-1382_01_H05 BCRs. Notably, HepCon C2P2-NPs and C2P2-I53-50A trimers activated igl-AT1618 B cell line, even though the NP failed to induce a measurable binding signal on BLI (Fig. 3G, Supplementary Fig. 12B).

In summary, AR3- and AR4-targeting inferred precursor B cells can be activated by multivalently displayed HepCon sE1E2 antigens. These findings further illustrate that HepCon sE1E2 is a promising antigen for (multi-epitope) germline-targeting strategies.

### Cryo-EM structure of the inferred germline precursor of 1416_01_E03 bNAb in complex with HepCon sE1E2.v4-LZ

The 1416_01_E03 bNAb is an exceptionally broad *VH1-69*-derived AR3 bNAb, which was isolated from a single human B cell from an elite HCV neutralizer.^19^ Because of its high nucleotide identity (92%) to its inferred germline precursor and its average CDRH3-loop length of 15 amino acids (Kabat classification) it is an attractive candidate for germline targeting. To determine its epitope and binding mode, we used cryo-EM to solve structures of HepCon sE1E2.v4-LZ in complex with 1416_01_E03 and its germline precursor at 3.4 Å and 3.5 Å, respectively (Fig. 4A, Supplementary Fig. 15A-F). The structure of E2-1416_01_E03 was obtained from a bigger complex generated to attempt to solve the structure of E1E2 in complex with AT1211, 1416_01_E03 and the AR4-targeting AT1618 bNAb.^17^ The complex contained all three antibodies (Supplementary Fig. 15G), but presented severe orientation bias in cryo-EM, which despite our best efforts we were unable to correct in order to solve the epitope of AT1618 at high resolution.

**Figure 4:**
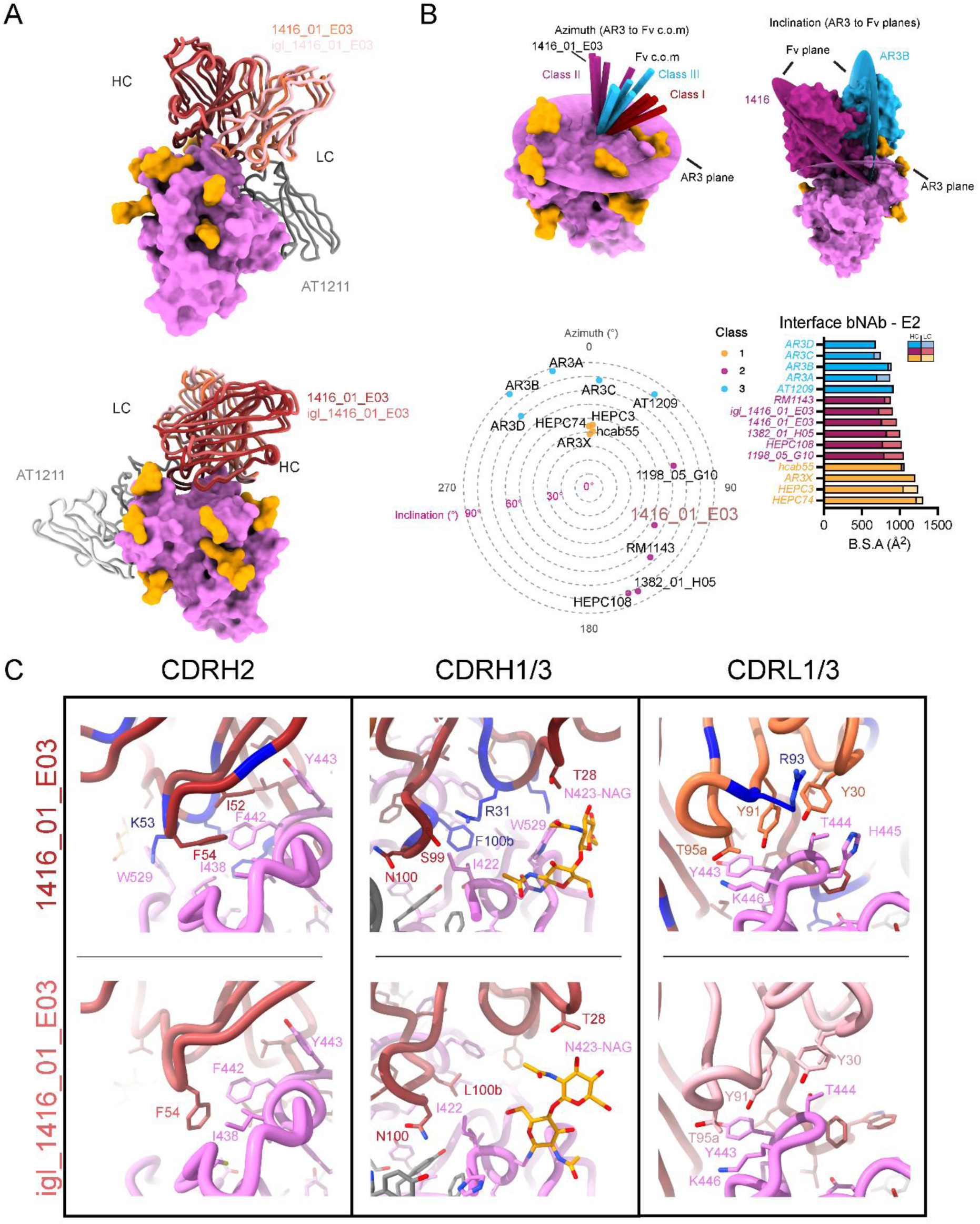
Cryo-EM of HepCon sE1E2.v4-LZ in complex with bNAb AT1211 and 1416_01_E03 or igl-1416_01_E03. **A** Cryo-EM model of AT1211 and 1416_01_E03 or igl-1416_01_E03 bNAb in complex with HepCon sE1E2.v4-LZ. Both bNAbs are represented as ribbon, with AT1211 heavy and light chains colored grey and light grey respectively, 1416_01_E03 / igl-1416_01_E03 heavy and light chains colored shades of red and pink, respectively. E2 is represented as a violet surface model, with modelled glycans in orange. **B** Distribution of azimuth (top left) and inclination angles (top right) for multiple AR3-binding antibodies, using defined classes^47^, and plotted in an alt-azimuth plot (bottom left). See methods for definitions and calculations of planes and angles. For the alt-azimuth plot, azimuth are all compared to Class-I HEPC74 mAb defined as 0°, and every other mAb measured counter clockwise relative to it. Buried surface area between each of these antibodies and E2 is also indicated, for each heavy and light chains, colored and sorted by class (bottom right). PDB accession codes used for this analysis: 4MWF (AR3C), 6BKB (AR3A), 6BKC (AR3B), 6BKB (AR3D), 6MEH (HEPC74), 6MEJ (HEPC3), 6URH (AR3X), 7JTG (RM1143), 7RFB (1198_05_G10), 7RFC (1382_01_H05), 8W0V (hcab55), 9O3D (HEPC108). **C** Comparison of bNAb 1416_01_E03 and igl-1416_01_E03 binding E2, showing the same point of view for each panel, comparing interactions from CDRH2, CDRH1/3 and CDRL1/3. Somatically mutated residues on 1416_01_E03 are colored in dark blue. Heteroatoms colored on stick representation as oxygen (red) and nitrogen (blue).

1416_01_E03 shares roughly the same epitope on E2 as AT1209, centered on the small helix (437-442) in AR3, but their approach angles differ substantially. AT1209 utilizes its elongated hydrophobic CDRH2 to reach into a pocket which includes residues I422, L427, F442, V515 and Y613 and is located between the E2 ß-sandwich and back layer, while 1416_01_E03 utilizes its CDRH3 to bind to the same pocket (Supplementary Fig. 15H). To better understand the different rotational parameters, we defined an AR3 epitope plane on E2 and measured direction of approach (azimuth) and inclination of the HC-LC planes relative to the epitope plane for 1416_01_E03 and other frontlayer (FRLY) bNAbs (Fig. 4B). We confirmed the three classes of FRLY bNAbs as defined before, and observed that 1416_01_E03 belongs to class II.^47^ 1416_01_E03 makes use of both its HC and LC to bind to E2 (Fig. 4B). The HC of 1416_01_E03, including the CDRH3, binds the small helix of AR3 for 437 to 442, while the lambda LC completes the interface by wrapping its CDRL1 and CDRL3 around 442-445, with Y30 interacting with T444 and H445, and Y91 contacting Y443 as well as participating in a network of hydrogen bonds between Y91, T95A and the E2’s K446 (Fig. 4C). Indeed, we found a substantial drop in binding of the 1416_01_E03 Fab when coupled with other lambda LCs (Supplementary Fig. 15I). In contrast, the LCs of AT1209 and many other VH1-69 AR3 bNAbs have no or only few interactions with their target epitope.^44^

Most residues involved in these interactions are germline residues and are conserved in 1416_01_E03 (Fig. 4C). However, some somatic mutations seem key in altering and improving the binding to E2, most prominently L100bF in the CDRH3, which is able to extend to reach back to this 422-428 region and generate more contacts (Supplementary Fig. 15J). Worthy of notice, the neighboring W100aS results in a lost tryptophan deeply embedded in the hydrophobic pocket in E2, but this liberates the CDRH3-loop, which allows it to move closer to E2 (movement of 1.6 Å of the α-carbon at position 100a between 1416_01_E03 and igl-1416_01_E03), promoting more contacts with the antigen (igl-1416_01_E03 CDRH3-E2 B.S.A: 363 Å^2^; 1416_01_E03 CDRH3-E2 B.S.A: 410 Å^2^). This also slightly alters the approach angle, with a Δ° difference of ∼2° between mature and igl-1416_01_E03 and ∼3.3 Å displacement ΔC at the distal end of the fragment variable domain, measured at the C-terminal residue of each chain (Supplementary Fig. 16A). This deviation is not due to a misalignment, since under the same alignment to E2 both models have AT1211 bNAb bound with nearly identical pose (Δ°= 0.4°, ΔC = 1.4 Å). Other mutations towards the front layer involve the improbable somatically mutated I53K (Supplementary Fig. 11A) extending towards the highly conserved W529 on E2 (Supplementary Fig. 16B). Additionally, S31R seems to stabilize the CDRH3 F100b residue by forming a network of π-stacking interactions that also includes somatically mutated Y32F residue. The somatically mutated G93R in the LC coordinates with carbonyl and backbone oxygens on the helix between S27a and Y31, potentially adding conformational stability to this E2-targeting region (Fig. 4C).

Lastly, (igl-)1416_01_E03 is in close proximity to AT1211, although the contacts are different compared to AT1211 and AT1209. A network of hydrogen bonds is formed between both the CDRH3 and CDRL2 of (igl-)1416_01_E03 and AT1211 (Supplementary Fig. 17). This once again shows the potential for bNAbs to maximize the area they occupy on an antigen and possibly synergize with one another. This suggests AT1211 might have biological relevance in interacting with AR3-targeting Abs.

## Discussion

Here we described the design and characterization of stabilized soluble consensus based native-like sE1E2 proteins, which induce heterologous (N)Ab responses in rabbits and mice. HepCon sE1E2 engages a wide range of igl-bNAbs and activates B cell lines expressing igl-bNAbs *in vitro* when multivalently presented on NPs. Finally, we structurally characterized the broad 1416_01_E03 bNAb and its inferred germline precursor. We propose HepCon E1E2 as a promising germline-targeting HCV vaccine candidate.

Antigenic consensus-based immunogens have been designed against HCV (using E2 only)^97^, influenza (hemagglutinin)^98^ and HIV-1 (envelope glycoprotein)^52,99^. Per definition, consensus sequences are most related to all other sequences within a population and consensus-based antigens might therefore be more prone to induce cross-reactive responses. The majority of HCV antigens are based on sequences from chronic infection that have developed escape mutations to thwart the early (germline precursor) NAb response. A consensus sequence effectively “averages away” these isolate-specific escape residues and we postulate that this explains why HepCon engages so many igl-bNAbs. A similar consensus-based approach might be a useful starting point for the design of germline-targeting antigens against other sequence diverse (viral) pathogens.

Targeting multiple E1E2 epitopes enhances (serum) neutralization breadth and increases the threshold for escape variants to develop.^40,41,100,101^ Additionally, a recent germinal center modeling study predicted that priming of multiple bNAb lineages with a single immunogen is potentially additive.^102^ Together, this suggests a benefit of using E1E2-based antigens, such as HepCon E1E2, for germline-targeting compared to E2 alone. Furthermore, besides AR3 and AR4, novel HCV neutralizing epitopes are still being uncovered^20,103^ and a consensus E1E2 likely increases the chance that Abs induced against such epitopes are cross-recognizing.

Nevertheless, HepCon E1E2 could benefit from targeted (epitope) optimization. Viral isolates could be screened to identify key residues, or directed evolution could be used to further optimize binding by igl-bNAbs, such as igl-AR3B.^104^ Additionally, improving glycan occupancy through glycan engineering may be necessary to shield partially exposed non-neutralizing epitopes.^105^ Sera from HepCon sE1E2.v4-LZ immunized rabbits competed strongly with binding by the non-neutralizing CBH-4B Ab which shows HepCon sE1E2 presents an epitope *in vivo* that is not accessible on native-like E1E2.^12,60,64,106^ Additional antigen engineering is necessary to reduce off-target responses and promote a more focused NAb response.

Furthermore, the sequence of HepCon E1E2 is biased with 4/11 sequences being derived from genotype 1 (H77, Con1a, −1b and −1c). A next-generation HepCon E1E2 sequence could use a weighting strategy to minimize oversampling of certain genotypes and utilize contemporary sequences to better represent currently circulating strains. Alternatively, contemporary geographically tailored (combinations of) consensus sE1E2s could be generated to more closely resemble regional differences in geno- and subtype prevalence.^107^ The described design templates could be implemented to generate such regionally tailored consensus sE1E2s. Implementing an mRNA vaccination platform would in turn speed up downstream processing for GMP production, lowering costs for the implementation of these geographically tailored vaccination regimens.

Display on NPs enhanced binding of C2P2 to AR3 bNAbs, but did not enhance binding to AR4 bNAbs in BLI. This shows that AR4 is less accessible on I53-50 NPs and it might affect how efficiently these epitopes are targeted.^92,108^ On the other hand, multivalent presentation of C2P2 on NP was necessary to efficiently activate B cells expressing igl-bNAbs *in vitro*, including the AR4-targeting igl-AT1618 (Fig. 3H). This would suggest that B cells with low affinity BCRs against less accessible epitopes can still be targeted as long as the antigen is repetitively presented.

Two HepCon immunized rabbits showed *bona fide* neutralizing breadth (Supplementary Fig. 8C,D). It is key to determine how to induce these broader responses more consistently. The C2P2-NP might provide the avidity required to do so. Additionally, limiting off-target responses might be important to drive a consistently broader response.

Most HCV bNAbs require little affinity maturation, suggesting only few immunizations should be necessary to acquire breadth if the appropriate B cell precursors are engaged. However, the animal models used in this study are not suitable to assess whether HepCon E1E2 activates the desired B cell precursors *in vivo*, since they lack (orthologues of) human antibody genes, such as *VH1-69*. Humanized knock-in animals expressing *VH1-69*, or specific igl-bNAbs as BCRs, would provide better platforms to assess germline-targeting potential.^35^ Additionally, non-human primates express *VH1.36* and *VH1-138*01*, which are homologs of human *VH1-69* alleles, which enables evaluation of HepCon E1E2 within a human-like competitive B cell pool.^51,109,110^

For consideration in humans, it is important to determine a suitable, safe and practical vaccine platform. The advent of mRNA vaccine technology has been transformative for rapidly advancing vaccines in humans.^111^ We tested HepCon C2P2 as mRNA vaccines in mice, and found that the membrane-bound version was superior to the soluble version, in line with previous findings on influenza HA and HIV-1 Env.^71,112^ This probably pertains to the fact that cell surface expression enables higher antigen density for more efficient B cell activation. Still, we observed limited neutralization potency and breadth after three immunizations. Optimization of the mRNA immunogen, for example by incorporating pseudouridine into the mRNA^113,114^ and/or optimization of the timing of the vaccination regimen^115–118^ might be required to enhance immunogenicity.

Previous structures of HCV bNAbs have been important for establishing the antigenic landscape of E2 and E1E2, but these structures did not show how the precursors of these bNAbs bind the epitopes, which would provide useful insights on how to selectively prime the desired bNAb lineages.^12,46,47,57,60,64^ Therefore, we exploited HepCon sE1E2 to determine the epitope of 1416_01_E03 bNAb and how an inferred precursor engages its epitope on E2. Obtaining additional structures of HepCon E1E2 with other igl-bNAbs will be valuable for engineering immunogens to enhance the engagement of HCV bNAb B cell precursors.

HepCon E1E2 will undoubtedly also be a useful tool for isolating bNAbs from vaccinated animals, HCV-infected individuals or human volunteers. Because its antigenic surface consists of conserved amino acids, it is probably well-suited to discover novel neutralizing epitopes. While we did show that HepCon displays the same E2 fold as wild-type E1E2 and AT1618 complexing indicated native-like E1E2 folding, visualization of full sE1E2 heterodimers remained troublesome, possibly because of remaining inherent flexibility of the complex. Therefore, further stabilization might be required to obtain a consistent platform for full sE1E2 structural validation.

In conclusion, HepCon sE1E2 is a promising vaccine candidate, because it engages a wide range of human inferred precursor HCV bNAbs and because it is compatible with different vaccine platforms, including clinically approved I53-50 nanoparticles and mRNA-LNPs. Furthermore, it is probably a useful tool for a wide array of immunological applications. Finally, the consensus-based design strategy might also be a useful starting point for the design of germline-targeting immunogens against other rapidly evolving pathogens.

## Methods

### HepCon sequence design

The sequence for H77 E1E2 (GenBank ID: AF009606) and E1E2 consensus sequences for subtypes 1a, 1b, 1c, 2a, 2b, 3a, 4a, 5a, 6a and 6k were obtained from the Los Alamos hepatitis C sequence database.^53^ Using the Jalview software tool^122^ the HepCon E1E2 consensus sequence was made from these 11 sequences, by determining the most prevalent amino acid per position. After obtaining this HepCon E1E2 sequence, the following residues were removed: EDK after position 474, RPV after position 571, LTWPAND after position 578 and LENLVIL at position 748.

### HepCon rarity analysis

A residue rarity analysis for the concatenated E1E2 protein was performed using a multiple sequence alignment generated using MAFFT v7.526 of around 15000 E1E2 protein sequences from the Los Alamos National Laboratory HCV sequence database.^53^ For each position in the alignment, the ‘positional rarity’ in sequences of interest, such as HepCon and H77, were calculated as the proportion of non-gap sequences in the alignment having the same residue as the sequence of interest. A per-genotype weighting factor was included in this calculation to counteract the bias induced by the vastly differing numbers of sequences per genotype in the database. These positional rarity scores were used to produce the positional rarity histogram as well as the 3D protein structure rarity representations. Custom Python scripts used for these analyses are available at https://gitlab.com/ruben.walen/hepcon-germline-targeting.

### Constructs

All E1E2 constructs are numbered according to the standard H77 polyprotein numbering (GenBank ID: AF009606).^123^ All recombinant E1E2 designs in this study, including those used as subunit of C2P2-I53-50A or C2P2-NP, include the N-terminal HVR1 (384–410) and were truncated at position 714 at the C-terminus. The construct UKNP4.1.1 sE1E2.v3-LZ was previously described by our group^50^ and will therefore not be described here. The designs of sE1E2.v3-LZ, v4-LZ and C2P2 were described recently by our group.^50^ All constructs contained a TwinStrepII-tag (SAWSHPQFEKGGGSGGGSGGSSAWSHPQFEK) to facilitate purification. Ab sequences were human codon-optimized and cloned into human IgG1 expression vectors for the corresponding HCs or LCs as described before (Genscript Biotech).^44,124,125^ Most mature and some germline inferred Ab sequences were retrieved from previously published papers.^43,46^ The LC of HEPC3 and HEPC74, and the HC and LC of AT1209, HC84.26, 1416_01_E03, 1198_05_G10, 1382_01_H05, 1334_03_A04, AR4A, AT1618, HEPC111, AT1211 and AR5A were inferred using IMGT/V-QUEST software tool^85,121^ which predicts the most likely V, D and J genes and arrangement based on the nucleotide sequence of an Ab. The inferred germline AT1209 sequences were a kind gift of Tim Beaumont and Sabrina Merat (AIMM therapeutics). All amino acid sequences are available in Supplementary Figures 1 and 11. The ARMADiLLO webtool was used to score likeliness of mutations by somatic hypermutation for all used HC and LC sequences.^86^

### Structure prediction and visualization

Molecular graphics and analyses were performed with UCSF ChimeraX, developed by the Resource for Biocomputing, Visualization, and Informatics at the University of California, San Francisco, with support from National Institutes of Health R01-GM129325 and the Office of Cyber Infrastructure and Computational Biology, National Institute of Allergy and Infectious Diseases.^126–128^ For predicting the structures of HepCon C2P2 AlphaFold-2.2.0 Multimer was used.^67,129,130^ Code available at: https://github.com/deepmind/alphafold.

### HCV glycoprotein and antibody expression and purification

The expression and purification of the sE1E2.v3-LZ, sE1E2.v4-LZ, C2P2, and C2P2-I53-50A glycoproteins and mAbs was performed as described before.^89^ Suspension HEK293F cells (Invitrogen, cat no. R79009) were maintained in FreeStyle medium (Life Technologies). Cells were transfected using 1 mg/mL PEI MAX (Polysciences Europe GmBH, Eppelheim, Germany) and DNA plasmid in a 3:1 (w:w) ratio at a density of 0.8–1.2 million cells/ mL. To facilitate complete cleavage between E1-E2, sE1E2 or sE1E2-I53-50A were co-expressed in a 1:1 ratio with a plasmid expressing furin. The supernatant was harvested 6 days after transfection, centrifuged, and filtered using Steritops (0.22 μm pore size; Millipore, Amsterdam, The Netherlands). BioLock (IBA Life Sciences) and 1:10 (v:v) 10x Buffer W (1.0 M Tris-HCl, 1.5 M NaCl, 10 mM EDTA, pH 8.0) were added to the supernatant and 750 µL Strep-TactinXT beads (IBA life sciences) were added after which this was rolled overnight at 4 °C. The following day the proteins were purified using Strep-TactinXT columns (IBA Life Sciences) by gravity flow. The proteins were eluted with Buffer BXT (IBA life sciences) and concentrated in 30-kDa cut-off Vivaspin20 filters (Sartorius) in TBS (sE1E2) or TBS/5% glycerol (C2P2-I53-50A). Subsequently, the proteins were run over a Superdex 200 Increase 10/300 GL or Superose 6 Increase 10/300 GL SEC column (GE Healthcare). mAbs were produced by co-transfecting the plasmids expressing IgG1 HC and LC in HEK293F cells in a 1:1 ratio using a 3:1 ratio of PEI MAX and DNA. The HEK293F supernatants were harvested 5 days post-transfection, centrifuged, and filtered using 0.22 μm Steritop filters. 1 mL immobilized protein G beads (Pierce) were added to the supernatant and rolled overnight at 4 °C. The following day the mAbs were purified using protein G column (Pierce), followed by extensive washing with PBS, after which the mAbs were eluted with 18 mL 0.1 M glycine pH 2.5 directly captured in 2 mL neutralization buffer (1 M TRIS pH 8.7). The purified mAbs were buffer-exchanged to PBS using 30 kDa VivaSpin20 columns (Sartorius). Protein and Ab concentrations were measured with Nanodrop (ThermoScientific, Wilmington DE, USA) using the theoretical molecular weight and extinction coefficients for the proteins and IgG molecular weight and extinction coefficient for the mAbs. The proteins were stored at −80 °C and the mAbs at −20 °C until further analysis.

### Full-length membrane bound E1E2 production and purification

Full-length membrane bound E1E2 production and purification was performed similarly as described previously.^50^ HEK 293T cells were transfected using 40 μg of E1E2 expression plasmids, 80 μL of Lipofectamine 2000 and Opti-MEM (Invitrogen by Thermo Fisher Scientific) per T150 flask. Three days after transfection, the supernatants were removed and the cells were washed twice with PBS before detaching them with Trypsin-EDTA. Non-transfected HEK-293Tcells were taken along as a negative control. Cell suspensions were spun down and washed with PBS before counting them. Cells were resuspended in 1% Triton buffer (50 mM tris pH 8.0 + 150 mM NaCl + 1% Trition) with 1x proteinase inhibitor cocktail (Thermo Scientific), 1 mL per 1 × 10^6^ cells. After 30 min incubation at 4 °C with rotation, cell lysates were centrifuged at 4000 g at 4 °C for 30 min.

Purification of full-length E1E2 was done with Galanthus Nivalis Lectin (GNL) (Vector laboratories, l-1240–5) columns. Briefly, cell lysates of transfected and non-transfected cells were diluted 1:3 with PBS and added to previously washed GNL columns after which the columns were washed three times with PBS + tritonX100 0.1%. Full-length E1E2 was eluted from the column with 1.0 M alpha-d-mannopyranoside in PBS pH 7.5. The full-length E1E2 were concentrated using Vivaspin 100 kDa filters (Sartorius).

### SDS-PAGE

SDS-PAGE analysis was performed similarly as described previously.^50^ Approximately 5 µg of sE1E2 protein was mixed with loading dye (25 mM Tris, 192 mM Glycine, 20 % v/v glycerol, 4 % m/v SDS, 0.1 % v/v bromophenol blue in milli-Q water) and incubated for 10 minutes at 95 °C. Afterwards the complete samples were loaded onto a 4–12 % Tris-Glycine gel (Invitrogen). For a reducing SDS-PAGE, dithiotreitol (DTT; 100 mM) was added to the protein-loading dye mixture, incubated for 10 minutes at 95 °C and loaded onto a 4–12 % Tris-Glycine gel (Invitrogen). Gels were run at 125V for ∼75 minutes in a buffer containing 25 mM Tris, 192 mM glycine and 0.5 % SDS. Afterwards, coomassie blue staining of the gels was performed using PageBlue Protein Staining Solution (Thermo Fisher Scientific), followed by thorough destaining in MQ.

### BN-PAGE

BN-PAGE analysis was performed as described previously.^131^ In short, the proteins were mixed with loading dye (500µL 20x MOPS Running Buffer (1M MOPS + 1M Tris, pH 7.7) + 1000µL Ultrapure Glycerol (Invitrogen cat#15514C011) + 50 µL Coomassie Brilliant Blue G-250 + 600µL milli-Q Water) and directly loaded onto a 3-12% Bis-Tris NuPAGE gel. The gels were run for 2h at 200V (0.07 A) using Anode-Buffer (20x NativePAGE Running Buffer (Invitrogen) in milli-Q water) and Cathode-Buffer (1% NativePAGE Cathode-Buffer Additive (Invitrogen) in Anode-Buffer (Invitrogen). BN-PAGE gels were stained using the Colloidal Blue Staining Kit (Life Technologies).

### Site-specific glycan analysis

Site specific glycan analyses were performed as described before.^89^ To assess the glycan shield composition of our recombinant sE1E2 proteins, the proteins were separately digested with chymotrypsin and alpha-lytic protease, yielding peptides and glycopeptides containing a single N-linked glycan site. Site-specific composition and occupancy of these glycopeptides were determined by LC-MS using an Orbitrap Eclipse mass spectrometer. These analyses were performed as described previously.^50^

Two aliquots of each sample were denatured and reduced for 1h in 50 mM Tris/HCl, pH 8.0 containing 6 M of urea and 5 mM dithiothreitol (DTT). Next, sE1E2 proteins were reduced and alkylated by adding 20 mM iodoacetamide (IAA) and incubated for 1h in the dark, followed by a 1h incubation with 20 mM DTT to eliminate residual IAA. The alkylated sE1E2 proteins were buffer exchanged into 50 mM Tris/HCl, pH 8.0 using Vivaspin columns (3 kDa) and two of the aliquots were digested separately overnight using chymotrypsin (Mass Spectrometry Grade, Promega) or alpha lytic protease (New England Biolabs) at a ratio of 1:30 (w/w). The next day, the peptides were dried and extracted using Oasis PRiME HLB µElution plate (Waters). The peptides were dried again, re-suspended in 0.1% formic acid, and analyzed by nanoLC-ESI MS with an Ultimate 3000 HPLC (Thermo Fisher Scientific) system coupled to an Orbitrap Eclipse mass spectrometer (Thermo Fisher Scientific) using stepped higher energy collision-induced dissociation (HCD) fragmentation. Peptides were separated using an EasySpray PepMap RSLC C18 column (75 µm × 75 cm). A trapping column (PepMap 100 C18 3μm particle size, 75μm × 2cm) was used in line with the LC prior to separation with the analytical column. The LC conditions were as follows: 280-minute linear gradient consisting of 4-32% acetonitrile in 0.1% formic acid over 260 minutes followed by 20 minutes of alternating 76% acetonitrile in 0.1% formic acid and 4% ACN in 0.1% formic acid, used to ensure all the sample had eluted from the column. The flow rate was set to 300 nL/min. The spray voltage was set to 2.5 kV and the temperature of the heated capillary was set to 40 °C. The ion transfer tube temperature was set to 275 °C. The scan range was 375−1500 m/z. Stepped HCD collision energy was set to 15, 25 and 45% and the MS2 for each energy was combined. Precursor and fragment detection were performed using an Orbitrap at a resolution MS1= 120,000. MS2= 30,000. The AGC target for MS1 was set to standard and injection time set to auto which involves the system setting the two parameters to maximize sensitivity while maintaining cycle time. Full LC and MS methodology can be extracted from the appropriate Raw file using XCalibur FreeStyle software or upon request.

Glycopeptide fragmentation data were extracted from the raw file using Byos (Version 4.6; Protein Metrics Inc.). The glycopeptide fragmentation data were evaluated manually for each glycopeptide; the peptide was scored as true-positive when the correct b and y fragment ions were observed along with oxonium ions corresponding to the glycan identified. The MS data was searched using the Protein Metrics 305 N-glycan library with sulfated glycans added manually. The relative amounts of each glycan at each site as well as the unoccupied proportion were determined by comparing the extracted chromatographic areas for different glycotypes with an identical peptide sequence. All charge states for a single glycopeptide were summed. The precursor mass tolerance was set at 4 ppm and 10 ppm for fragments. A 1% false discovery rate (FDR) was applied. The relative amounts of each glycan at each site as well as the unoccupied proportion were determined by comparing the extracted ion chromatographic areas for different glycopeptides with an identical peptide sequence. Glycans were categorized according to the composition detected.

HexNAc(2)Hex(10+) was defined as M9Glc, HexNAc(2)Hex(9−5) was classified as M9 to M3. Any of these structures containing a fucose were categorized as FM (fucosylated mannose). HexNAc(3)Hex(5−6)X was classified as Hybrid with HexNAc(3)Hex(5-6)Fuc(1)X classified as Fhybrid. Complex-type glycans were classified according to the number of HexNAc subunits and the presence or absence of fucosylation. As this fragmentation method does not provide linkage information compositional isomers are grouped, so for example a triantennary glycan contains HexNAc5 but so does a biantennary glycan with a bisect. Core glycans refer to truncated structures smaller than M3. M9Glc-M4 were classified as oligomannose-type glycans.

### Enzyme-Linked Immunosorbent Assay (ELISA)

To assess binding of bNAbs and igl-bNAbs to the constructs, the Strep-TactinXT and SEC purified proteins were tested on 96-well Strep-TactinXT coated microplates (IBA LifeSciences). These Strep-TactinXT ELISA analyses were performed as described previously, with some minor adjustments.^50^

Purified TwinStrepII-tagged sE1E2 heterodimers (1 μg/mL in TBS) were coated for 1 h at room temperature or overnight at 4 °C on 96-well Strep-TactinXT coated microplates (IBA LifeSciences). Plates were washed with TBS three times before incubating with serially diluted mAbs in casein blocking buffer (Thermo Fisher Scientific) for 90 min. After three washes with TBS, a 1:3000 dilution of HRP-labeled goat anti-human IgG (Jackson Immunoresearch) in casein blocking buffer was added for 45 min. After washing the plates five times with TBS + 0.05% Tween-20, plates were developed by adding develop solution [1% 3,3′,5,5′-tetraethylbenzidine (Sigma-Aldrich), 0.01% H2O2, 100 mM sodium acetate, 100 mM citric acid] and the reaction was stopped after 3.5 min by adding 0.8 M H2SO4. Absorbance was measured at 450 nm. EC_50_ values were calculated using Graphpad Prism 9.5.1.

To assess binding of bNAbs and igl-bNAbs to HepCon C2P2-NP, a Galanthus nivalis lectin ELISA was performed, similar as described earlier with some alterations.^89^ Half-well 96-well plates were coated with 50 μL Galanthus nivalis lectin (Vector Laboratories) at 20 μg/mL in 0.1 M NaHCO3 pH 8.6. The next day, the plates were blocked with casein blocking buffer (Thermo Fisher Scientific) for 30 min. Afterwards the plates were washed twice with TBS and then 50 μL 1 μg/mL antigen was added to the lectin-coated plates. After 2 h, the plates were washed twice with TBS and incubated with 50 μL serially diluted mAbs in casein blocking buffer for 2h. The plates were washed three times with TBS and 50 μL of a 1:3000 dilution of HRP-labeled goat anti-human IgG (Jackson Immunoresearch) in casein blocking buffer was added for 1h. After washing the plates five times with TBS + 0.05% Tween-20, plates were developed by adding 50 μL develop solution [1% 3,3′,5,5′-tetraethylbenzidine (Sigma-Aldrich), 0.01% H2O2, 100 mM sodium acetate, 100 mM citric acid] and the reaction was stopped after 3.5 min by adding 25 μL 0.8 M H2SO4. Absorbance was measured at 450 nm. EC_50_ values were calculated using Graphpad Prism 9.5.1.

To assess binding of polyclonal rabbit or mouse sera to full-length E1E2, a Galanthus nivalis lectin ELISA was performed, similar to those described earlier with some alterations.^89^ High-Binding half-well 96-well plates were coated with 50 μL Galanthus nivalis lectin (Vector Laboratories) at 20 μg/mL in 0.1 M NaHCO3 pH 8.6. The next day, the plates were blocked with casein blocking buffer (Thermo Fisher Scientific) for 30 min. Afterwards the plates were washed twice with TBS and then 50 μL 1 μg/mL antigen was added to the lectin-coated plates. After 2 h, the plates were washed twice with TBS and incubated with 50 μL serially diluted serum in casein blocking buffer for 2 h. The plates were washed three times with TBS and 50 μL of a 1:3000 dilution of the animal-matched HRP-labeled goat anti-rabbit, or goat anti-mouse IgG (Jackson Immunoresearch) in casein blocking buffer was added for 1h. After washing the plates five times with TBS + 0.05% Tween-20, plates were developed by adding 50 μL develop solution [1% 3,3′,5,5′-tetraethylbenzidine (Sigma-Aldrich), 0.01% H2O2, 100 mM sodium acetate, 100 mM citric acid] and the reaction was stopped after 3.5 min by adding 25 μL 0.8 M H2SO4. Absorbance was measured at 450 nm. EC50 values were calculated using Graphpad Prism 9.5.1.

### Bio-layer interferometry (BLI)

BLI assays were performed as described previously.^44^ BLI assays were performed using an Octet K2 instrument (ForteBio). All assays were performed at 30 °C and with an agitation speed of 1000 rpm. mAbs, sE1E2.v3-LZ, sE1E2.v4-LZ, C2P2, C2P2-I53-50A trimer and C2P2-NP samples were dissolved in running buffer (PBS, 0.02% Tween, 0.1% bovine serum albumin (BSA)) in a volume of 250 μL/well. mAbs (1 μg/mL) were immobilized onto protein A biosensors (FortéBio, cat no. 18–5010) until a loading threshold of 1.0 nm was reached, followed by a 30 s baseline measurement in the running buffer. Purified proteins were diluted to 250 nM and association and dissociation (in a separate well containing only running buffer) were measured for 300 s each. A well containing running buffer without protein was used for background correction. Data was analyzed and visualized in GraphPad Prism 9.5.1.

### I53-50B.4PTI expression and purification

The I53-50B.4PTI component was expressed and purified as described before.^89,92^ Lemo21 (DE3) (NEB) cells expressing I53-50B.4PT1 were grown in a 10 L BioFlo 320 Fermenter (Eppendorf) or a 2 L shake flask. Cells were grown in LB (10 g Tryptone, 5 g Yeast Extract, 10 g NaCl) at 37 °C to an OD600 of 0.8. After the cells were induced with 1 mM of IPTG, temperature was reduced to 18 °C and cells were grown for 16 h. The cells were then lysed in 50 mM Tris, 500 mM NaCl, 30 mM imidazole, 1 mM PMSF, 0.75% CHAPS using a Microfluidics M110P at 18,000 psi. Lysate was centrifuged at 24,000 × g for 30 min. Clarified lysate was next applied to a Ni Sepharose 6 FF column (Cytiva) linked to an AKTA Avant150 FPLC system for immobilized metal affinity chromatography. I53-50B.4PT1 was eluted using a 30–500 mM imidazole linear gradient in 50 mM Tris pH 8, 500 mM NaCl and 0.75% CHAPS. Fractions containing I53-50B.4PT1 were pooled, concentrated using centrifugal filters with a 10,000 kDa cutoff (Millipore), sterilized and applied to a Superdex 200 Increase 10/300 (Cytiva) for further purification. Batches were tested to ensure low levels of endotoxin before use.

### Nanoparticle assembly

The C2P2-NPs were assembled as described before.^89^ To remove aggregated protein, C2P2-I53-50A proteins were passed through a Superdex 200 Increase 10/300 GL SEC column (GE Healthcare) in TBS/5% glycerol, pH 7.5. The column fractions containing non-aggregated C2P2-I53-50A trimers were pooled and mixed in an equimolar ratio with I53-50B.4PT1 (produced as described above) for an overnight (∼16 h) incubation at 4 °C. The assembly mix was then concentrated at 1000 × g using Vivaspin filters with a 10 kDa molecular weight cutoff (Sartorius) and passed through a Superose 6 Increase 10/300 GLcolumn in TBS/5% glycerol, pH 7.5. The fractions corresponding to the assembled NPs were pooled and concentrated at 500 × g using Vivaspin filters with a 10 kDa molecular weight cutoff. Nanoparticle concentrations were determined with a Nanodrop using the peptide molecular weight and extinction coefficient.

### Negative-stain EM

Negative-stain EM experiments were performed as described previously.^89^ C2P2-NP and free NP subunit samples were diluted to 20–50 μg/mL and loaded onto the carbon-coated 400-mesh Cu grid that had previously been glow-discharged at 15 mA for 25 s. Grids were negatively stained with 2% (w/v) uranyl formate for 60 s. Data collection was performed on a Tecnai Spirit electron microscope operating at 120 keV. The magnification was ×52,000 with a pixel size of 2.06 Å at the specimen plane. All imaging was performed with a defocus value of −1.50 μm. The micrographs were recorded on a FEI Eagle CCD (4k) camera using Leginon automated imaging interface. Data processing was performed in Appion data processing suite. With nanoparticle samples, particles were picked from the micrographs and 2D-classified using the Iterative multivariate statistical analysis (MSA)/multireference alignment (MRA) algorithm.

### Cryo-EM sample preparation and data collection

Cryo-EM sample preparation and data collection were performed as described previously.^50^ Purified sE1E2.v4-LZ and Fabs were mixed at 1:1 mass ratio overnight at 4 °C. The next day, samples were filtered and loaded on an S200 increase size-exclusion chromatography column in TBS. The complex peaks were isolated from unbound Fab fractions, concentrated to a working range of 1-3 mg/mL and 3.5 μL of the concentrated samples were loaded on a glow-discharged grid (Quantifoil 1.2/1.3, Cu 300), blotted using a Vitrobot Mark IV (ThermoFisher) at 4 °C and 100% humidity and flash-frozen in liquid ethane.

Imaging was done on a Titan Glacios II (ThermoFisher) equipped with a Falcon 4(i) direct electron detector (ThermoFisher). Data was collected at a nominal pixel size of 0.718 Å 2 / px and a total dose of 60 electron / Å 2. After screening for grid quality, automated data collection was set-up with EPU (ThermoFisher) to collect movies. Movies were processed in CryoSPARC Live^132^, aligned with Patch Motion Correction and corrected with Patch CTF, rejecting micrographs with an estimated CTF above 5 Å. Blob picker was used to select particles between 60 and 160 Å. Junk particles were removed by an initial round of 2D and selected particles were taken to 3D refinement. Rounds of ab-initio model building and 3D Non-Uniform refinement jobs ensued^133^, at which point selected unbinned particles were processed to the final stage. Global and local CTF refinement was applied to the unbinned batch of final particles. The final map was generated by local refinement using a mask created in USCF ChimeraX to hide the constant region of the antibody and focus on the variable domain and antigen density.

### Protein model building and refinement

Protein model building and refinement were performed as described previously.^50^ The sharpened map and the antibody sequences were provided to Modelangelo^134^ for an initial model building. The resulting model was then manually refined in Coot^135^ and then real-space refined with Phenix^136^. The model was relaxed using Rosetta and then refined iteratively through Coot and Phenix. Data collection, image processing and map and model refinement parameters are shown in Supplementary Table 1. Structural Figures were made using USCF ChimeraX.^126–128^

### Binding angles and azimuth calculations

For every indicated antibody, a plane was defined in ChimeraX using the command *define plane*, working only with the variable domain (VH, VL) of each antibody to account for models deposited without CH1/CL domains. An axis was defined, using the command *define axis,* as the axis between the center of mass of each variable domains and the AR3 center of mass. Azimuth then represents the angle of such axis (i.e. where does it point to) with respect to the AR3 plane normal axis. The Class-I HEPC74 mAb was arbitrarily defined as 0°, and every other mAb measured counter-clockwise relative to it. Inclination is the angle between the planes (i.e. how does the fab approach the antigen), with 0° for a parallel approach and 90° for a perpendicular approach. All plots were then generated with R-Studio.^137^

### B cell activation assay

AR3C-, HEPC74-, COVA 1-16-, igl-AR3A-, igl-AR3C-, igl-HEPC74-, igl-1416_01_E03-, igl-1382_01_H05-and igl-AT1618-carrying B cells were generated, as described previously.^89^ In short, the gl2-1261 gene of the pRRL EuB29 gl2-1261 IgG TM.BCR.GFP.WPRE plasmid107 was exchanged for the heavy and light chain genes of the aforementioned (igl-)bNAbs using Gibson assembly (Integrated DNA Technologies). Lentiviruses were produced by co-transfecting the expression plasmid with pMDL, pVSV-g, and pRSV-Rev into HEK293T cells using lipofectamine 2000 (Invitrogen). Two days post transfection, IgM-negative Ramos B cells were transduced with HEK293T supernatant. Seven days post transduction, the aforementioned (igl-)bNAb-expressing B cells were sorted using Fluorescence activated cell sorting (FACS) on IgG and GFP double-positivity using a FACS Aria-II SORP (BD Biosciences). B cells were expanded and cultured indefinitely.

B cell activation experiments of the aforementioned (igl-)bNAb -carrying Ramos B cells were performed as previously described.^80^ In short, 4 million cells/mL in RPMI++ (RPMI with 1x penicillin-streptomycin and 10% FCS) were loaded with 1.5 μM of the calcium indicator Indo-1 (Invitrogen) for 30 min at 37 °C, washed with Hank’s Balance Salt Solution supplemented with 2 mM CaCl_2_, followed by another incubation of 30 min at 37 °C. Antigen-induced Ca^2+^ influx of B cells were monitored on a LSR Fortessa (BD Biosciences) by measuring the 379/450 nm emission ratio of Indo1 fluorescence upon UV excitation. Following 30 s of baseline measurement, aliquots of 1 million cells/mL were then stimulated for 270 s at room temperature (RT) with 0.13µM amounts of C2P2 for C2P2, HepCon C2P2-I53-50A trimers and HepCon C2P2-NP. Ionomycin (Invitrogen) was added to a final concentration of 1 mg/mL to determine the maximum Indo-1-fluorescence. Equimolar amounts of SARSCoV-2 spike were used as negative control.^85^ Kinetics analyses were performed using FlowJo v8.1.

### Rabbit immunizations

The rabbit immunization was performed as described before.^50^ 18 rabbits (New Zealand White, female, 3 groups, 6 animals/group) were immunized under subcontract at Labcorp (Denver, USA. Study No. 0201-23) with either 18 μg AMS0232 sE1E2.v4-LZ, 18 μg HepCon sE1E2.v4-LZ or 4 x 4.5 μg H77, AMS3a, UKNP4.4.4 and AMS0232 sE1E2.v4-LZ. Antigens were mixed 1:1 with squalene o/w emulsion (SE) (250 μL of antigen in PBS combined with 250 μL SE) (Polymun, Klosterneuburg, Austria) and administered by two intramuscular immunizations in each quadriceps (2 × 250 μL) at weeks 0, 4 and 20. Rabbits were bled at weeks 0, 4, 6, 12, 20 and 22. All procedures in this study design are in compliance with the U.S. Department of Agriculture’s (USDA) Animal Welfare Act (9 CFR Parts 1, 2, and 3); the Guide for the Care and Use of Laboratory Animals (Institute of Laboratory Animal Resources, National Academy Press, Washington, D.C., 2011); and the National Institutes of Health, Office of Laboratory Animal Welfare. Whenever possible, procedures in this study are designed to avoid or minimize discomfort, distress, and pain to animals.

### HCVpp production

HCVpp generation and neutralization assays were performed as described elsewhere.^68^ One day prior to transfection for generating HCVpp, 1.5 × 10^6^ HEK-293T-CD81KO cells were seeded on a 10 cm^2^ dish. Cells were co-transfected with three plasmids: MLV Gag-Pol packaging construct, firefly luciferase and HCV E1E2 in optimized ratios^68,138^ with a total amount of 6 µg of DNA and 12 µL of Lipofectamine 2000 (Invitrogen) in Opti-MEM (ThermoFisher). After an incubation overnight, Opti-MEM was replaced by DMEM (Gibco)/10% FCS/1x penicillin-streptomycin/1x MEM-NEAA/10mM HEPES. Two days later, the supernatant containing the HCVpps was passed through a 0.45 µm filter and frozen at −80 °C for long-term storage or 4 °C when used within a week. Infectivity of the HCVpp was assessed as described previously^68^.

### Neutralization assays

The neutralization assays were performed as described before.^50^ Huh-7 cells, a gift from François-Loїc Cosset, a hepatocyte-derived carcinoma cell line, were seeded at 1.0 × 10^4^ cells per well in a 96-well plate 24 h prior to the experiment in 100 µL DMEM supplemented with 10% FCS, 1x penicillin/streptomycin, 1x MEM-NEAA and 10mM HEPES buffer (Huh-7 medium). HCVpps were incubated with serially diluted serum concentration in duplicate at 37 °C in 5% CO2. After 1 h, media was removed from Huh-7 cells and the HCVpp/serum mixture (30 μL) was added and incubated for 4 h. After the incubation, 220 µL of Huh-7 medium was added and incubated for 72 h. After removing the media, cells were lysed and luciferase signal was measured using the Luciferase Assay System (Promega) and a GloMax luminometer (Promega, USA). Data was analyzed and visualized in GraphPad Prism 9.5.1.

### sE1E2 coupling to Luminex beads

The coupling of sE1E2 proteins to the Magplex beads was performed similarly as described previously.^69^ Each purified soluble E1E2 was covalently coupled to one Magplex bead region (Luminex Corporation) using a two-step carbodiimide reaction. Briefly, - 0.47 µg of each soluble E1E2 was coupled to 0.47 × 10^6^ Magplex beads (Luminex). The soluble E1E2 proteins included H77 sE1E2.v4-LZ, UKNP4.1.1 sE1E2.v4-LZ, AMS3a sE1E2.v4-LZ, AMS0232 sE1E2.v4-LZ, an HepCon E1E2.v4. Magplex beads were washed with 100 mM monobasic sodium phosphate pH 6.2 and activated by the addition of Sulfo-N-Hydroxysulfosuccinimide (Thermo Fisher Scientific) and 1-Ethyl-3-(3-dimethylaminopropyl) carbodiimide (Thermo Fisher Scientific) for 30 min on a rotor at room temperature (RT) in the dark. Activated beads were washed three times with 50 mM MES (Thermo Fisher Scientific) pH 5.0 before the addition of soluble E1E2 diluted in 245 μL of 50 mM MES pH 5.0. The mix containing the beads and soluble E1E2 was incubated for 3 h on a rotor at room temperature in the dark before washing with PBS to elute any unbound protein. Subsequently, the beads were incubated with blocking buffer (PBS containing 2% BSA, 3% FBS, 0.02% Tween-20) for 30 min on a rotator at RT. Beads were then washed and stored with 0.05% Sodium Azide in PBS at pH 7.0. Prefusion stabilised trimeric RSV-fusion glycoprotein^139,140^ and empty beads were used as positive control and negative control, respectively.

### Competition Luminex assay

The binding of antibodies to the sE1E2 proteins coupled to the Magplex beads was studied similarly as described previously.^141^ Briefly, in 50 μl of blocking buffer (PBS containing 2% BSA, 3% FBS, 0.02% Tween-20), 750 coupled beads per region were incubated with 25 μL of competitors for 1 h in the dark. The competitors included rabbit serum diluted 1:500 and blocking buffer only. Subsequently, 25 μL of mAb (AP33 at a final concentration of 0.5 μg/ml, IGH505: 0.5 μg/mL, AR4A: 1 μg/mL, AT1618: 10 μg/mL, CD81-fc: 0.5 μg/mL, 1382_01_H05: 0.5 μg/mL, AT1211: 1 μg/mL or CBH-4B: 10 μg/mL) were added, bringing the total volume to 100 μL. Plates were then incubated overnight at 4 °C with rotation in the dark. The next day, plates were washed twice with TBS containing 0.05% Tween-20 (TBST) using a hand-held magnetic separator. Beads were resuspended in 50 μL of blocking buffer containing the detection antibody mouse-anti-human-IgG1-PE at a final concentration of 1.3 ng/mL. After 2 h of incubation at RT with rotation in the dark, the beads were washed twice with TBST using a hand-held magnetic separator. Finally, the beads were resuspended in 70 μL of Bioplex sheath fluid (Bio-Rad), and after a few minutes of rotation at RT, readouts were performed on the Bioplex 200 (Bio-Rad).

Resulting median fluorescence intensity (MFI) values were corrected by subtracting MFI values from buffer and beads-only wells before proceeding with the analysis. Next, we set any value above 10 times the highest median value (10 MFI) of the RSV/uncoupled beads as a threshold for genuine binding of the mAbs. We observed that some mAbs co-incubated with blocking buffer only produced low MFI values. We considered the maximum MFI of mAbs binding to beads coupled with RSV and uncoupled beads as 0% binding. The fold reduction in binding was calculated considering the non-competitor as 100%.

### In vitro transcription of mRNA

mRNA for the different HCV constructs was synthetized from linearized plasmids as a template via an in vitro transcription (IVT) reaction using the HiScribe kit (New England BioLabs), according to the manufacturer’s instructions. Transcripts were purified by lithium chloride precipitation, and mRNA pellets were resuspended in water and quantified by NanoDrop One (Thermo Fisher Scientific, UK). RNA quality was assessed by RNA gel electrophoresis and stored at −80 °C. Specifically for the first mice immunization experiment (Supplementary Fig. 10D) mRNA with N1-Methyl-Pseudouridine-5’-Triphosphate (N1-m-UTP) was used, where for the other mice immunization experiments the mRNA did not contain N1-m-UTP.

### mRNA formulation into LNPs

LNP formulations were prepared as previously described.^142^ Briefly, C12-200, 1,2-distearoyl-sn-glycero-3-phosphocholine (DSPC), cholesterol (plant-derived), and 1,2-dimyristoyl-sn-glycero-3-phosphoethanolamine-N-[methoxy(polyethylene glycol)-2000] (ammonium salt) (DMPE-PEG2000) were purchased from Avanti Polar Lipids (Merck), and stock solutions prepared in absolute ethanol. Lipids were mixed at molar ratios of 35:16:46.5:2.5 C12-200:DSPC:Cholesterol:DMPE-PEG2000 at a total lipid concentration of 15 mM. mRNA solutions were prepared in 50 mM sodium acetate (Sigma-Aldrich) and 100 mM sodium chloride buffer (Sigma-Aldrich), adjusted to pH 5.5. LNPs were prepared using an RNA:lipid ratio of 1:55 (w/w). LNPs were formulated using the NanoAssemblr™ Ignite system (Cytiva) at a flow rate of 8 mL/min. Formulated LNPs were diluted in Dulbecco’s phosphate-buffered saline (DPBS, Gibco) at a ratio of 1:5 (v/v) and subsequently concentrated using Amicon Ultra-15 centrifugal filters with a 10 kDa molecular weight cutoff (Merck). LNPs were stored at 4 °C for immediate use in 20 mM Tris-HCl buffer pH 7.5 or supplemented with 10% sucrose for long-term storage at −80 °C. The encapsulated RNA was stained using the Quant-iT RiboGreen RNA Assay Kit (Thermo Fisher Scientific) according to the manufacturer’s instructions and quantificatied based on fluorescence intensity measured with a FLUOstar® Omega plate reader (BMG Labtech) at excitation/emission wavelengths of 485/535 nm. LNP average size of 100-150 nm, polydispersity index lower or equal to 0.3, and zeta potential negative or close to neutral; all these parameters were determined using a Zetasizer Nano ZS (Malvern Instruments).

### In vivo immunogenicity studies

In the first study, BALB/c mice (n = 5) received doses of 5 µg of mRNA-LNPs for each antigen via intramuscular injection. The immunisations were administered at weeks 0 and 4. The study endpoint was at week 7 when animals were culled for terminal bleeding. Blood was spun down, serum collected and stored at −80 °C.

In the second study, BALB/c mice (n = 5) received doses of 5 µg of mRNA-LNPs for each antigen via intramuscular injection. The immunisations were administered at weeks 0 and 4. The study endpoint was at week 6 when animals were culled for terminal bleeding. Blood was spun down, serum collected and stored at −80°C. Spleens from week 6 were harvested and processed for obtention of splenocytes.

In the third study, BALB/c mice (n = 5) received doses of 5 µg of mRNA-LNPs for each antigen on weeks 0 and 4 and a dose of 1µg mRNA-LNPs for each antigen on week 10 via intramuscular injection. Blood was collected at weeks 0, 4 and 10 before injections. The study endpoint was at week 12 when animals were culled for terminal bleeding. Blood was spun down, serum collected and stored at −80°C.

### ELISPOTS

Production of IFNγ by splenocytes in ELISPOTS was assessed using precoated 96 well plates (Mabtech) and the assays were performed according to manufacturer’s instructions. In brief, 2.5×10⁵ of fresh splenocytes were seeded per well in complete RPMI (supplemented with 10% FCS and 1x penicillin-streptomycin). Splenocytes were stimulated with 5µg of antigen, negative controls (RPMI only) and positive controls (RPMI plus cell Stimulation Cocktail 1:1000). Cells were incubated for 24h at 37 °C in an incubator with 5% CO_2_. After, plates were washed with PBS and incubated with IFNg antibody, followed by incubation with streptavidin-ALP antibody. Spots were developed using BCIP/NBT as substrate, then plates were rinsed with tap water, air-dried, and quantified using an automated ELISpot reader AID iSpot (Autoimmun Diagnostika GMBH).

## Data availability

The sE1E2.v4-LZ-AT1211-AT1209, sE1E2.v4-LZ-AT1211-1416_01_E03 and sE1E2.v4-LZ-AT1211-igl-1416_01_E03 models have been deposited into the RSCB PDB (https://www.rcsb.org) under accession numbers 37M0, 37MM and 37ML, respectively, and the corresponding electron microscopy data at the EMDB database (https://www.ebi.ac.uk/emdb/) under accession number EMD-78312, EMD-78310 and EMD-78309. Mass spectrometry RAW files have been deposited on the MassIVE server (https://massive.ucsd.edu) under accession number MSV000102230.

## Acknowledgements

We acknowledge the Scripps Research Institute CryoEM Facility and additional scientific resources at the Scripps Research Institute. We thank Rashmi Ravichandran and Neil P King for providing the I53-50B.4PT1 protein. We thank Tim Beaumont, Sabrina Merat, Ana Chumbe and Suzanne van den Aardweg for providing sequences and data on AT1211. We thank James E Crowe and Justin Bailey for providing the HEPC111, HEPC108, HEPC146 and HEPC167 antibodies. We also thank Marit van Gils for scientific discussions and valuable input. Finally, we thank Marlon de Gast for the *in vitro* B cell activation assay protocol.

## Author contributions

Conceptualization: F.M., J.C.P., F.C., S.P., R.W.S., and K.S. Funding acquisition: K.S., R.W.S., J.S, M.C, and R.J.S. Investigation: F.M., J.C.P., F.C., S.P., M.L.N., M.P., W.O., S.v.d.P., W.L., L.G., M.B.O., K.P., R.W., L.R. and I.Z. Methodology: F.M., J.C.P., F.C., L.R., M.L.N., M.C, R.J.S., R.W.S., and K.S. Resources: T. W. and F. K. Project administration: F.M., J.C.P., F.C., R.W.S., and K.S. Supervision: J.S., R.W.S., A.W., R. J. S., M.C. and K.S. Writing—original draft: F.M., J.C.P., F.C., R.W.S., and K.S. Writing—review & editing: all authors

