## Supplementary Information for "Consensus native-like hepatitis C virus E1E2 engages broadly neutralizing antibody precursors"

***Supplementary Table 1. Data collection, image processing and map and model refinement parameters.***

| | **Hepcon sE1E2.v4 AT1211 – AT1209** (EMDB-78312) (PDB 37MO) | **Hepcon sE1E2.v4 AT1211 – 1416_01_E03** (EMDB-78310) (PDB 37MM) | **Hepcon sE1E2.v4 AT1211 – igl-1416_01_E03** (EMDB-78309) (PDB 37ML) |
|---|---|---|---|
| **Data collection and processing** | | | |
| Magnification | **190x** | **190x** | **190x** |
| Voltage (kV) | **200** | **200** | **200** |
| Electron exposure (e–/Å$^2$) | **60** | **60** | **60** |
| Defocus range (μm) | **-0.8 to -1.8** | **-0.8 to -1.8** | **-0.8 to -1.8** |
| Pixel size (Å) | **0.718** | **0.718** | **0.718** |
| Symmetry imposed | **C1** | **C1** | **C1** |
| Initial particle images (no.) | 1 358 882 | 1 472 808 | 5 696 316 |
| Final particle images (no.) | 178 433 | 104 505 | 173 832 |
| Map resolution (Å) | 3.33 | 3.4 | 3.5 |
| FSC threshold | 0.143 | 0.143 | 0.143 |
| **Refinement** | | | |
| Map sharpening *B* factor (Å$^2$) | -114.9 | -80.2 | -106.9 |
| Model composition | | | |
| Non-hydrogen atoms | 5537 | 5392 | 5410 |
| Protein residues | 700 | 682 | 685 |
| Ligands | 15 | 13 | 13 |
| *B* factors (Å$^2$) | | | |
| Protein | 77.27 | 86.57 | 73.26 |
| Ligand | 99.36 | 98.81 | 74.36 |
| R.m.s. deviations | | | |
| Bond lengths (Å) | 0.004 | 0.005 | 0.004 |
| Bond angles (°) | 0.730 | 0.875 | 0.871 |
| Validation | | | |
| MolProbity score | 1.7 | 1.78 | 1.66 |
| Clashscore | 4.81 | 6.27 | 4.17 |
| Poor rotamers (%) | 0 | 0 | 0 |
| Ramachandran plot | | | |
| Favored (%) | 92.9 | 93.3 | 92.74 |
| Allowed (%) | 7.1 | 6.7 | 7.26 |
| Disallowed (%) | 0 | 0 | 0 |

A

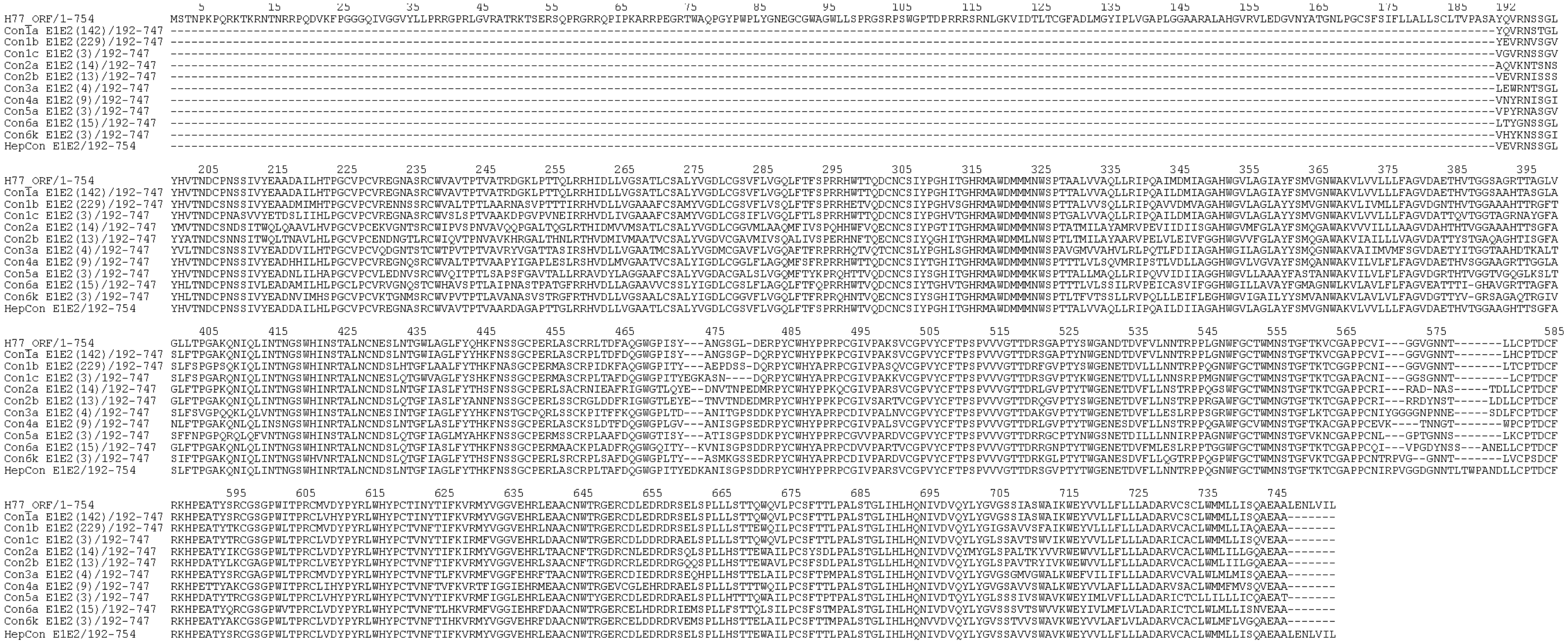
H77_ORF/1-754
Con1a E1E2(142)/192-747
Con1b E1E2(229)/192-747
Con1c E1E2(3)/192-747
Con2a E1E2(14)/192-747
Con2b E1E2(13)/192-747
Con3a E1E2(4)/192-747
Con4a E1E2(9)/192-747
Con5a E1E2(3)/192-747
Con6a E1E2(15)/192-747
Con6k E1E2(3)/192-747
HepCon E1E2/192-754

B

HepCon sE1E2.v3-LZ/192-714
VEVRNSSGLYHVTNDCPNSSIVYEADDAILHLPGCVPCVREGNASRCWVAVTPTVAARDAGAPTTGLRRGGSGGSGGSPRRHWTVQDCNCSIYPGHITGHRMAWDMMMNPSPTTALVVAQLLRIPQAILDIIAPGGRIARLEEKVKTLKAQNSELASTANMLREQVAQLKQKVMNYRRRRRRETHVTGGAAGHTTS
GFASLFTPGAKQNIQLINTNGSWHINRTALNCNDSLNTGFIAGLFYTHKFNSSGCPERLASCRPLTAFDQGWGPITYANISGPSDDRPYCWHYPPRPCGIVPARSVCGPVYCFTPSPVVVGTTDRSGVPTYTWGENETDVFLLNNTRPPQGNWFGCTWMNSTGFTKTCGAPPCNIGGDGNNTLTCPTDCFRKHPEA
TYSRCGSGPWLTPRCLVDYPYRLWHYPCTANFTIFKVRMYVGGVEHRLEAACNWTRGERCDLEDRDRSELSPLLHSTTEWAILPCSFTTPPALSTGLIHLHQNIVDVQYLYGVSSAVVSWAVPGGLTDTLQAETDQLEDKKSALQTEIANLLKEKEKLEFILAAYGGSAWSHPQFEKGGGSGGGSGGSSAWSHPQF
EK

HepCon sE1E2.v4-LZ/192-714
VEVRNSSGLYHVTNDCPNSSIVYEADDAILHLPGCVPCVREGNASRCWVAVTCTVAARDAGAPTTGLRRGGSGGSGGSPRRHWTVQDCNCSIYPGHITGHRMAWDMMMNPSPTTALVVAQLLRIPQAILDIIAPGGRIARLEEKVKTLKAQNSELASTANMLREQVAQLKQKVMNYRRRRRRETHVTGGAAGHTTS
GFASLFTPGAKQNIQLINTNGSWHINRTALNCNDSLNTGFIAGLFYTHKFNSSGCPERLASCRPLTAFDQGWGPITYANISGPSDDRPYCWHYPPRPCGIVPARSVCGPVYCFTPSPVVVGTTDRSGVPTYTWGENETDVFLLNNTRPPQGNWFGCTWMNSTGFTKTCGAPPCNIGGDGNNTLTCPTDCFRKHPEA
TYSRCGSGPWLTPRCLVDYPYRLWHYPCTANFTIFKVRMYVGGVEHRLEAACNWTRGERCDLEDRDRSELSPLLHSTTEWAILPCSFTTPPALSCGLIHLHQNIVDVQYLYGVSSAVVSWAVPGGLTDTLQAETDQLEDKKSALQTEIANLLKEKEKLEFILAAYGGSAWSHPQFEKGGGSGGGSGGSSAWSHPQF
EK

HepCon C2P2/192-714
VEVRNSSGLYHVTNDCPNSSIVYEADDAILHLPGCVPCVREGNASRCWVAVTCTVAARDAGAPTTGLRRGGSGGSGGSPRRHWTVQDCNCSIYPGHITGHRMAWDMMMNPSPTTALVVAQLLRIPQAILDIIAPGGRRRRRRETHVTGGAAGHTTSGFASLFTPGAKQNIQLINTNGSWHINRTALNCNDSLNTGF
IAGLFYTHKFNSSGCPERLASCRPLTAFDQGWGPITYANISGPSDDRPYCWHYPPRPCGIVPARSVCGPVYCFTPSPVVVGTTDRSGVPTYTWGENETDVFLLNNTRPPQGNWFGCTWMNSTGFTKTCGAPPCNIGGDGNNTLTCPTDCFRKHPEATYSRCGSGPWLTPRCLVDYPYRLWHYPCTANFTIFKVRMY
VGGVEHRLEAACNWTRGERCDLEDRDRSELSPLLHSTTEWAILPCSFTTPPALSCGLIHLHQNIVDVQYLYGVSSAVVSWAVPGGSAWSHPQFEKGGGSGGGSGGSSAWSHPQFEK

HepCon C2P2-I53-50A/192-714
VEVRNSSGLYHVTNDCPNSSIVYEADDAILHLPGCVPCVREGNASRCWVAVTCTVAARDAGAPTTGLRRGGSGGSGGSPRRHWTVQDCNCSIYPGHITGHRMAWDMMMNPSPTTALVVAQLLRIPQAILDIIAPGGRRRRRRETHVTGGAAGHTTSGFASLFTPGAKQNIQLINTNGSWHINRTALNCNDSLNTGF
IAGLFYTHKFNSSGCPERLASCRPLTAFDQGWGPITYANISGPSDDRPYCWHYPPRPCGIVPARSVCGPVYCFTPSPVVVGTTDRSGVPTYTWGENETDVFLLNNTRPPQGNWFGCTWMNSTGFTKTCGAPPCNIGGDGNNTLTCPTDCFRKHPEATYSRCGSGPWLTPRCLVDYPYRLWHYPCTANFTIFKVRMY
VGGVEHRLEAACNWTRGERCDLEDRDRSELSPLLHSTTEWAILPCSFTTPPALSCGLIHLHQNIVDVQYLYGVSSAVVSWAVPGGSGGSGGSGGSGGSEKAAKAEEAARKMEELFKKHKIVAVLRANSVEEAIEKAVAVFAGGVHLIEITFTVPDADTVIKALSVLKEKGAIIGAGTVTSVEQCRKAVESGAEFIV
SPHLDEEISQFCKEKGVFYMPGVMTPTELVKAMKLGHDILKLFPGEVVGPEFVKAMKGPFPNVKFVPTGGVDLDNVCEWFDAGVLAVGVGDALVEGDPDEVREKAKEFVEKIRGCTEGSLEWSHPQFEK

AMS0232 C2P2-
YQVRNSTGLYHVTNDCPNSSIVYETADAILHTPGCVPCVREGNASRCWVPMTCTVATRDGKLPATQLRRGGSGGSGGSPRRHWTTQDCNCSIYPGHVTGHRMAWDMMMNPSPTTALVVAQLLRIPQAILDMIAPGGRRRRRRQTYVTGGTAARATSGLANFFSPGAKQDVQLINTNGSWHINRTALNCNTSLETGW
IAGLIYLNKFNSSGCPERMASCRPLADFAQGWGPISYANGSGPDHRPYCWHYPPKPCGIVSAKSVCGPVYCFTPSPVVVGTTNKLGAPTYSWGENETDVFVLNNTRPPLGNWFGCTWMNSTGFTKVCGAPPCAIGGVGNNTLHCPTDCFRKHPEATYSRCGSGPWITPRCLVDYPYRLWHYPCTANYTRFKVRMYI
GGVEHRLDAACNWTRGERCDLEDRDRSELSPLLLSTTQWQVLPCSFTTPPALSCGLIHLHQNIVDVQYLYGVGSSIVSWAI

-Native E2 TMD
KWEYVVLLFLLLADARVCSCLWMMLLISQAEA
-SARS-COV-2 TMD
PGGSGGSYEQYIKWPWYIWLGFIAGLIAIVMVTIMLCCMTSCCSCLKGCCSCGSCC
-INFLUENZA HA TMD
PGGSGGSGVKLESMGIYQILAIYSTVASSLVLLVSLGAISFWMCSNGSLQCRICI
-VSVG TMD
PGGSGGSFCFIIGLIIGLFLVLRVGIYLCIKLKHTKKRQIYTDIEMNRLGK
-MARVG TMD
PGGSGGSWGVLTNLGILLLLSIAVLIALSCICRIFTKYIG
-ZEBOVG TMD
PGGSGGSWIPAGIGVTGVIIAVIALFCICKFVF
-BACVgp64 TMD
PGGSGGSSFMFGHVVNFVIILIVILFLYCMIRNRNRQY
-INTAIIB TMD
PGGSGGSALEERAIPIWWVLVGVLGGLLLLTILVLAMWKVGFFKRNR
-B7-1 TMD
PGGSGGSTLVLFGAGFGAVITVVVIVVIIKCFCKHRSCFRRNEASRETNNSLTFGPEEALAEQTVFL
-PDGFR TMD
PGGSGGSAVGQDTQEVIVVPHSLPFKVVVISAILALVVLTIISLIILIMLWQKKPR
-FasL TMD
PGGSGGSKKRGNHSTGLCLLVMFFMVLVALVGLGLGMFQLFHLQKETG

AMS0232 C2P2-TMD
YQVRNSTGLYHVTNDCPNSSIVYETADAILHTPGCVPCVREGNASRCWVPMTCTVATRDGKLPATQLRRGGSGGSGGSPRRHWTTQDCNCSIYPGHVTGHRMAWDMMMNPSPTTALVVAQLLRIPQAILDMIAPGGRRRRRRQTYVTGGTAARATSGLANFFSPGAKQDVQLINTNGSWHINRTALNCNTSLETGW
IAGLIYLNKFNSSGCPERMASCRPLADFAQGWGPISYANGSGPDHRPYCWHYPPKPCGIVSAKSVCGPVYCFTPSPVVVGTTNKLGAPTYSWGENETDVFVLNNTRPPLGNWFGCTWMNSTGFTKVCGAPPCAIGGVGNNTLHCPTDCFRKHPEATYSRCGSGPWITPRCLVDYPYRLWHYPCTANYTRFKVRMYI
GGVEHRLDAACNWTRGERCDLEDRDRSELSPLLLSTTQWQVLPCSFTTPPALSCGLIHLHQNIVDVQYLYGVGSSIVSWAIPGGSGGSWYIWLGFIAGLIAIVMVTIMLCCMTSCCSCLKGCCSCGSCC

AMS0232 C2P2-TMDdCT
YQVRNSTGLYHVTNDCPNSSIVYETADAILHTPGCVPCVREGNASRCWVPMTCTVATRDGKLPATQLRRGGSGGSGGSPRRHWTTQDCNCSIYPGHVTGHRMAWDMMMNPSPTTALVVAQLLRIPQAILDMIAPGGRRRRRRQTYVTGGTAARATSGLANFFSPGAKQDVQLINTNGSWHINRTALNCNTSLETGW
IAGLIYLNKFNSSGCPERMASCRPLADFAQGWGPISYANGSGPDHRPYCWHYPPKPCGIVSAKSVCGPVYCFTPSPVVVGTTNKLGAPTYSWGENETDVFVLNNTRPPLGNWFGCTWMNSTGFTKVCGAPPCAIGGVGNNTLHCPTDCFRKHPEATYSRCGSGPWITPRCLVDYPYRLWHYPCTANYTRFKVRMYI
GGVEHRLDAACNWTRGERCDLEDRDRSELSPLLLSTTQWQVLPCSFTTPPALSCGLIHLHQNIVDVQYLYGVGSSIVSWAIPGGSGGSWYIWLGFIAGLIAIVMVTIML

HepCon C2P2-TMD
VEVRNSSGLYHVTNDCPNSSIVYEADDAILHLPGCVPCVREGNASRCWVAVTCTVAARDAGAPTTGLRRGGSGGSGGSPRRHWTVQDCNCSIYPGHITGHRMAWDMMMNPSPTTALVVAQLLRIPQAILDIIAPGGRRRRRRETHVTGGAAGHTTSGFASLFTPGAKQNIQLINTNGSWHINRTALNCNDSLNTGF
IAGLFYTHKFNSSGCPERLASCRPLTAFDQGWGPITYANISGPSDDRPYCWHYPPRPCGIVPARSVCGPVYCFTPSPVVVGTTDRSGVPTYTWGENETDVFLLNNTRPPQGNWFGCTWMNSTGFTKTCGAPPCNIGGDGNNTLTCPTDCFRKHPEATYSRCGSGPWLTPRCLVDYPYRLWHYPCTANFTIFKVRMY
VGGVEHRLEAACNWTRGERCDLEDRDRSELSPLLHSTTEWAILPCSFTTPPALSCGLIHLHQNIVDVQYLYGVSSAVVSWAVPGGSGGSWYIWLGFIAGLIAIVMVTIMLCCMTSCCSCLKGCCSCGSCC

HepCon C2P2-TMDdCT
VEVRNSSGLYHVTNDCPNSSIVYEADDAILHLPGCVPCVREGNASRCWVAVTCTVAARDAGAPTTGLRRGGSGGSGGSPRRHWTVQDCNCSIYPGHITGHRMAWDMMMNPSPTTALVVAQLLRIPQAILDIIAPGGRRRRRRETHVTGGAAGHTTSGFASLFTPGAKQNIQLINTNGSWHINRTALNCNDSLNTGF
IAGLFYTHKFNSSGCPERLASCRPLTAFDQGWGPITYANISGPSDDRPYCWHYPPRPCGIVPARSVCGPVYCFTPSPVVVGTTDRSGVPTYTWGENETDVFLLNNTRPPQGNWFGCTWMNSTGFTKTCGAPPCNIGGDGNNTLTCPTDCFRKHPEATYSRCGSGPWLTPRCLVDYPYRLWHYPCTANFTIFKVRMY
VGGVEHRLEAACNWTRGERCDLEDRDRSELSPLLHSTTEWAILPCSFTTPPALSCGLIHLHQNIVDVQYLYGVSSAVVSWAVPGGSGGSWYIWLGFIAGLIAIVMVTIML

***Supplementary Figure 1: Construct overview and positional amino acid rarity scores. A*** *The E1E2 sequences of con1a, -1b, -1c, -2a, -2b, -3a, -4a, -5a, -6a, -6k and HepCon aligned to the reference H77 sequence (GenBank ID: AF009606). Sequences were aligned using Clustal Omega webtool.*[119] ***B*** *E1E2 sequences of HepCon sE1E2.v3-LZ, sE1E2.v4-LZ, C2P2, C2P2-I53-50A, and AMS0232 and HepCon C2P2 coupled to several TMDs.*

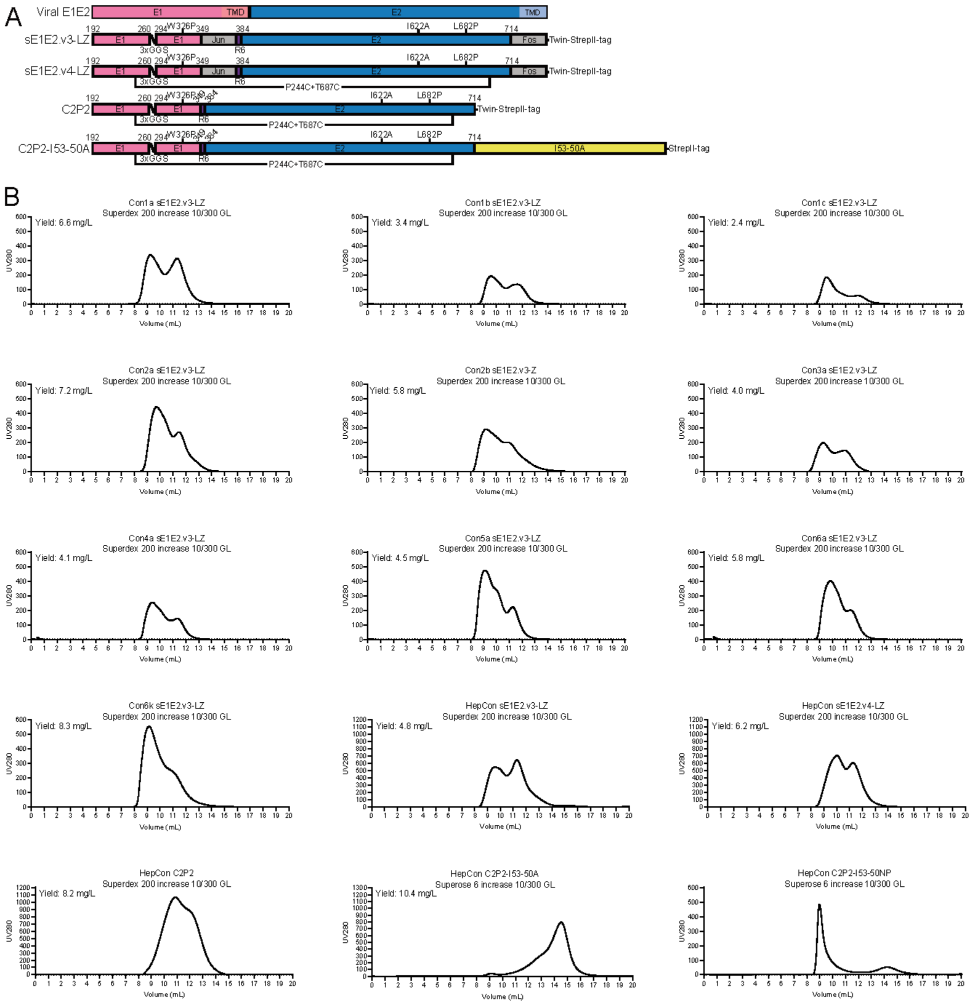


***Supplementary Figure 2: Protein design and SEC characterization A** Linear representation of viral E1E2 and sE1E2.v3-LZ, sE1E2.v4-LZ, C2P2 and C2P2-I53-50A designs with the transmembrane domain (TMD), dimerization domains (Jun, Fos), I53-50A domain, Twin-StrepII-tag, linkers and mutations indicated, as previously described.[50] **B** All SEC profiles were obtained using a Superdex 200 Increase 10/300 GL SEC column, except for HepCon C2P2-I53-50A and C2P2-I53-50NP which were passed through a Superose 6 Increase 10/300 GL column. Total yields before SEC are indicated.*

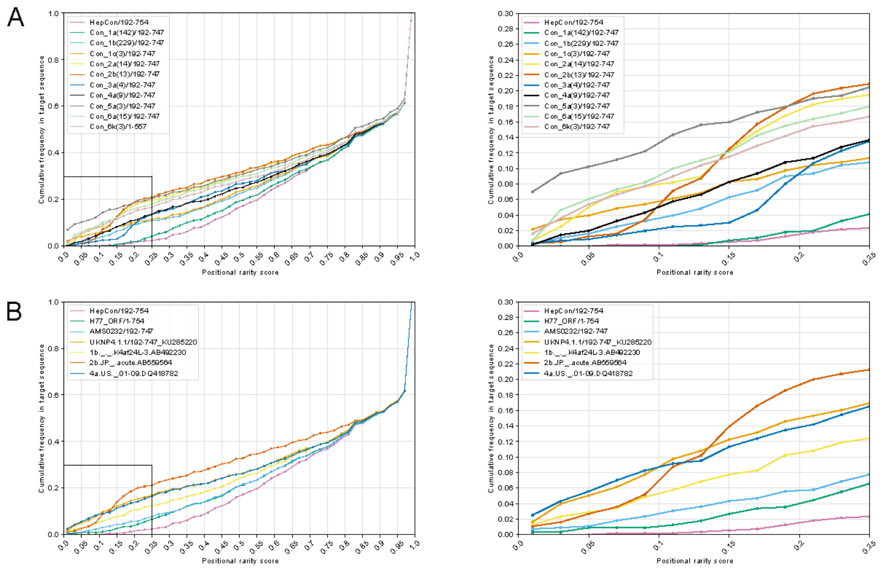


***Supplementary Figure 3 Residue rarity analysis. A,B*** *Residue rarity analysis for the concatenated E1E2 protein was performed using a multiple sequence alignment generated using MAFFT v7.526*[120] *of around 15000 E1E2 protein sequences from the Los Alamos National Laboratory HCV sequence database.*[53] *For each position in the alignment, the 'positional rarity' in sequences of interest were calculated as the proportion of non-gap sequences in the alignment having the same residue as the sequence of interest. A per-genotype weighting factor was included in this calculation to counteract the bias induced by the vastly differing numbers of sequences per genotype in the database. These positional rarity scores were used to produce the positional rarity histogram, of which the cumulative frequency in target sequence is depicted, with the full histogram (left) and partial histogram depicting uncommon (<25%) frequencies (right). Custom Python scripts used for these analyses are available at https://gitlab.com/ruben.walen/hepcon-germline-targeting.*

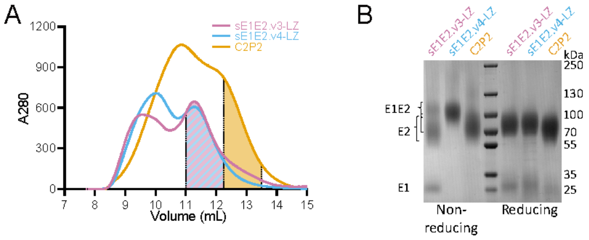


***Supplementary Figure 4: HepCon sE1E2 characterization. A*** *SEC profiles of HepCon sE1E2.v3-LZ, sE1E2.v4-LZ and C2P2 purified with Strep-TactinXT-based purification from 500 mL cultures of HEK293F cells on a Superdex 200 Increase 10/300 GL column. Pooled heterodimer fractions are indicated for each construct.* ***B*** *Non-reducing (left) and reducing (right) SDS-PAGE gels of HepCon sE1E2.v3-LZ, sE1E2.v4-LZ and C2P2. Bands corresponding to E1, E2 and E1E2 (± Jun and/or Fos) are indicated.*

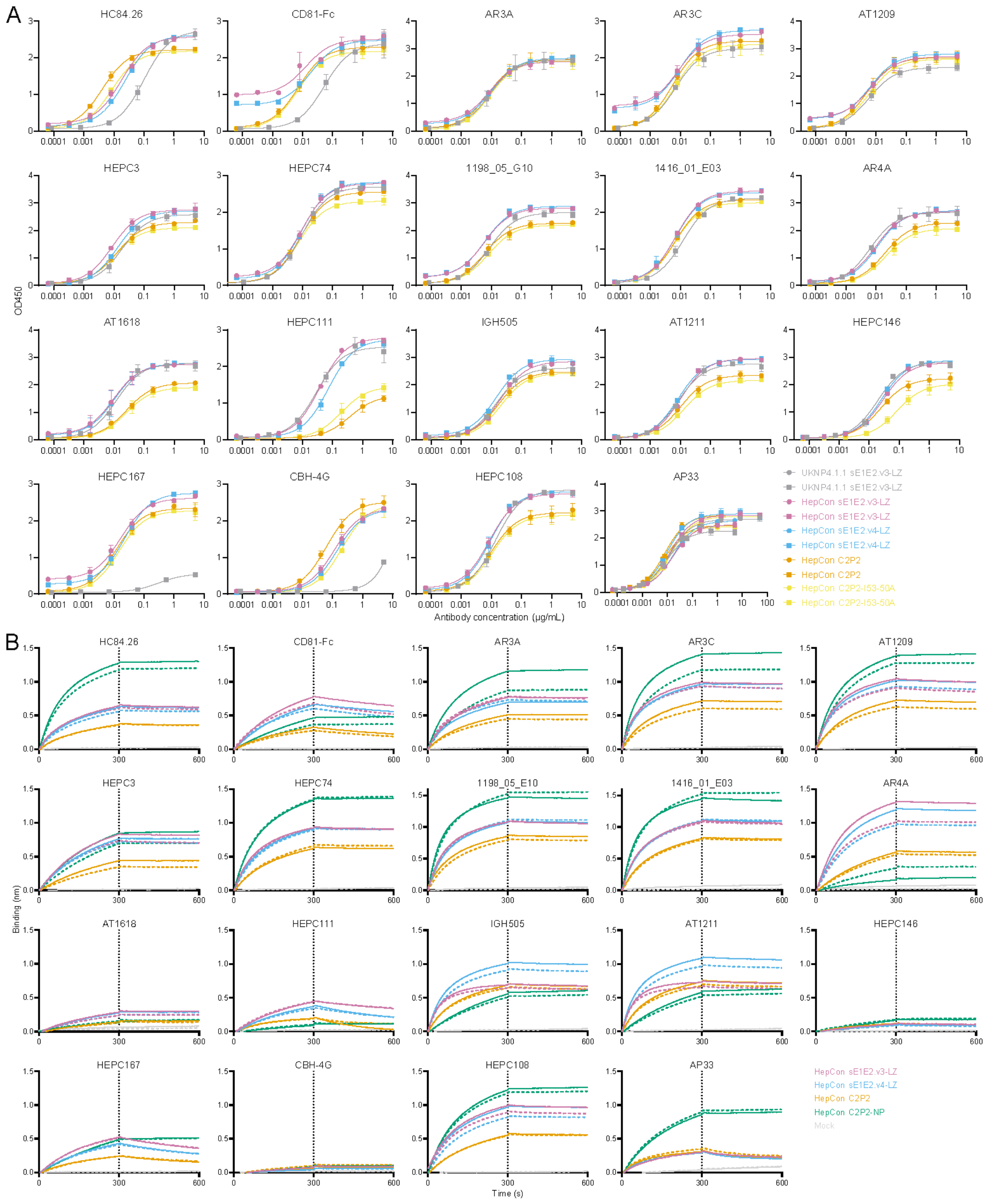


***Supplementary Figure 5: ELISA and BLI binding of Abs targeting different epitopes on E1 and E2. A*** *ELISA binding curves for HepCon sE1E2.v3-LZ, sE1E2.v4-LZ, C2P2 and C2P2-I53-50A were obtained using Strep-TactinXT ELISA with UKNP4.1.1 sE1E2.v3-LZ tested as comparator.*[50] *AP33 binding was determined as loading control. Results obtained in different experiments are indicated by different symbols. Means and SD are shown.* ***B*** *Binding profiles for HepCon sE1E2.v3-LZ, sE1E2.v4-LZ, C2P2 and C2P2-NP with the Abs were obtained using BLI. The indicated Abs were immobilized onto protein A biosensors and incubated for 300 seconds with 250 nM sE1E2.v3-LZ, sE1E2.v4-LZ, C2P2 or C2P2-NP and subsequently incubated for 300 seconds with running buffer. Replicates are shown in corresponding colors, as full and dotted lines. All ELISA and BLI analyses were performed in duplo.*

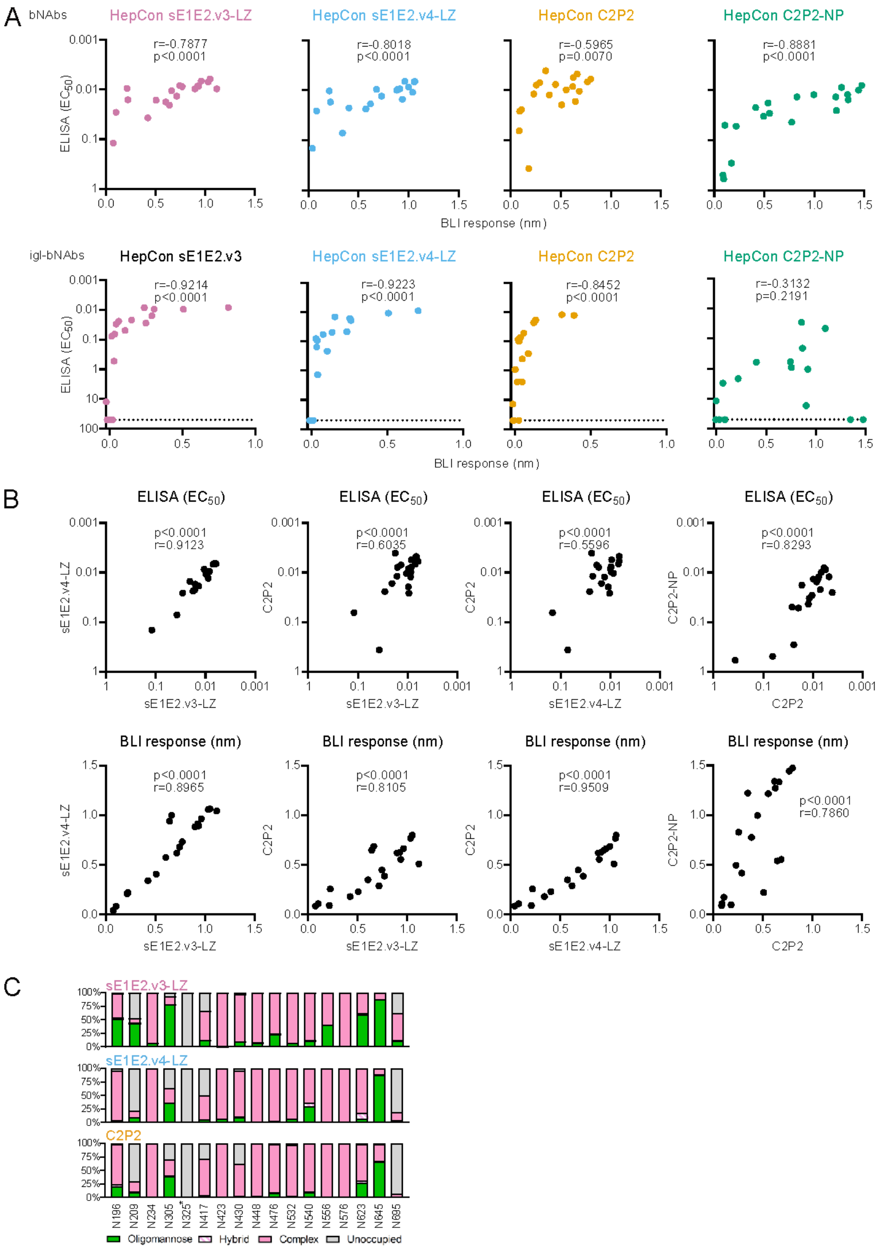


***Supplementary Figure 6: Correlations between BLI and ELISA and between sE1E2 constructs. A** Correlations between maximum BLI response and ELISA $EC_{50}$-values are shown for HepCon sE1E2.v3-LZ, sE1E2.v4-LZ, C2P2 and C2P2-NP for all tested bNAbs (top) and igl-bnAbs (bottom). **B** Correlations of ELISA $EC_{50}$-values (top) and BLI response (bottom) between the different HepCon sE1E2 constructs. Spearman r and p-values (two-tailed) are indicated in **A** and **B**. **C** Relative abundance of distinct glycan types obtained with quantitative site-specific glycan analysis of HepCon sE1E2.v3-LZ, sE1E2.v4-LZ and C2P2. Glycan compositions are grouped into their corresponding categories, with complex-type glycans displayed in pink, hybrid as hatched pink and white and oligomannose in green. *N325 is never glycosylated. ELISA and BLI values plotted in **A** and **B** are mean values obtained from Supplementary Figures 5A,B, 12A,B, 14A,B.*

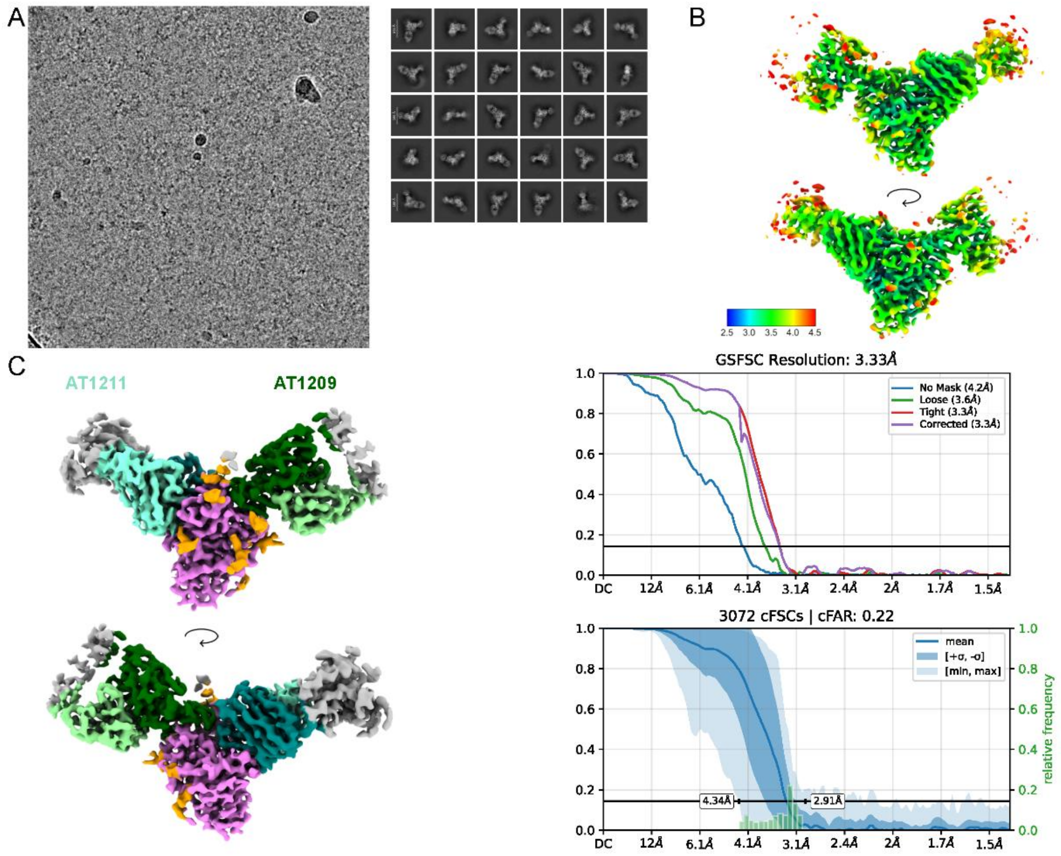


D

| | | Model | | |
|---|---|---|---|---|
| | | HepCon sE1E2.v4-LZ - AT1211 - AT1209 | HepCon sE1E2.v4-LZ - AT1211 - 1416_01_E03 | HepCon sE1E2.v4 - LZ- AT1211 - igl- 1416_01_E03 |
| ASNposition | N423 | NAG | NAG-NAG | NAG-NAG |
| | N430 | NAG-NAG | NAG | NAG |
| | N448 | NAG-NAG | NAG-NAG | NAG-NAG |
| | N532 | NAG | NAG | NAG |
| | N540 | NAG-NAG | NAG-NAG | NAG-NAG |
| | N556 | NAG-NAG-BMA | NAG-NAG-BMA | NAG-NAG-BMA |
| | N623 | NAG | NAG | NAG |
| | N645 | NAG-NAG-BMA | NAG | NAG |

E

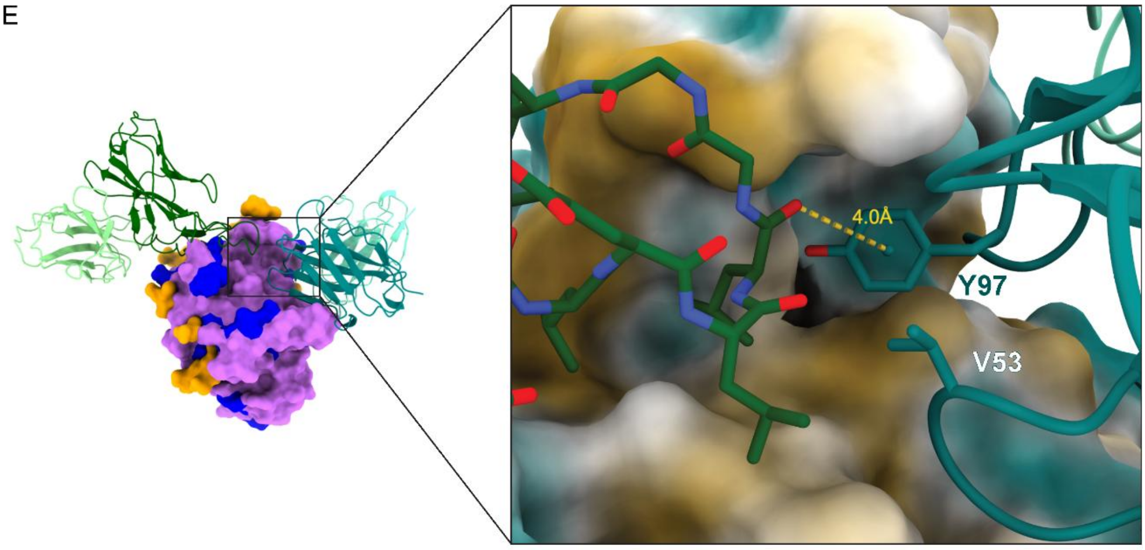


***Supplementary Figure 7: Cryo-EM of HepCon sE1E2.v4-LZ in complex with bNAb AT1211 and AT1209.*** ***A*** *Representative micrographs and 2D classes of AT1211-AT1209-E1E2 complexes.* ***B*** *Local resolution estimates of the complex, with resolution steps indicated in the key.* ***C*** *Cryo-EM map of AT1211-AT1209-*

*E1E2 and corresponding Fourier Shell Correlation (FSC). Color code on the map follows the color code on Fig.1E.* ***D*** *Table depicting glycans per PNGS for each of the three cryo-EM models described in Fig. 1E and Fig. 4.* ***E*** *Heterotypic contacts between AT1211 (teal) and AT1209 (green), with E2 represented as a hydrophobic surface.*

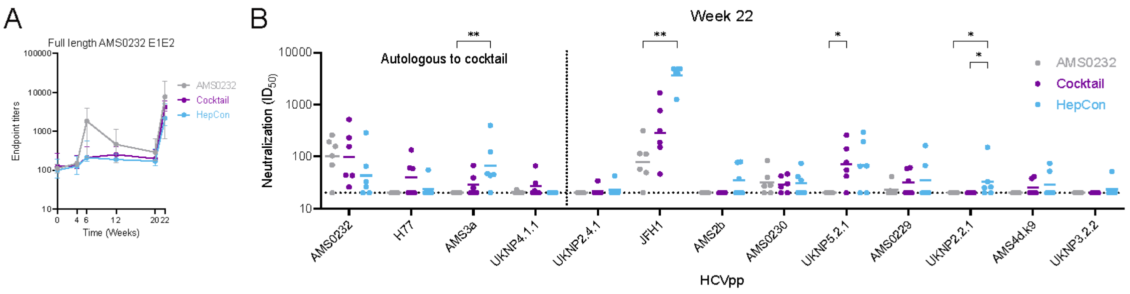


C

| HCVpp | | AMS0232 | | | | | | Cocktail | | | | | | HepCon | | | | | |
|---|---|---|---|---|---|---|---|---|---|---|---|---|---|---|---|---|---|---|---|
| Genotype | Rabbit | UA269 | UA270 | UA271 | UA272 | UA273 | UA274 | UA287 | UA288 | UA289 | UA290 | UA291 | UA292 | UA281 | UA282 | UA283 | UA284 | UA285 | UA286 |
| 1 | AMS0232 | 103 | 259 | 191 | 157 | 69 | 20 | 26 | 155 | 44 | 230 | 517 | 43 | 20 | 20 | 33 | 65 | 287 | 26 |
| 1 | H77 | 20 | 20 | 20 | 20 | 20 | 20 | 60 | 20 | 20 | 58 | 20 | 134 | 20 | 20 | 20 | 20 | 55 | 20 |
| 3 | AMS3a | 20 | 20 | 20 | 20 | 20 | 20 | 20 | 25 | 20 | 67 | 20 | 39 | 45 | 47 | 43 | 124 | 396 | 20 |
| 4 | UKNP4.1.1 | 23 | 20 | 20 | 20 | 20 | 20 | 20 | 20 | 31 | 20 | 66 | 22 | 20 | 20 | 20 | 20 | 20 | 20 |
| 2 | UKNP2.4.1 | 20 | 20 | 20 | 20 | 20 | 20 | 20 | 20 | 20 | 34 | 20 | 20 | 20 | 20 | 20 | 20 | 20 | 42 |
| 2 | JFH1 | 111 | 49 | 114 | 313 | 57 | 20 | 46 | 765 | 151 | 1687 | 316 | 187 | 4183 | 4860 | 3860 | 4860 | 4860 | 1264 |
| 2 | AMS2b | 20 | 20 | 20 | 20 | 20 | 20 | 20 | 20 | 20 | 20 | 20 | 20 | 20 | 20 | 20 | 37 | 79 | 77 |
| 1 | AMS0230 | 28 | 40 | 20 | 27 | 84 | 20 | 42 | 26 | 46 | 20 | 30 | 20 | 74 | 20 | 20 | 20 | 41 | 35 |
| 5 | UKNP5.2.1 | 20 | 20 | 20 | 20 | 20 | 20 | 42 | 55 | 141 | 258 | 77 | 20 | 65 | 20 | 294 | 192 | 66 | 20 |
| 1 | AMS0229 | 22 | 21 | 41 | 20 | 20 | 20 | 20 | 34 | 20 | 59 | 20 | 61 | 20 | 20 | 20 | 20 | 163 | 67 |
| 2 | UKNP2.2.1 | 20 | 20 | 20 | 20 | 20 | 20 | 20 | 20 | 20 | 20 | 20 | 20 | 33 | 25 | 20 | 20 | 151 | 26 |
| 4 | AMS4d.k9 | 20 | 20 | 20 | 20 | 20 | 20 | 20 | 38 | 20 | 41 | 20 | 20 | 20 | 20 | 20 | 20 | 52 | 74 |
| 3 | UKNP3.2.2 | 20 | 20 | 20 | 20 | 20 | 20 | 20 | 20 | 20 | 20 | 20 | 20 | 20 | 20 | 20 | 20 | 51 | 20 |

Dilution range: 20-4860
Autologous to cocktail

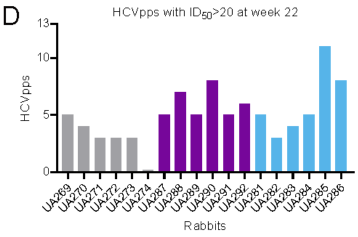


***Supplementary Figure 8: Rabbit AMS0232 full-length E1E2 autologous longitudinal ELISA binding and heterologous neutralization ID50s. A*** *Rabbit serum ELISA endpoint binding titers to membrane-bound AMS0232 E1E2 for all six timepoints. Data has partially been reported previously.[50] Data represented as median with interquartile range.* ***B*** *Heterologous neutralization $ID_{50}$-values plotted for each group against all tested HCVpps. The same data is presented in* ***C****, with the HCVpps of strains included in the cocktail group underlined.* ***D*** *Number of neutralized ($ID_{50}$>20) HCVpps per rabbit. Groups are indicated by color. Significant differences between groups in* ***B*** *were determined using a Kruskal-Wallis test, followed by a Dunn's post-test. Significant differences are indicated on top of the graphs (* <0.05; ** < 0.01).*

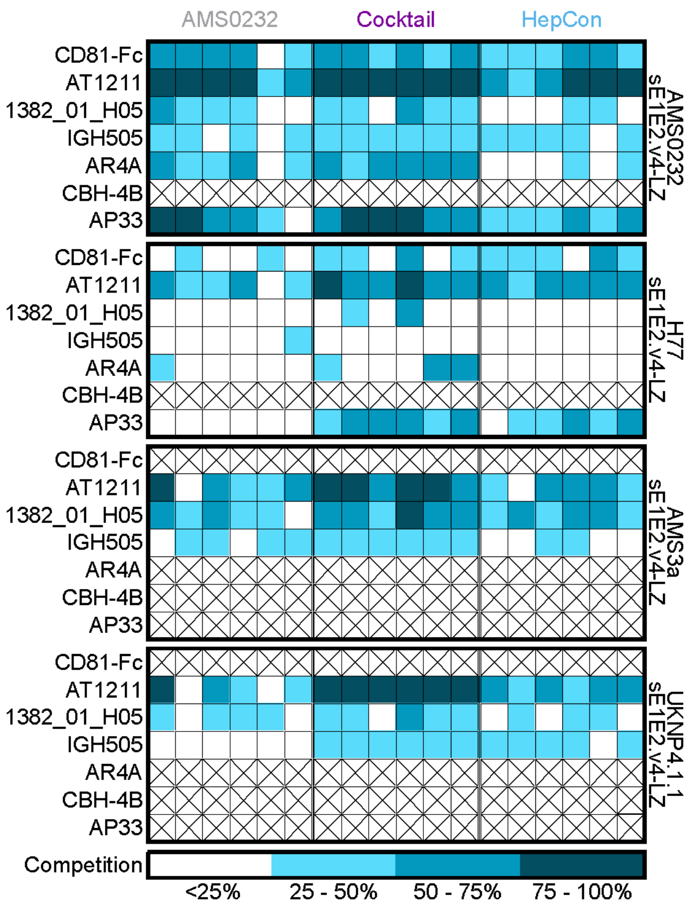


***Supplementary Figure 9: Luminex competition assay.*** *The binding of varying concentrations of HCV bNAbs to sE1E2.v4-LZ coupled Magplex beads was determined after pre-incubation of the beads with rabbit sera, yielding a final percentage of competition with the respective Abs for their known epitopes. Each column represents one rabbit. Stratification of competition is indicated.*

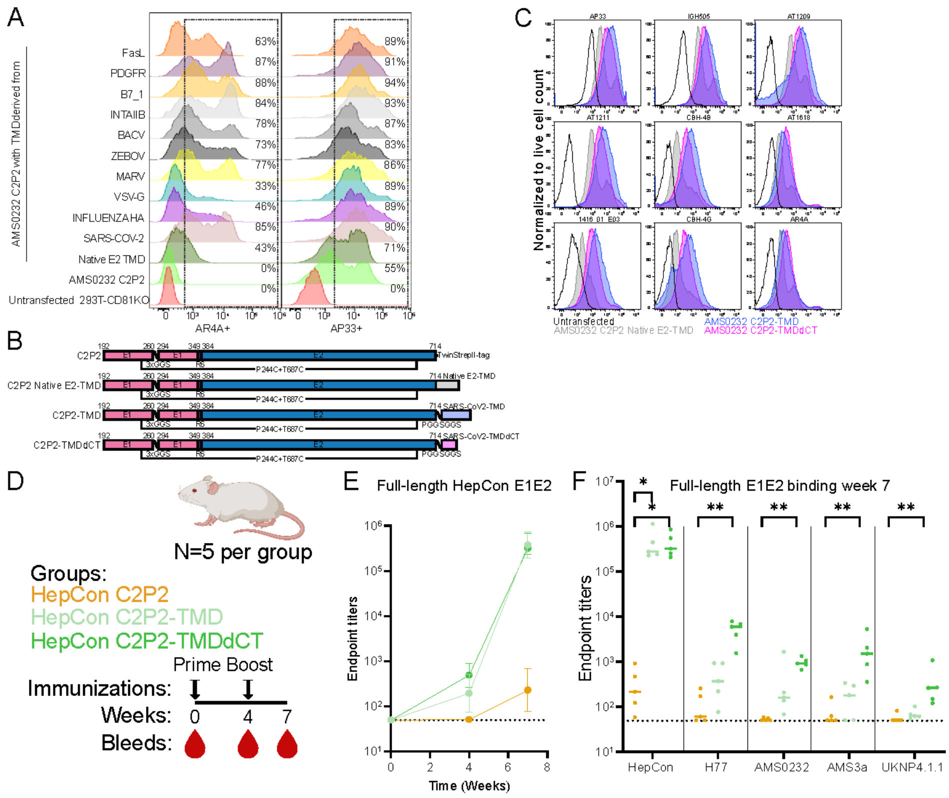


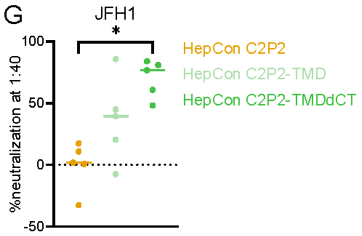


H

| Group | % neutralization JFH1 at 1:40 | | | | |
|---|---|---|---|---|---|
| HepCon C2P2 | 0.7 | 10.8 | -32.6 | 17.3 | 1.8 |
| HepCon C2P2-TMD | 39.4 | 20.5 | 85.7 | 44.8 | -7.5 |
| HepCon C2P2-TMDdCT | 76.7 | 48.0 | 60.7 | 81.0 | 83.9 |

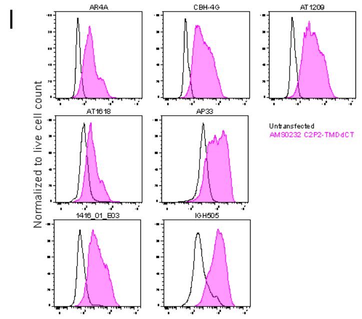


J

| Group | JFH1 $ID_{50}$ | | | | |
|---|---|---|---|---|---|
| AMS0232 C2P2-TMDdCT | 40 | 40 | 40 | 40 | 40 |
| HepCon C2P2-TMDdCT | 213.1 | 40 | 1124 | 40 | 153.9 |

Starting dilution: 40

***Supplementary Figure 10: mRNA design and validation and pilot mice mRNA study. A** Screening of cell surface expression of AR4A and AP33 binding AMS0232 C2P2 fused to heterologous TMDs from human- and virus-derived proteins.[71] The constructs were transfected as DNA in HEK293T cells and screened using flow cytometry. **B** Linear representation of C2P2, C2P2 Native E2-TMD, C2P2-TMD and C2P2-TMDdCT designs with the TMD, Twin-StrepII-tag, linkers and mutations indicated. **C** Flow cytometry analysis of HEK293T cells transfected with DNA encoding optimized membrane-bound E1E2 designs based on AMS0232 C2P2. Engagement of several HCV mAbs was determined for and compared between untransfected HEK293T cells (black line), and HEK293T cells transfected with AMS0232 C2P2 with a Native E2-transmembrane domain (TMD) (grey), a SARS-CoV-2 TMD (blue) and a SARS-CoV-2 TMD without the c-terminal cytoplasmic tail (TMDdCT) (pink). All signals have been normalized to live cell counts. **D** Groups of five female BALB/c mice were immunized at weeks 0 and 4 with 5μg LNP* *encapsulated mRNA encoding HepCon C2P2, HepCon C2P2-TMD or HepCon C2P2-TMDdCT. Bleeds were taken at weeks 0, 4 and 7.* ***E,F*** *Mice serum ELISA endpoint binding titers to full-length HepCon E1E2 for all three timepoints (****E****) and to full-length HepCon, H77, AMS0232, AMS3a and UKNP4.1.1 E1E2 for week 8 (****F****). Data in* ***E*** *is represented as median with interquartile range and in* ***F*** *as median.* ***G*** *Percentage neutralization of several HCVpps measured at 1:40 serum dilution at week 8. Medians are indicated.* ***H*** *Percentage neutralization values used for* ***G****.* ***I*** *Flow cytometry analysis of HEK293T cells transfected with mRNA encoding optimized membrane-bound E1E2 designs based on AMS0232. Engagement of several HCV mAbs was determined for, and compared between, untransfected HEK293T cells (black line), and HEK293T cells transfected with AMS0232 C2P2-TMDdCT (pink). All signals have been normalized to live cell counts.* ***J*** *Neutralization $ID_{50}$-values used for Fig. 2K. Significant differences between groups in (****F,G****) were determined using a Kruskal-Wallis test, followed by a Dunn's post-test. Significant differences are indicated on top of the graphs (* <0.05; ** < 0.01) Figure* ***F*** *was created with Biorender.com.*

A

```
                                                      CDRH1                           CDRH2                                       CDRH3
                                   10        20          31  36 40        50               54          66  70     80         90    95                  103      110
AR3A_HC (Class 3 FRLY)          QVQLLEQSGAEVKTPGSSVRVSCRPPGGNFNSYSINWVRQAPGHGLEWVGTFI---------------PMFGTSKYAQKFQGRVTITADGSSGTAYMDLNSLRSDDTAFYYCVRPET---PRYCSGGFCYG------EFDNW---GQGTLVTVSS
igl-AR3A_HC                     .....V.......K.....K...KAS..T.S..A.S.......Q....M.GI.---------------.I...AN..............E.TS....E.S....E...V.........---.......T...------...Y.---..........

AR3B_HC (Class 3 FRLY)          QVQLLEQSGPEVKKPGSSVKVSCKDSGDTFN-EPVTWVRQAPGQGLEWIGGII---------------PAFGVTKYAQKFQGRVIISADASTATAYLELSSLRSEDTAVYYCAKVG---LRGIVMVGGLAMN-----WLDPW---GQGTQVTVSS
igl-AR3B_HC                     .....V...A..............A..G..SSYAIS...........M....---------------.I..TAN..........T.T..E..S...M...............R..---....AVA.T....-----.F...---..........

AR3C_HC (Class 3 FRLY)          QVQLLEQSGAEVKKPGSSVKVSCETSGGTFDNYALNWVRQAPGQGLEWIGGVV---------------PLFGTTRNAQKFQGRVTISDDKSTGTGHMELRSLRSEDTAVYYCVRSVT---PRYCGGGFCYG------EFDYW---GQGTLVTVSS
igl-AR3C_HC                     ....V.................KA.....SS..IS...........M..II---------------.I...ANY...........TA....S.AY.................A...---....S......------....---..........

AR3D_HC (Class 3 FRLY)          QVQLLEQSGAEVKKPGSSVKVSCKASGDTFRSYVITWARQAPGQGLEWMGAII---------------PFFGTTNLAQKFQGRVTITADESTKTVYMDLSSLRSDDTAVYYCAKAGDLSVGGVLAGGVPHLR-----HFDPW---GQGTLVTVSS
igl-AR3D_HC                     .....V....................G..S..A.S.V...........G..---------------.I...A.Y...................S.A..E.....E........R....TF...I.........-----.....---..........

AT1209_HC (Class 3 FRLY)        --QVLVQSGAEVKKPGSSVRVSCKASGDTFKTHAISWVRQAPGQGLEWVGGTVRQAPGDGLELLGGFVPILAPANDAQKFQGRVTITADGSTGPVYMDLSTLTSEDTAMYYCVTSLSEPIPRSCRGGRCYSGP--FDAFGVW---GQGTMVTVSS
igl-AT1209_HC                   --...............K.......G..SSY...............M..II---------------..FGT..Y.............E..STA..E..S.R.....V...A...........S..S.....--....D..---..........

HEPC3_HC (Class 1 FRLY)         QVQLV-QSGAEVKKPGSSVKVSCKASGGTLNSYEITWVRQAPGQGLEWMGGIT---------------PIFET-TYAQKFQGRVTITADESTSTTYMELSSLRPEDTAVYYCARDGV----RYCGGGRCYN------WFDPW---GQGTLVTVSS
igl-HEPC3_HC                    .....-......................FS..A.S...............I---------------...G.AN.................A........S..............----...S..S...------.....---..........

HEPC74_HC (Class 1 FRLY)        QVQLV-QSGAEVKKPGSSVKVSCTTSGGTYINYAISWVRQAPGQGLEWVGGMS---------------PISNTPKYAQKFQGRVTITADESTSTTYMELSSLRPEDTAVYYCARDLL----KYCGGGNCHSL-----LVDPW---GQGTLVTVSS
igl-HEPC74_HC                   .....-.................KA....FSS...............M..II---------------..FG.AN.................A........S..............----...S..S.Y..-----.....---..........

HC84.26_HC (Class 2 FRLY)       QVQLV-QSGAEVKKPGSSVKVSCEASGGTLSNYVITWVRQAPGQGLEWMGGFI---------------PTFRTAMYAQGFQGRVTITADESTSIAYMELTNLRSEDTAVYYCARGP-------LSRG----------YYDYW---GQGTLVTVSS
igl-HC84.26_HC                  .....-.................K.....F.S.A.S...............I.---------------.I.G..N...K...............T.....SS...................-------....----------.....---..........

1416_01_E03_HC (Class 2 FRLY)   QVQLV-QSGAEVKKPGSSVKVSCKASGGTFSRFGISWVRQAPGQGLEWMGGII---------------PKFGTTNYAQKFQGRVTITADKSTNIAYLEVSSLTSEDTAVYYCAREVAP------SNSFFLS------WFDPW---GQGTLVTVSS
igl-1416_01_E03_HC              .....-......................SYA....................---------------.T...A....................ST..M.L...R..................------..WL...------.....---..........

1198_05_G10_HC (Class 2 FRLY)   QVQLV-QSGAEVKKPGSSVKVSCRTSGGTVN--TLHWVRQAPGQGLEWMGSIF---------------PLLGVPTYAQKFQGRVTVTADRSTSTAYMELRSLRADDTAVYYCAKDGVG----WSGHGPSQWS-----GVDVW---GPGTTVTVSS
igl-1198_05_G10_HC              .....-.................KA....FSSYAIS.............R.I---------------.I..IAN..........I...K.........S...SE.........R...----...Y......-----.....---.Q........

1382_01_H05_HC (Class 2 FRLY)   QVQLV-QSGAEVKKPGSSVKVSCTTSGGSFGRDAVSWVRQAPGQGLEPMGGII---------------PIFGTSSYAQTFQGRVTFTAGKSTSTAYMELSSLKSEDTAVYYCVRWSFG------DYGLFLT------EGDYW---GQGTLVIVSS
igl-1382_01_H05_HC              .....-.................KA...T.SSY.I...........W.....---------------.....AN...K.....I..D.............R.........A....------.......------.....---......T...

1334_03_A04_HC                  QVQLV-QSGAEVKKPGSSVKVSCKASGGTFRSYTFSWVRQAPGHGLEWMGGII---------------SVGNSATYAQKFQGRVTITADKSTSTVYMELSGLTSEDTAIYFCARDGNFD---WRTARNEQN------WFDPW---GQGTLVTVSS
igl-1334_03_A04_HC              .....-.......................S..AI.......Q.........---------------PIFGT.N.................A.....S.R.....V.Y.........---.........------.....---..........

AR4A_HC                         EVQLLEQSGPEVKKPGDSLRISCKMSGDSLVTTWIGWVRQKPGQGLEWMG------IINPGDSSTNIYPGDSATRYGPSFQGQVTISIDKSTSTAYLQWNNVKASDTGIYYCARHVPVPISGTFLWREREMHDFG--YFDDW---GQGTLVIVSS
igl-AR4A-1_HC im3               ....V-...A......E..K....G..Y.FTSY.......M..K......------...............D...S...........A...I.......SSL.....AM...........S..............--.....---..........
igl-AR4A-2_HC g3                ....V-...A......E..K....G..Y.FTSY.......M..K......------..Y...---------.D...S...........A...I.......SSL.....AM.............SYR..........--...Y.---......T...
igl-AR4A-3_HC ig3               ....V-...A......E..K....G..Y.FTSY.......M..K......------..Y....D............S...........A...I.......SSL.....AM.............SYR..........--...Y.---......T...

AT1618_HC                       EVNLL-ESGGGLVQPGGSLRLSCVASGGTVASDAMSWVRQAPGKGLTWVS---------------AISGIGVYTYYADSVKGRFTISRDKSKNTLYLQMNSLRVEDTAVYYCAKGDFG-------PGSLHNDP---FYLKSW---GQGTLVAVSS
igl-AT1618_HC                   ..Q..-.................A...F.FS.Y............E...---------------....S.GS..................N.............A...................-------........---.....---..........

AT1211_HC                       QVQLV-QSGAEVKKPGSSVKVSCKASGGTFSGYGITWVRQAPGQGLEWMGTII---------------PVSATETYAQRLQGRVTISADEHSTTSYMEVSSLKSEDTALYYCARHDY-------FWGT---------PLDIW---GQGTMVIVSS
igl-AT1211_HC                   .....-......................S.A.S...............R..---------------.IFG.AN...KF......T...STS.A...L...R.....V.........-------V...---------.....---......T...
```

B

```
                                         CDHL1                          CDRL2                          CDRL3
                            10      20  24  30      35   40       50   57 60     70       80      89               98
AR3A_LC              ELTLTQSPGTLSLSPGKRATLSCRASQSVSGS----YLAWYQQKPGQAPRLLIYGASNRATGIPHRFSGSGSGTDFTLTISRLEPEDFAVYYCQQYG----------SSP-TFGQGTRVDIK
igl-AR3A_LC          .IV.............E.............S.----..................S......D............................ ....----------...-........E..
```

```
AR3A_LC              ELTLTQSPGTLSLSPGKRATLSCRASQSVSGS----YLAWYQQKPGQAPRLLIYGASNRATGIPHRFSGSGSGTDFTLTISRLEPEDFAVYYCQQYG----------SSP-TFGQGTRVDIK
igl-AR3A_LC          .IV.............E.............S.----..................S......D................................----------...-........E..

AR3B_LC              AAELTQSPGTLSLSPGERATLSCRASQSVSSS----YLAWYQQKPGQAPRLLIYGASSRATGIPDRFSGSGSGTDFTLTISRLEPEDFAVYYCQQYG----------SSPQTFGQGTKVEIK
igl-AR3B_LC          EIV.............................----........................................................----------...............

AR3C_LC              EIELTQSPATLSVSPGERATLSCRASQSVS-S----NLAWYQQKPGQAPRLLIYGASTRATGIPARFSGSGSGTEFTLTVSRLEPEDSAVYFCQQYY----------RSPLTFGGGTKVEIK
igl-AR3C_LC          ..VM..........................-.----.......................................I.S.QS..F...Y.....N----------...............

AR3D_LC              EIELTQSPGTLSLSPGERATLSCRAGQTVASN----SLAWYQHKPGQAPRLLIYGASIRASGIPDRFSGSGSGTDFTLTISRLEPEDFAVYYCQQYG----------LSS-TFGQGTRLEIK
igl-AR3D_LC          ..V......................S.S.S.S----Y.....Q..............S..T....................................----------...-...........

AT1209_LC            EIMLTQSPVTLSVSPGERATLSCRASQSIG-T----NLAWYQQKPGQAPRLLIHGASTRATGVPVSFSGSGSGTEFTLTISSLQSEDFAVYYCQQYN----------NWPLTFGGGTKVDFK
igl-AT1209_LC        ..VM....A...................VS-S----..............Y........I.AR...............................----------............EI.

HEPC3_LC             DIQMTQSPSSLSASVGDRVTITCRAGQNIN-N----YLNWYQQKPGKAPKVLIYAASNLQSGVPSRFSGSGSGTDFTLTISSLQPEDFATYYCQQSHS---------TVR-TFGQGTKVEIK
igl-HEPC3_LC         .........................S.S.S-S----..............L......S.......................................Y.---------...-...........

HEPC74_LC            DIVMTQSPSTLSASVGDRVTISCRASQSIS-S----WLAWYQQKPGRAPKLLIYKASSLETGVPSRFSGSGSGTEFTLTISSLQPDDFATYYCQHYNT---------Y-LFTFGPGTKVDLK
igl-HEPC74_LC        ..Q..................T........-.----..........K..............S................................Q..S---------.-...........I.

HC84.26_LC           SYVLTQ-PPSVSVAPGKTARITCGGN--NIGSK---SVHWYQQKPGQAPVLVVYDDSDRPSGIPERFSGSNSGNTATLTISRVEAGDEADYYCQVWDS----------SSVVFGGGTKLTVL

1416_01_E03_LC       QSALTQ-PASVSGSPGQSITISCTGTSSDIGSYN--LVSWYQQHPGKAPKLMISEVTERPSGVSARFSGSKSGNTASLTISGLQAEDEADYFCCSYARG--------STYWIFGGGTKLTVL
igl-1416_01_E03_LC   ......-.......................V....--.................Y..SK......N..........................Y.....G.--------....V..........

1198_05_G10_LC       DIVMTQSPLSLPVTPGEPASISCTSSQSLLHSTGYNYLDWYVQKPGQSPQLLIYLGSIRASGVPDRFSGSGSGTDFTLRISRVEAGDVGIYYCMQALE---------IPRLTFGGGTKLEIK
igl-1198_05_G10_LC   .......................R........N........L...................N..................K......E...V........---------T..........V...

1382_01_H05_LC       EIVLTQSPASLSLSPGERATLSCRASQSVD-K----YFAWYQQKPGQAPRLLIYETSKRATGIPARFSGSGSGTDFTLTISNLEPDDFAIYYCHHRGNW--------PPSFTFGQGTRLEIK
igl-1382_01_H05_LC   .........T.........................S-S----.L.................DA.N.......................S...E...V...QQ....--------...............

1334_03_A04_LC       DIQMTQSPSSLSASVGDRVTLTCRASQTIG-N----FLNWYQQKPGRAPKLLVYGASNLQSGVPSRFSGSGSGTDFTLTISSLEPEDFATYYCQQTYN---------SPRVTFGQGTRLDIK
igl-1334_03_A04_LC   ....................I......S.S-S----Y.........K.....I.A..S.........................Q............S.S---------............E..

AR4A_LC              ELTLTQSPGTLSLSPGERATLSCRASQSVSNN----YLAWYQQKPGQAPRLLIYGASSRATGIPDRFSGSGSGTGFTLIISRLEPEDFAVYYCQQYG----------SSSITFGQGTRLEIK
igl-AR4A_LC          .IV...........................SS----..............................D...T..................----------...............

AT1618_LC            DIQMTQSPSTLSASVGDRVTITCRASQSIS-R----WLAWYQQKPGKAPKLLIYDASSLESGVPSRFSGSGSGTEFTLTISSLQPDDFATYYCQQYHN---------Y-EWTFGHGTKVDFK
igl-AT1618_LC        ..............................-S----......................................................NS---------.-.............

AT1211_LC            EIVLTQSPDFQSVTPKEKVTITCRASQSIGSN-----LHWYQQKPGQSPKLLIKYASQSFSGVPSRFSGSGSGTDFTLTINSLEAEDAATYFCHQSYN---------LPR-TFGGGTKVEIK
igl-AT1211_LC        ...............................S-----.........D.....................................Y....SS---------...-...........
```

C

| HCV bNAb | IGHV | IGHD | IGHJ | IGHV length amino acids | IGHV amino acid mutations | IGHV amino acid affinity maturation (%) | Number of improbable amino acid mutations | insertion or deletion | IGKV/IGLV | IGKJ/IGLJ | IGKV/IGLV length amino acids | IGKV/IGLV amino acid mutations | IGKV/IGLV amino acid affinity maturation (%) | Number of improbable amino acid mutations |
|---|---|---|---|---|---|---|---|---|---|---|---|---|---|---|
| AR3A | IGHV1-69*01 | IGHD3-10*01 | IGHJ4*02 | 99 | 24 | 24 | 2 | No | IGKV3-20*01 | IGKJ2*01 | 95 | 6 | 6 | 1 |
| AR3B | IGHV1-69*01 | IGHD6-19*01 | IGHJ5*02 | 98 | 21 | 21 | 2 | No | IGKV3-20*01 | IGKJ2*01 | 95 | 3 | 3 | 0 |
| AR3C | IGHV1-69*06 | IGHD6-13*01 | IGHJ4*02 | 99 | 20 | 20 | 2 | No | IGKV3D-15*01 | IGKJ4*01 | 94 | 9 | 10 | 0 |
| AR3D | IGHV1-69*01 | IGHD3-16*03 | IGHJ2*01 | 99 | 15 | 15 | 1 | No | IGKV3-20*01 | IGKJ2*02 | 95 | 9 | 9 | 0 |
| AT1209 | IGHV1-69*01 | IGHD2-15*01 | IGHJ4*02 | 97 | 21 | 22 | 2 | Yes | IGKV3D-15*01 | IGKJ4*01 | 94 | 10 | 11 | 0 |
| HEPC3 | IGHV1-69*01 | IGHD1-26*01 | IGHJ5*02 | 97 | 10 | 10 | 1 | No | IGKV1-39*01 | IGKJ2*02 | 95 | 7 | 7 | 0 |
| HEPC74 | IGHV1-69*01 | IGHD2-2*01 | IGHJ5*02 | 98 | 14 | 14 | 1 | No | IGKV1-5*05 | IGKJ2*01 | 94 | 6 | 6 | 1 |
| HC84.26 | IGHV1-69*01 | IGHD4-11*01 | IGHJ4*02 | 98 | 13 | 13 | 2 | No | IGLV3-21*03 | IGLJ2*01 | 93 | N/A | N/A | 0 |
| 1416_01_E03 | IGHV1-69*14 | IGHD3-9*01 | IGHJ5*02 | 98 | 10 | 10 | 1 | No | IGLV2-23*02 | IGLJ3*02 | 98 | 7 | 7 | 2 |
| 1198_05_G10 | IGHV1-69*09 | IGHD3-3*01 | IGHJ6*02 | 96 | 19 | 20 | 3 | Yes | IGKV2-28*01 | IGKJ4*01 | 100 | 8 | 8 | 1 |
| 1382_01_H05 | IGHV1-69*06 | IGHD4-17*01 | IGHJ4*02 | 98 | 15 | 15 | 1 | No | IGKV3-11*01 | IGKJ5*01 | 96 | 12 | 13 | 3 |
| 1334_03_A04 | IGHV1-69*06 | IGHD3-9*01 | IGHJ5*02 | 98 | 15 | 15 | 2 | No | IGKV1-39*01 | IGKJ5*01 | 95 | 12 | 13 | 0 |
| AR4A | IGHV5-51*01 | IGHD2-21*01** | IGHJ4*02 | 99 | 21 | 21 | 3 | Yes | IGKV3-20*01 | IGKJ5*01 | 95 | 6 | 6 | 1 |
| AT1618 | IGHV3-23*05 | IGHD3-22*01 | IGHJ1*01 | 98 | 12 | 12 | 3 | No | IGKV1-5*01 | IGKJ1*01 | 94 | 3 | 3 | 0 |
| AT1211 | IGHV1-69*18 | IGHD3-16*01 | IGHJ3*02 | 98 | 19 | 19 | 3 | No | IGKV6-21*01 | IGKJ4*01 | 95 | 5 | 5 | 2 |

**D gene used for igl-AR4A-ig3/g3: IGHD3-16*02

D

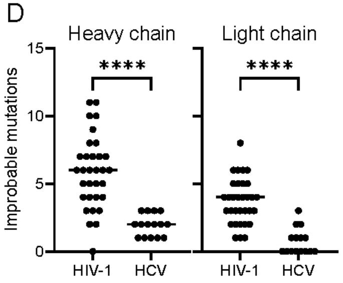


***Supplementary Figure 11: Inferred igl-bNAb sequences, gene usage and mutation probability scoring. A*** *Aligned sequences of the HC of the different igl-bNAbs aligned to their mature counterpart. The CDRH1, CDRH2 and CDRH3 are indicated in blue, orange and red, respectively. Where applicable, classes of*

*FRLY Abs are specified.[47] **B** Aligned sequences of the LC of the different igl-bNAbs aligned to their mature counterpart. The CDRL1, CDRL2 and CDRL3 are indicated in blue, orange and red, respectively. The LC of HEPC3 and HEPC74, and the HC and LC of AT1209, HC84.26, 1416_01_E03, 1198_05_G10, 1382_01_H05, 1334_03_A04, AR4A, AT1618 and AT1211 in **A** and **B** were inferred using IMGT/V-QUEST software tool.[85,121] **C** HC and LC gene alleles incorporated in the corresponding igl-bNAbs. Indicated are the IGHV and IGKV/IGLV affinity maturation on amino acid level, the numbers of improbable (<1% likely) mutations based on ARMADiLLO[86,87] analysis and whether the IGHV sequence contains an insertion or deletion. Identical scoring was performed for all used HC and LC sequences and <1% likely mutations are highlighted in bold and underscored in **A** and **B**. **D** Comparison of improbable (<1% likely) mutations based ARMADiLLO scoring of HIV-1 (n=32) and HCV (n=15) bNAbs. The selected HIV-1 bNAbs consist of CD4 binding site, V3 glycan, V2 apex, MPER, fusion peptide and silent face targeting bNAbs, obtained from precomputed ARMADiLLO analyses. Significant differences are indicated on top of the graphs (**** < 0.0001)*

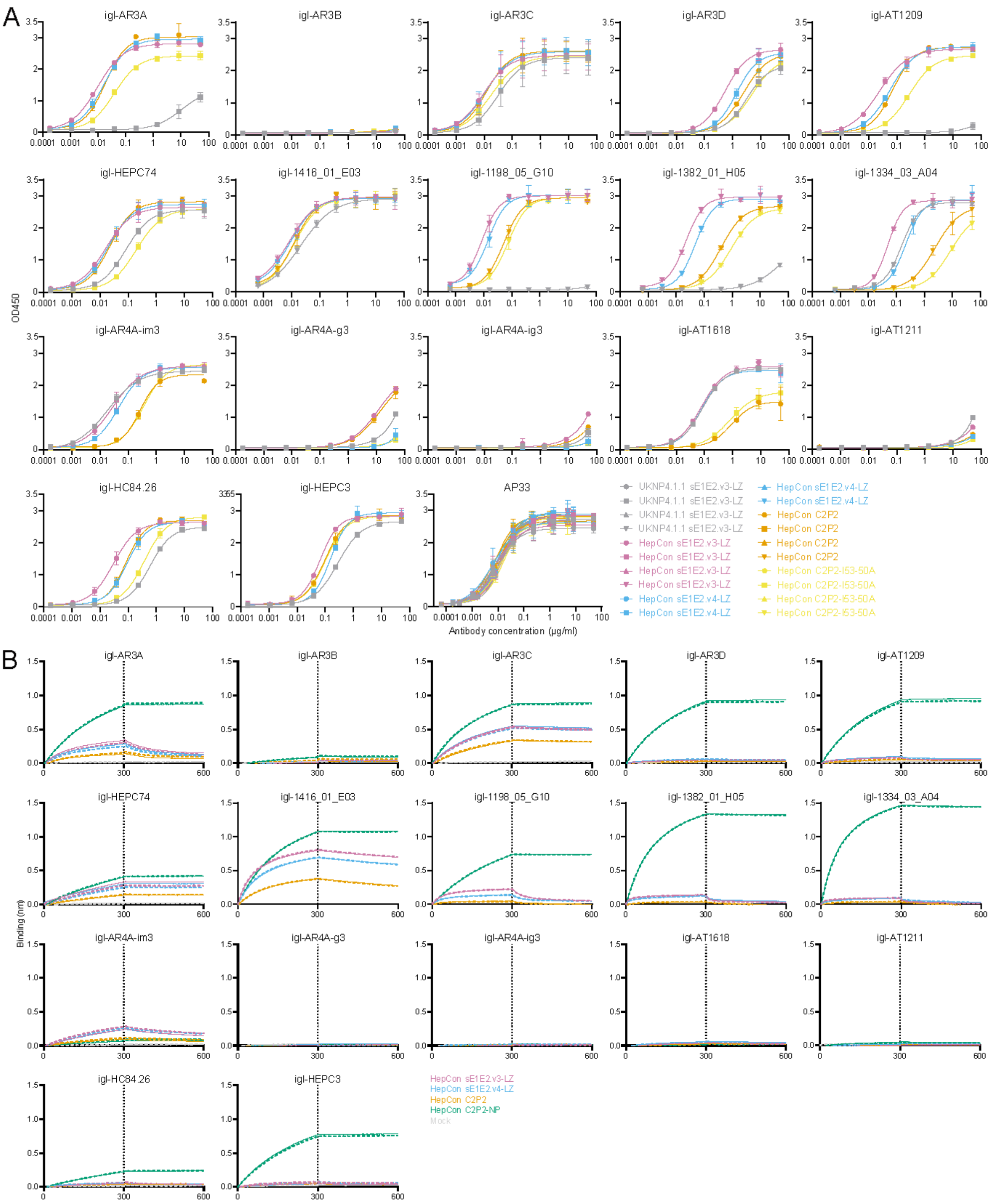
A
igl-AR3A
igl-AR3B
igl-AR3C
igl-AR3D
igl-AT1209
igl-HEPC74
igl-1416_01_E03
igl-1198_05_G10
igl-1382_01_H05
igl-1334_03_A04
igl-AR4A-im3
igl-AR4A-g3
igl-AR4A-ig3
igl-AT1618
igl-AT1211
igl-HC84.26
igl-HEPC3
AP33
OD450
Antibody concentration (µg/ml)
UKNP4.1.1 sE1E2.v3-LZ
HepCon sE1E2.v3-LZ
HepCon sE1E2.v4-LZ
HepCon C2P2
HepCon C2P2-I53-50A
B
igl-AR3A
igl-AR3B
igl-AR3C
igl-AR3D
igl-AT1209
igl-HEPC74
igl-1416_01_E03
igl-1198_05_G10
igl-1382_01_H05
igl-1334_03_A04
igl-AR4A-im3
igl-AR4A-g3
igl-AR4A-ig3
igl-AT1618
igl-AT1211
igl-HC84.26
igl-HEPC3
Binding (nm)
HepCon sE1E2.v3-LZ
HepCon sE1E2.v4-LZ
HepCon C2P2
HepCon C2P2-NP
Mock

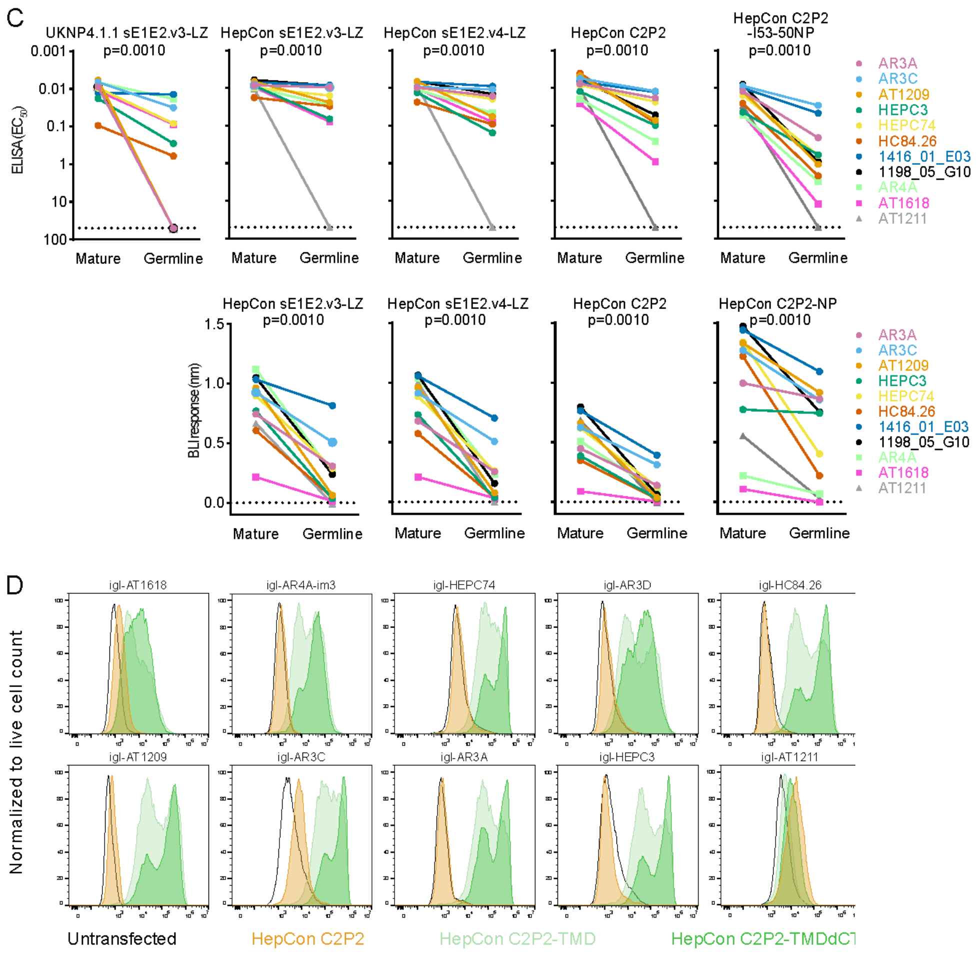


***Supplementary Figure 12: ELISA, BLI and flow cytometry analyses of binding of igl-bNAbs targeting different epitopes on E1 and E2. A** ELISA binding curves for HepCon sE1E2.v3-LZ, sE1E2.v4-LZ, C2P2 and C2P2-I53-50A were obtained using Strep-TactinXT ELISA with UKNP4.1.1 sE1E2.v3-LZ tested as comparator.[50] AP33 binding was determined as loading control. Results obtained in different experiments are indicated by different symbols. The means and SD are shown. **B** Binding profiles for HepCon sE1E2.v3-LZ, sE1E2.v4-LZ, C2P2 and C2P2-NP with the igl-bNAbs were obtained using BLI. The indicated igl-bNAbs were immobilized onto protein A biosensors and incubated for 300 seconds with 250 nM sE1E2.v3-LZ, sE1E2.v4-LZ, C2P2 or C2P2-NP and subsequently incubated for 300 seconds with running buffer. Replicates are shown in corresponding colors, as full and dotted lines. **C** (top) Comparison of $EC_{50}$-values from Supplementary Figures 5A, 12A, 14A,B for binding to bNAbs and their inferred germline counterparts for UKNP4.1.1 sE1E2.v3-LZ, HepCon sE1E2.v3-LZ, sE1E2.v4-LZ, C2P2 and C2P2-NP. (bottom) Comparison of the maximum BLI response values from Supplementary Figures 5B and 12B for binding to bNAbs and their inferred germline counterparts for HepCon sE1E2.v3-LZ, sE1E2.v4-LZ, C2P2 and C2P2-NP. The p-values for Wilcoxon matched-pairs signed-rank tests are shown. **D** Flow cytometry analysis of HEK293T cells transfected with mRNA encoding optimized membrane-bound E1E2 designs based on HepCon. Engagement of several HCV igl-bNAbs was determined for, and compared between, untransfected HEK293T cells (black line) and HEK293T cells transfected with HepCon C2P2 (orange), HepCon C2P2-TMD (light green) and HepCon C2P2-TMDdCT (green). All signals have been normalized to live cell counts. All ELISA and BLI analyses were performed in duplo.*

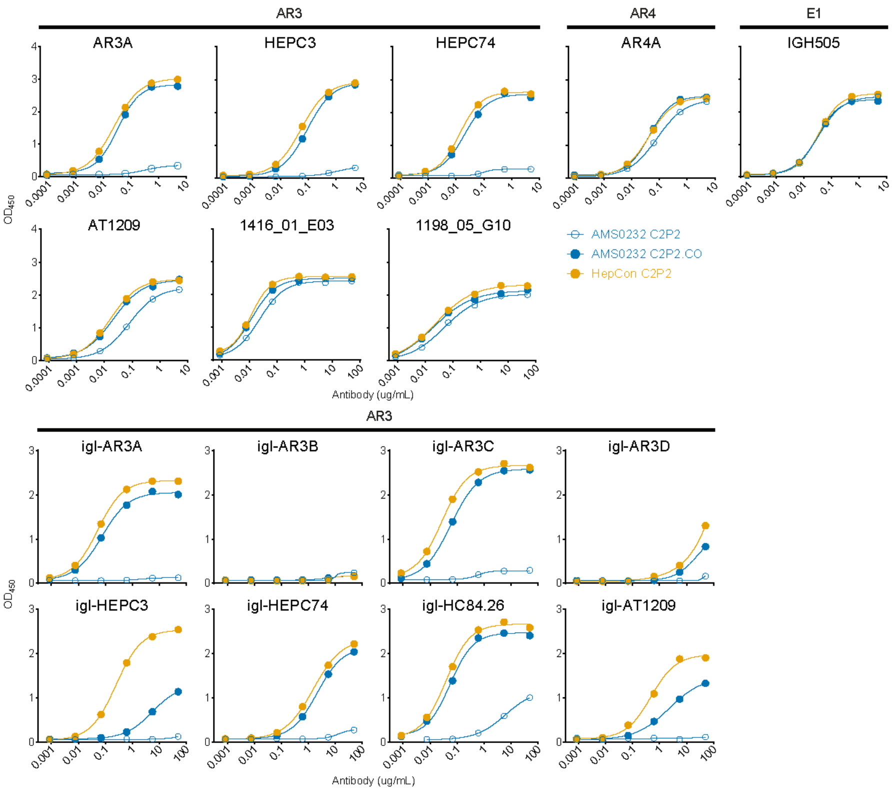


***Supplementary Figure 13: ELISA binding profiles of AMS0232 C2P2.CO. A*** *AMS0232 C2P2.CO binding was determined for a subset of bNAbs (top) and igl-bNAbs (bottom). AMS0232 C2P2 and HepCon C2P2 were used as comparators. Targeted epitopes are indicated.*

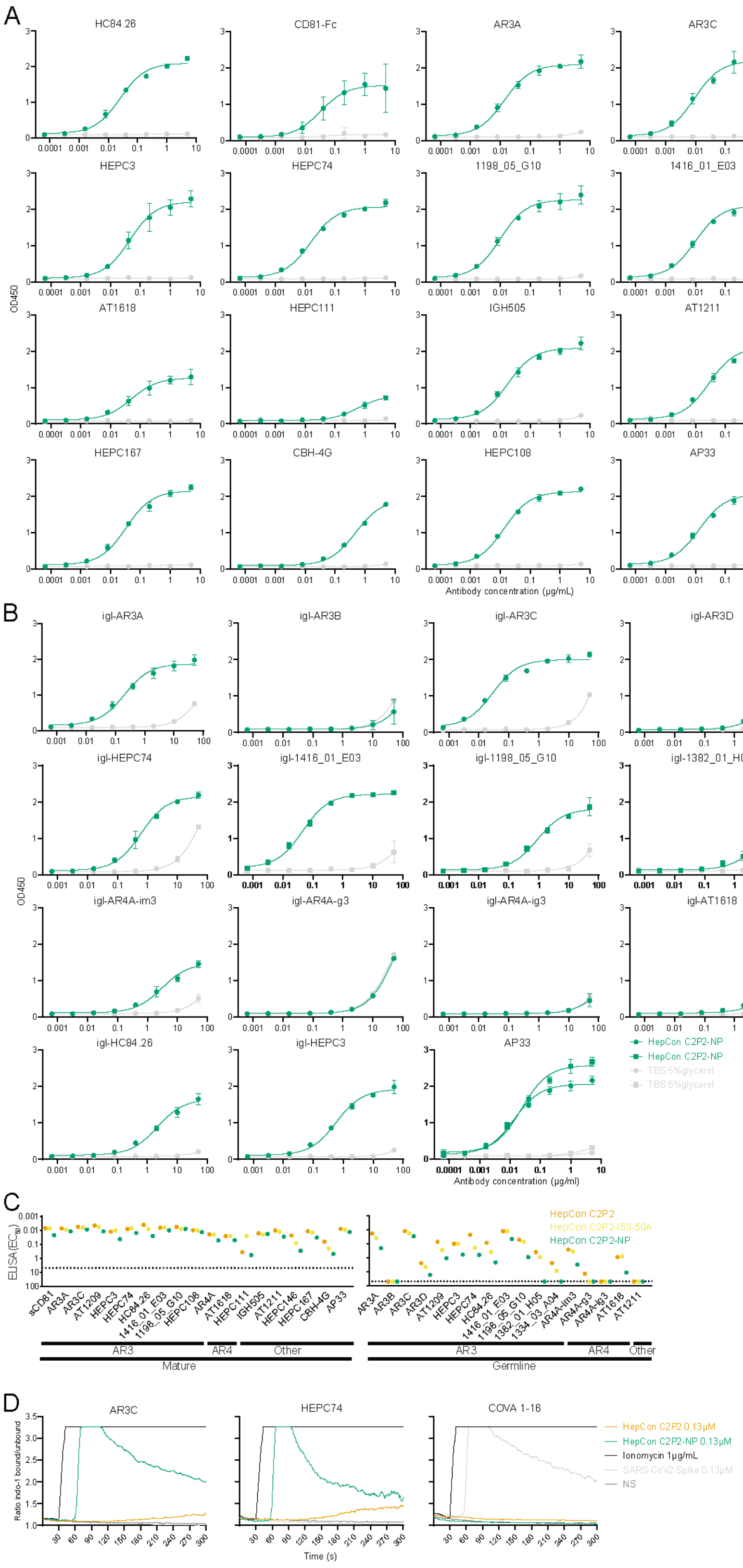


***Supplementary Figure 14: ELISA binding profiles and B cell activation of HepCon C2P2-NP. A,B** ELISA binding curves of bNAbs (**A**) and igl-bNAbs (**B**) for HepCon C2P2-NP were obtained using Galanthus nivalis lectin ELISA with TBS + 5% glycerol as control. AP33 binding was determined as loading control. Results obtained in different experiments are indicated by different symbols. All ELISAs were performed in duplo. **C** Plotted EC$_{50}$-values from bNAb and igl-bNAb binding ELISAs for HepCon C2P2, C2P2-I53-50A*

*and C2P2-I53-50NP, derived from Supplementary Figures 5A, 12A, 14A,B.* ***D*** *Activation was determined by measuring calcium flux in Ramos B cells expressing mature HCV-specific AR3C, HEPC74 or the SARS-CoV-2-specific COVA1-16 as BCR. Ionomycin and absence of stimulation were used as positive and negative controls, respectively. The combination of SARS-CoV2 spike and the COVA1-16 cell line is the positive control for the assay and the SARS-CoV-2 spike antigen served as an additional negative control for the HCV cell lines. Datapoints in* ***A*** *and* ***B*** *are represented as mean with SD and in* ***C*** *the means are indicated.*

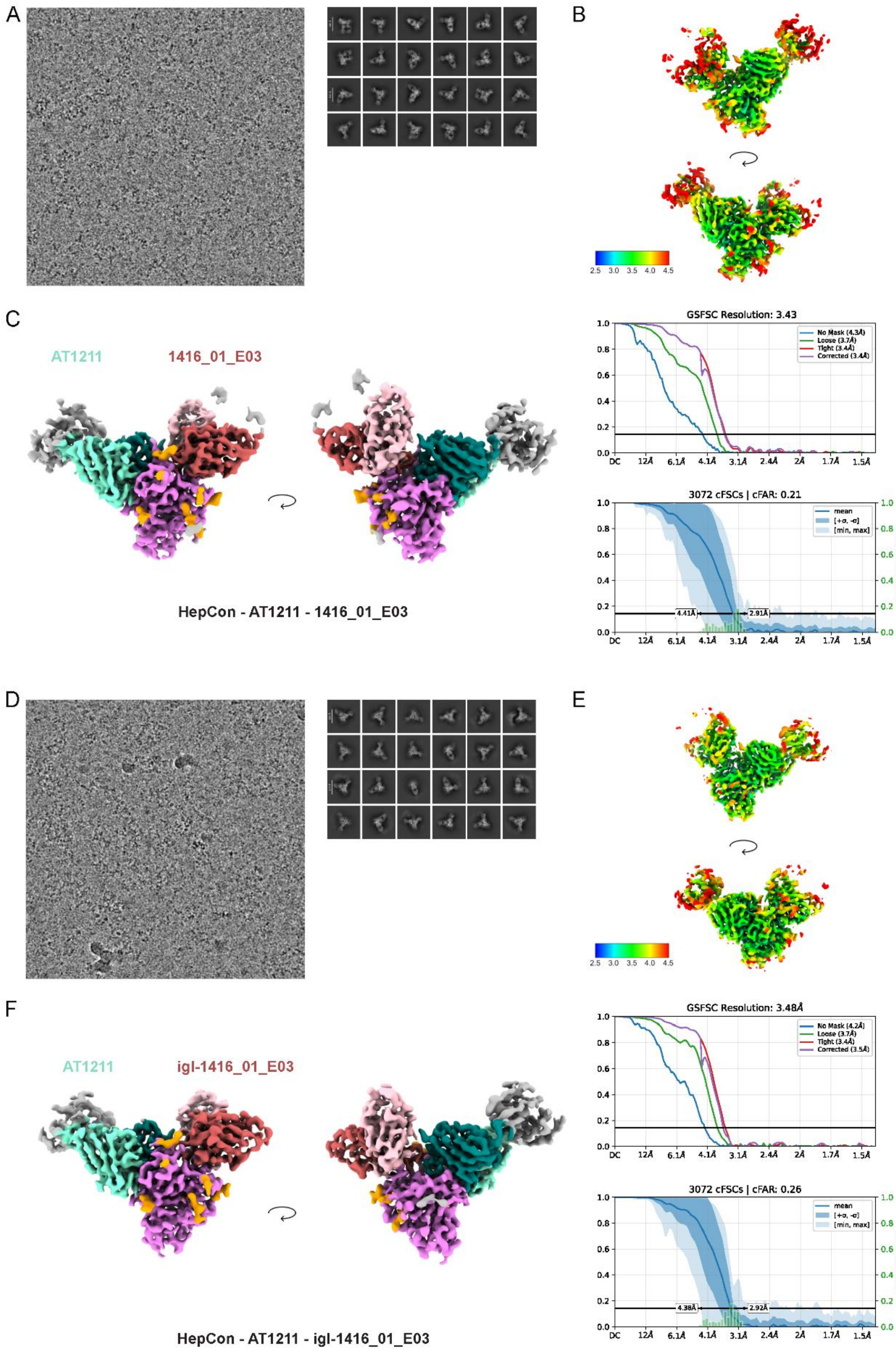
A
B
2.5 3.0 3.5 4.0 4.5
C
AT1211
1416_01_E03
HepCon - AT1211 - 1416_01_E03
GSFSC Resolution: 3.43
No Mask (4.3Å)
Loose (3.7Å)
Tight (3.4Å)
Corrected (3.4Å)
DC 12Å 6.1Å 4.1Å 3.1Å 2.4Å 2Å 1.7Å 1.5Å
3072 cFSCs | cFAR: 0.21
mean
[+σ, -σ]
[min, max]
relative frequency
4.41Å
2.91Å
D
E
2.5 3.0 3.5 4.0 4.5
F
AT1211
igl-1416_01_E03
HepCon - AT1211 - igl-1416_01_E03
GSFSC Resolution: 3.48Å
No Mask (4.2Å)
Loose (3.7Å)
Tight (3.4Å)
Corrected (3.5Å)
DC 12Å 6.1Å 4.1Å 3.1Å 2.4Å 2Å 1.7Å 1.5Å
3072 cFSCs | cFAR: 0.26
mean
[+σ, -σ]
[min, max]
relative frequency
4.38Å
2.92Å

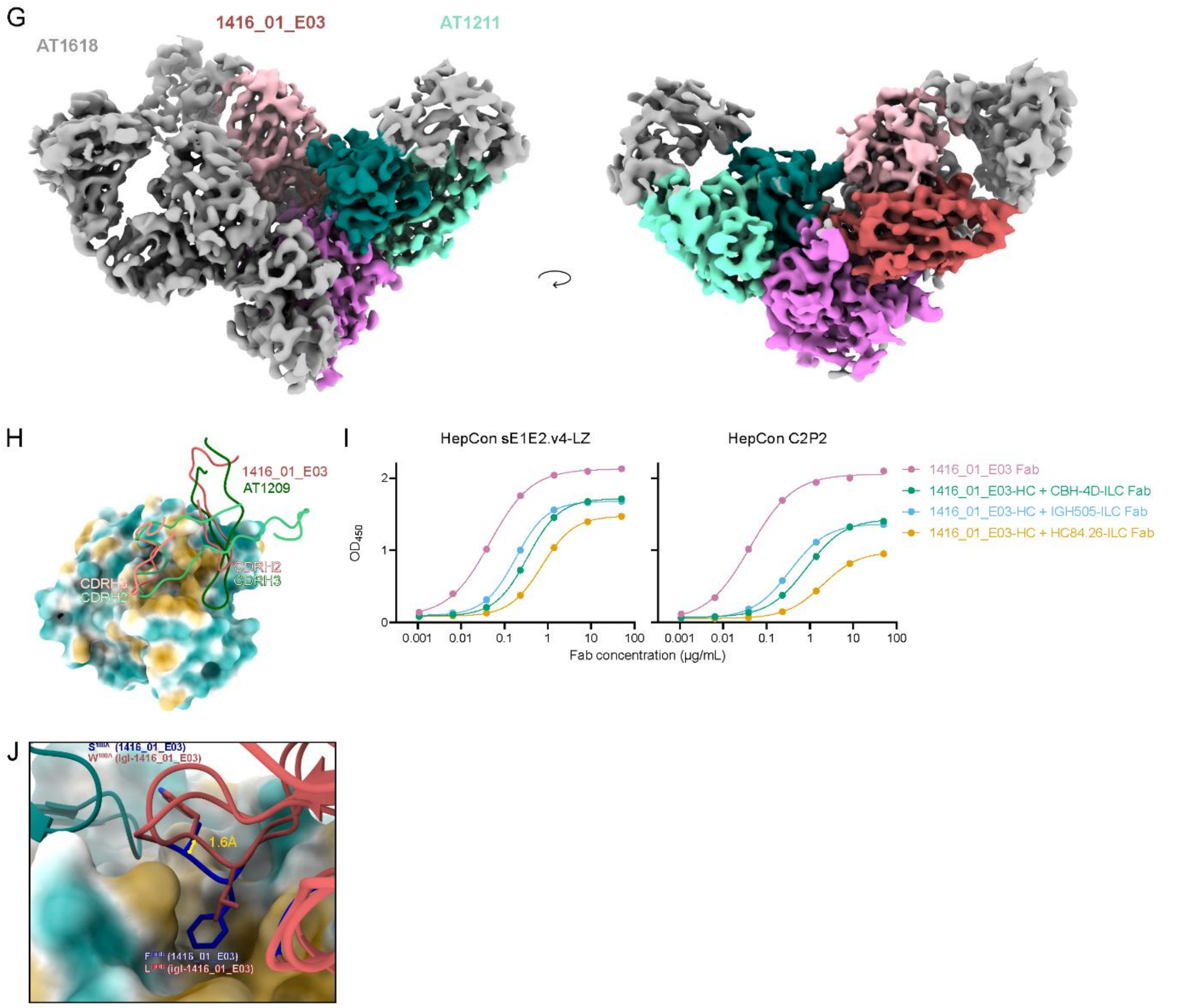


***Supplementary Figure 15: Cryo-EM of HepCon sE1E2.v4-LZ in complex with AT1211 and igl-1416_01_E03 or AT1211, 1416_01_E03 and AT1618. A,D*** *Representative micrographs and 2D classes of AT1211-E1E2-1416_01_E03 (**A**) or igl-1416_01_E03 (**D**). **B,E** Local resolution estimates of AT1211-E1E2-1416_01_E03 (**B**) or igl-1416_01_E03 (**E**). **C,F** Cryo-EM maps of the complexes with individual protein densities colored, with corresponding FSC. AT1211 heavy and light chains are colored teal and light blue and 1416_01_E03 / igl-1416_01_E03 heavy and light chains red and pink. **G** Cryo-EM map of the complex with all three bNAbs, with AT1211 in teal/light blue, 1416_01_E03 in red/pink, and E2 in violet. Another density is clearly visible for AT1618, however severe orientation bias impeded our effort to build a high-resolution model of this complex. **H** CDRH2 and CDRH3 of AT1209 from the AT1211-AT1209-E1E2 cryo-EM map and of 1416_01_E03 with HepCon E2 from the AT1211-E1E2-1416_01_E03 cryo-EM map. E2 is represented as a hydrophobic surface, AT1209 is colored shades of green and 1416_01_E03 shades of red. **I** ELISA binding curves for HepCon sE1E2.v4-LZ and C2P2 against 1416_01_E03 fragment antigen-binding domain (Fab) and Fabs comprised of the 1416_01_E03-Fab HC with varying lambda LCs, using Strep-TactinXT ELISA. **J** CDRH3 – E2 interactions for 1416_01_E03 and igl-1416_01_E03. E2 is represented as a hydrophobic surface, AT1211 is colored teal and 1416_01_E03 / igl-1416_01_E03 red. Somatically mutated residues on 1416_01_E03 are colored in dark blue.*

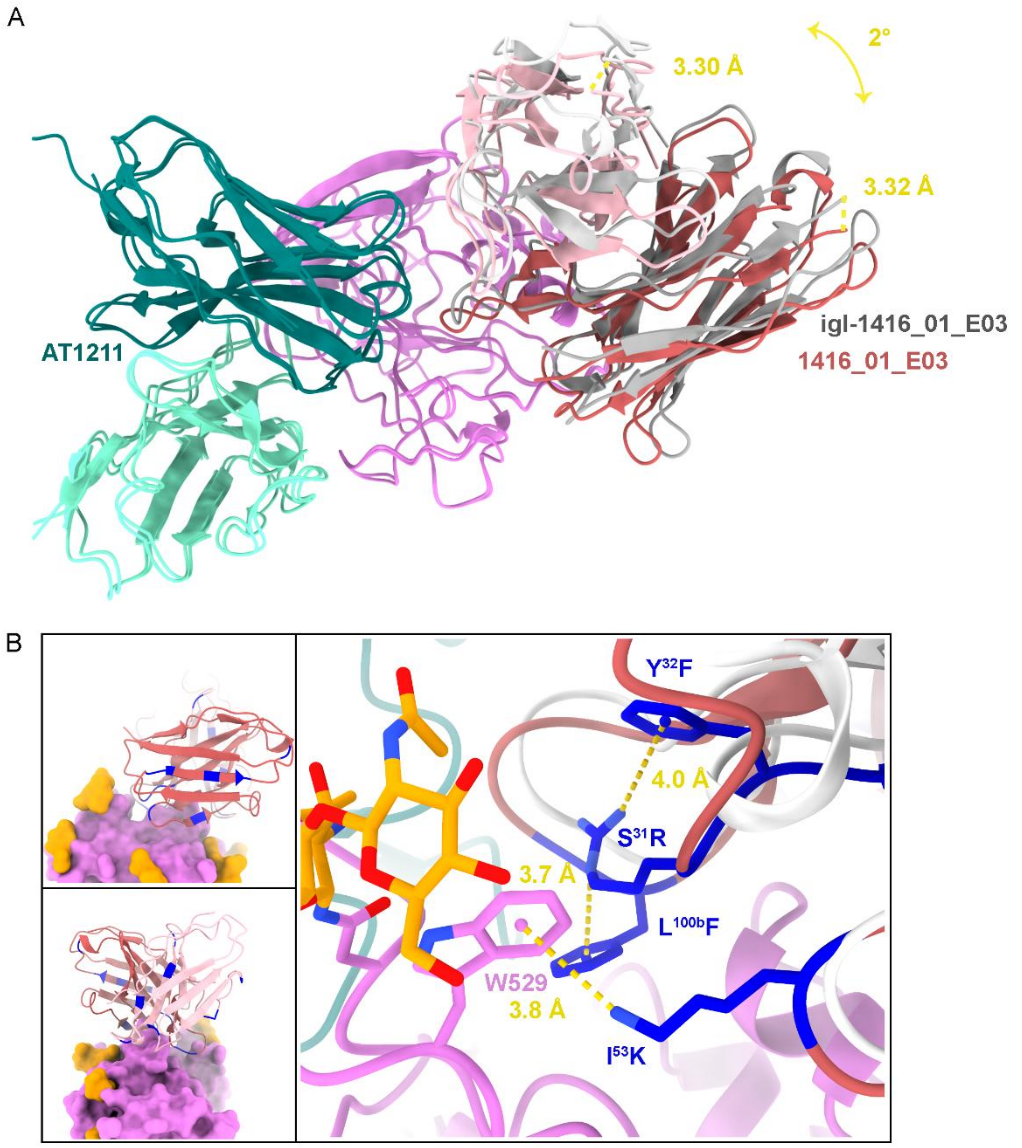


***Supplementary Figure 16:*** *Comparison of 1416_01_E03 and igl-1416_01_E03 binding to E2 in complex with AT1211.* ***A*** *Models for each complex were aligned on E2 molecule, and color coded as before, with the exception of igl-1416_01_E03 colored grey. Distances indicate the displacement between the most C-terminal residue on heavy and light chains between the germline and mature bNAb, while the angle represents the angle between their center of mass and AR3 plane's center of mass.* ***B*** *Somatically mutated residues on 1416_01_E03 are highlighted in blue. Residues interacting with E2 are shown as sticks, with heteroatoms (blue: nitrogen, red: oxygen).*

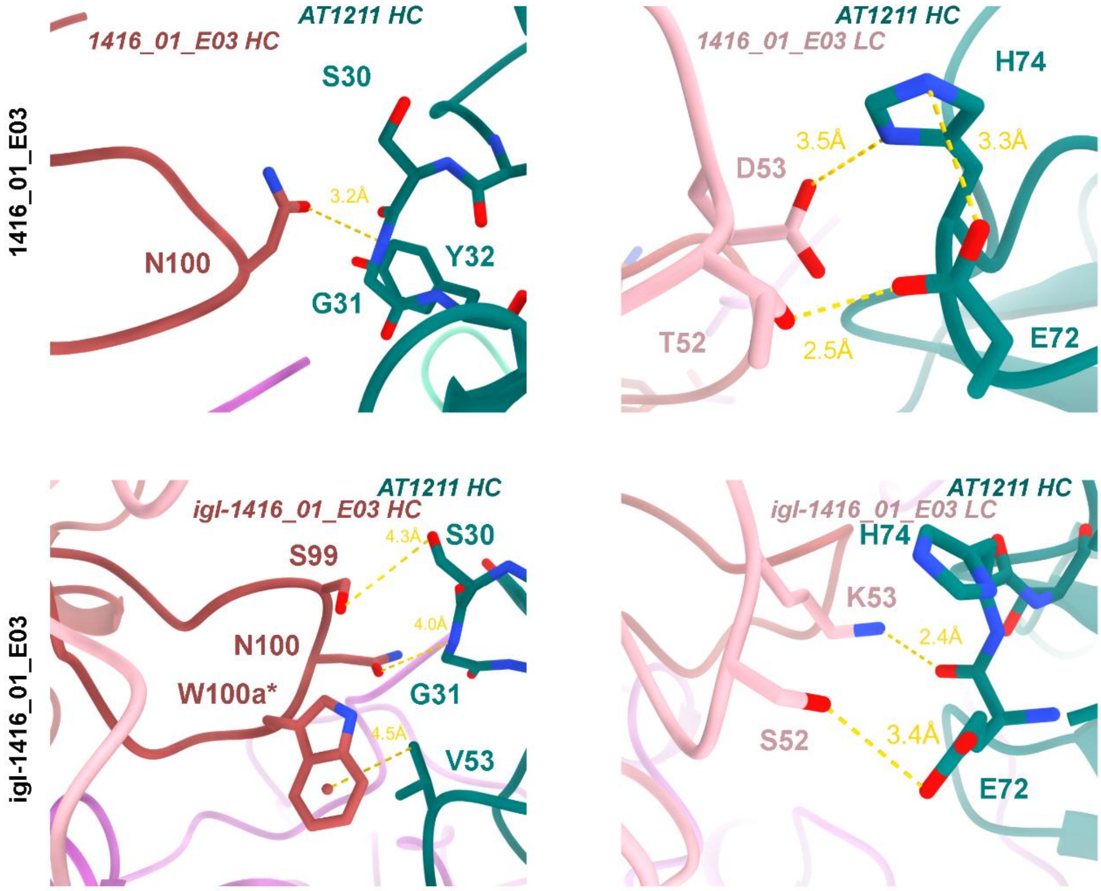


***Supplementary Figure 17: Comparison of 1416_01_E03 and igl-1416_01_E03 binding to E2 in complex with AT1211.*** *Heterotypic contacts between AT1211 heavy chain (teal) and 1416_01_E03 or igl-1416_01_E03 heavy and light chain (red and pink, respectively). Star above a residue label indicates position at which the residue is somatically mutated in the mature 1416_01_E03 bNAb.*
